# Somatodendritic HCN channels in hippocampal OLM cells revealed by a convergence of computational models and experiments

**DOI:** 10.1101/633941

**Authors:** Vladislav Sekulić, Feng Yi, Tavita Garrett, Alexandre Guet-McCreight, Yvette Y. Lopez, Mychael Solis-Wheeler, Rui Wang, Xiaobo Liu, J.Josh Lawrence, Frances K. Skinner

**Affiliations:** Krembil Research Institute, University Health Network, Toronto, Ontario, Canada; Department of Physiology, University of Toronto, Toronto, Ontario, Canada; Department of Biomedical and Pharmaceutical Sciences, Center for Structural and Functional Neuroscience, and Center for Biomolecular Structure and Dynamics, University of Montana, Missoula, Montana, USA; Neuroscience Graduate Program and Oregon Hearing Research Center and Vollum Institute; Oregon Health and Science University, Portland, Oregon, USA; Department of Pharmacology and Neuroscience, Texas Tech University Health Sciences Center, Lubbock, Texas, USA; Honors College, Texas Tech University, Lubbock, Texas, USA; Graduate School of Biomedical Sciences, TTUHSC, Lubbock, TX USA; Center of Excellence for Translational Neuroscience and Therapeutics, Texas Tech University Health Sciences Center, Lubbock, Texas, USA; Departments of Medicine (Neurology) and Physiology, University of Toronto, Toronto, Ontario, Canada

## Abstract

Determining details of spatially extended neurons is a challenge that needs to be overcome. The oriens-lacunosum/moleculare (OLM) interneuron has been implicated as a critical controller of hippocampal memory making it essential to understand how its biophysical properties contribute to function. We previously used computational models to show that OLM cells exhibit theta spiking resonance frequencies that depend on their dendrites having hyperpolarization-activated cation channels (h-channels). However, whether OLM cells have dendritic h-channels is unknown. We performed a set of whole-cell recordings of OLM cells from mouse hippocampus and constructed multi-compartment models using morphological and electrophysiological parameters extracted from the same cell. The models matched experiments only when dendritic h-channels were present. Immunohistochemical localization of the HCN2 subunit confirmed dendritic expression. These models can be used to obtain insight into hippocampal function. Our work shows that a tight integration of model and experiment tackles the challenge of characterizing spatially extended neurons.

## Introduction

The challenge of understanding brain function given its many cell types and circuits is being greatly aided by the development of sophisticated experimental techniques, big data, and interdisciplinary collaborations (*Ecker et al., 2017*). Furthermore, the use of computational brain models is becoming more established as an important tool that can bridge across scales and levels (*Bassett et al., 2018*; *Cutsuridis et al., 2010*; *O’Leary et al., 2015*). It is becoming increasingly clear that it is essential to consider the unique contributions of specific cell types and circuits in order to understand brain behaviour (*Luo et al., 2018*). In particular, we know that different inhibitory cell types can control circuit output and brain function in specific ways (*Abbas et al., 2018*; *Cardin, 2018*; *Kepecs and Fishell, 2014*; *Roux and Buzsáki, 2015*) and, by extension, disease states (*Marín, 2012*).

The contribution of a specific cell type to network and behavioural function is necessarily grounded in its biophysical properties. While immunohistochemical and single-cell transcriptomic studies provide insight into which ion channels might be present in a particular cell type, how different cell types contribute to function must necessarily include its activity within circuits (*Kopell et al., 2014*). An individual neuron’s activity largely arises from its ion channel kinetics, densities, and localization across its neuronal compartments. In this regard, mathematical multi-scale (channel and cellular), multi-compartment computational models are needed to help provide insights and hypotheses of how specific cell types contribute to brain function and disease processes. However, developing such mathematical models come with their own set of caveats and limitations. Creating such models requires quantitative knowledge of the precise characteristics of the particular cell type, and it is highly challenging, if not impossible, to obtain comprehensive knowledge of all the relevant biophysical parameters of each compartment of each cell type experimentally. Conversely, mathematical models, no matter how detailed, are always a simplification relative to biological reality and limited by the available experimental data. It is therefore important not to lose sight of the limitations of both model and experiment by having an ongoing dialogue between the two.

It is now well-known that the characteristics of a given cell type are not fixed (Marder and Goaillard, 2006), and thus a component of experimental variability reflects heterogeneity inherent in specific neuronal populations and thus in circuits. Moreover, such variability is likely to be functionally important (*Wilson, 2010*). Previous work has shown that conductance densities for a given ion channel in an identified cell type can have a two to six-fold range of values (*Goaillard et al., 2009*; *Ransdell et al., 2013*). Despite this variability in channel conductances, robust neuronal as well as circuit output is maintained, as most clearly shown in the crustacean stomatogastric ganglion circuit (*Bucher et al., 2005*; *Schulz et al., 2006*; *Tang et al., 2012*). The conservation of individual neuronal electrical output despite variable underlying ion channel conductance densities has furthermore been demonstrated in mammalian CNS neurons (*Swensen and Bean, 2005*), most likely arising from complex homeostatic mechanisms for maintaining circuit stability that are not fully understood. Therefore, how should one proceed in building cellular, computational models? Averaging of experimental variables such as conductance densities as a way of accounting for variability leads to erroneous conductance-based models (*Golowasch et al., 2002*). As a consequence, single, “canonical” biophysical models cannot capture inherent variability in the experimental ion channel data. A more desirable approach is to develop multiple models to capture the underlying biological variability (*Marder and Taylor, 2011*). Indeed, such populations of models representing a given cell type have been developed to examine, for example, co-regulations between different conductances that might exist in a given cell type (*Hay et al., 2011*; *Sekulić et al., 2014*; *Soofi et al., 2012*). In this way, populations of models could help suggest what balance of conductances are important for cellular dynamics and their function in circuits. Ideally, one should obtain biophysical properties of a given cell type using recordings from the *same* cell. It is of course unrealistic to consider an experimental characterization of all the various ion channel types using the same cell of a given cell type. This impracticality is further enhanced in consideration of channel types in the dendrites of neurons. Besides needing to patch from the same cell, there is also the practical limitations of invasively investigating the biophysical characteristics of fine dendritic compartments. However, dendrites are where most synaptic contacts are made and where signal integration in neurons occurs (*Stuart and Spruston, 2015*). Thus, these aspects must be tackled along with considerations of cellular variability.

In this work we focus on the oriens-lacunosum/moleculare (OLM) cell, an identified inhibitory cell type in the hippocampal CA1 area (*Maccaferri and Lacaille, 2003*; *Müller and Remy, 2014*). OLM cells receive excitatory glutamatergic input predominantly from local CA1 pyramidal neurons and form GABAergic synapses onto the distal dendrites of CA1 pyramidal neurons, as well as onto other CA1 inhibitory cells (*Blasco-Ibáñez and Freund, 1995*; *Klausberger, 2009*; *Leão et al., 2012*; *Maccaferri et al., 2000*). Functionally, proposed roles of OLM cells include gating sensory and contextual information in CA1 (*Leão et al., 2012*), and supporting the acquisition of fear memories (*Lovett-Barron et al., 2014*). Moreover, OLM cell firing is phase-locked to the prominent theta rhythms in the hippocampus of behaving animals (*Katona et al., 2014*; *Klausberger et al., 2003*; *Klausberger and Somogyi, 2008*; *Varga et al., 2012*). Although it has long been known that OLM cells express hyperpolarization-activated cation channels (h-channels) (*Maccaferri and McBain, 1996*), it is still unclear whether these channels are present in their dendrites. From a functional perspective, the consequences of dendritic h-channel expression in OLM cells was explored in our previous computational study where h-channels were found to modulate the spiking preference of OLM cell models – incoming inhibitory inputs recruited either a higher or lower theta frequency (akin to Type 1 or Type 2 theta, respectively - *Kramis et al. (1975)*) depending on the presence or absence of dendritic h-channels (*Sekulić and Skinner, 2017*). In that computational study, our OLM cell models were derived from previously built populations of OLM cell multi-compartment models in which appropriate OLM cell models were found with h-channels present either in the soma only or uniformly distributed in the soma and dendrites (*Sekulić et al., 2014*). We had previously leveraged these models and showed that appropriate OLM cell model output could be maintained, even if h-channel conductance densities and distributions co-vary, so long as total membrane conductance due to h-channels is conserved (*Sekulić et al., 2015*) – a finding that was also demonstrated in cerebellar Purkinje neurons (*Angelo et al., 2007*). Moreover, these OLM cell models were developed using morphological and electrophysiological data obtained from different OLM cells as well as h-channel characteristics from the literature, resulting in non-uniqueness of the fitted model parameters (*Holmes et al., 2006*; *Rall et al., 1992*). Prior to the work here, we did not actually know whether h-channels were present in the dendrites of OLM cells. Dendritic patch-clamp experiments have not yet been performed on OLM cells to determine their existence, and immunohistochemistry studies had so far demonstrated expression of the HCN2 subunit of h-channels only in the somata of OLM cells (*Matt et al., 2011*).

Considering all of the above, we aimed to build “next generation” multi-compartment models of OLM cells to achieve a two-pronged goal. First, we wanted to determine whether multi-compartment models built using morphological and electrophysiological data from the *same* cell would produce *consistent* results regarding h-channel localization to dendrites or not, and second, to determine the biophysical characteristics of h-channels in OLM cells. We considered our models to be next generation over previous multi-compartment OLM cell modelling efforts because each model was built using experimental data from the same cell, including its morphology, passive properties, and biophysical h-channel characteristics. Using transgenic mice in which yellow fluorescent protein (YFP) was expressed in somatostatin (SOM)-containing neurons, we visually targeted OLM cells from CA1 hippocampus, and fully reconstructed three OLM cells for multi-compartment model development with h-channel characteristics fit to each particular cell. We found that in order to be compatible with the experimental data, all three models needed to have h-channels present in their dendrites. Further, we performed immunohistochemical experiments that supported this prediction of dendritic h-channel expression in OLM cells. Finally, using two of these reconstructed models, we completed their development into full spiking models by including additional ion channel currents whose parameters were optimized based on voltage recordings from the same cell. These resulting models and associated experimental data are publicly available and can be subsequently used to develop further insight into the biophysical specifics of OLM cells and to help understand their contributions to circuit dynamics and behaviour. This work also demonstrates the feasibility of combining experimental and computational studies to address the challenging issue of determining the density and distribution of specific dendritic ion channel types.

## Results

### YFP-positive stratum oriens interneurons from SOM-Cre/Rosa26YFP mice contain a population of oriens-lacunosum/moleculare (OLM) cells

Patch-clamp recordings from 45 YFP-positive neurons in the stratum oriens of SOM-Cre/Rosa26YFP mice were obtained. Of these recordings, 11 of them met criteria for stability (see Methods) and were morphologically confirmed as OLM interneurons, having horizontal cell body and dendrites confined to the oriens layer, and perpendicularly projecting axons which ramify in the lacunosum/moleculare layer. Morphological and electrophysiological characteristics for three of these OLM cells are shown in *Figure 1*. An analysis of the 11 cells is given in *Table 1*. YFP-positive stratum oriens interneurons from SOM-CRE/Rosa26YFP mice had slow membrane time constants, and high input resistances, in accordance with our previous study (*Yi et al., 2014*). Action potential half-widths were larger and membrane time constants were slower than previously reported for YFP neurons from PV-CRE/Rosa26YFP mice, consistent with the exclusion of PV-positive basket and bistratified cells from this population. Moreover, this population had considerable hyperpolarization-induced sag, which, when combined with their higher input resistance, is considered a hallmark feature of OLM cells (*Maccaferri and Lacaille, 2003*).

**Table 1.** Passive and active properties of OLM cells from SOM-Cre/Rosa26YFP mice.

| Property | SOM-YFP (n=11) |
| --- | --- |
| $R_{in}$ (MΩ) | 314.3 ± 33.8 |
| $C_m$ (pF) | 107.1 ± 13.0 |
| Sag ratio (SS/peak) | 0.89 ± 0.03 |
| $\tau_m$ (ms) | 31.1 ± 2.5 |
| $I_{hold}$ (pA) | 3.4 ± 4.8 |
| First AP half-width (μs) | 595.1 ± 26.4 |
| First AP height (mV) | 66.0 ± 2.7 |
| Adaptation coefficient | 0.5 ± 0.1 |
| Frequency at 90 pA (Hz) | 15.5 ± 2.1 |
| Frequency at 60 pA (Hz) | 8.3 ± 1.5 |
| Frequency at 30 pA (Hz) | 2.1 ± 0.8 |

Values are presented as means ± SEM. Abbreviations: $R_{in}$ , input resistance; $\tau_m$ , membrane time constant; AP, action potential; $C_m$ , membrane capacitance; SS, steady state; $I_{hold}$ , the current injected to hold the recorded cell at -60 mV (not junction potential corrected). Passive properties were extracted at a current injection of -60 pA. Active properties were extracted at a current injection of 90 pA, unless otherwise specified. APs were detected at a derivative threshold of 20 mV/ms. The adaptation coefficient was calculated by dividing the first inter-spike interval by the last inter-spike interval of the AP train during current injection of 90 pA.

### Multi-compartment model fits to the charging portion of the membrane potential describe passive membrane responses

From the 11 usable cells (*Table 1*), three cells were selected for multi-compartment model development (***Cell 1, Cell 2, Cell 3***). We used the NEURON simulation environment (*Hines and Carnevale, 2001*) to develop our multi-compartment models. *Figure 1A,B* shows imaging of the three chosen cells, with the reconstructed cell morphologies shown in *Figure 1C*, and typical electrophysiological OLM cell profiles with sag characteristics shown in *Figure 1D*. Details of the model reconstructions are given in the Methods.

**Figure 1.**
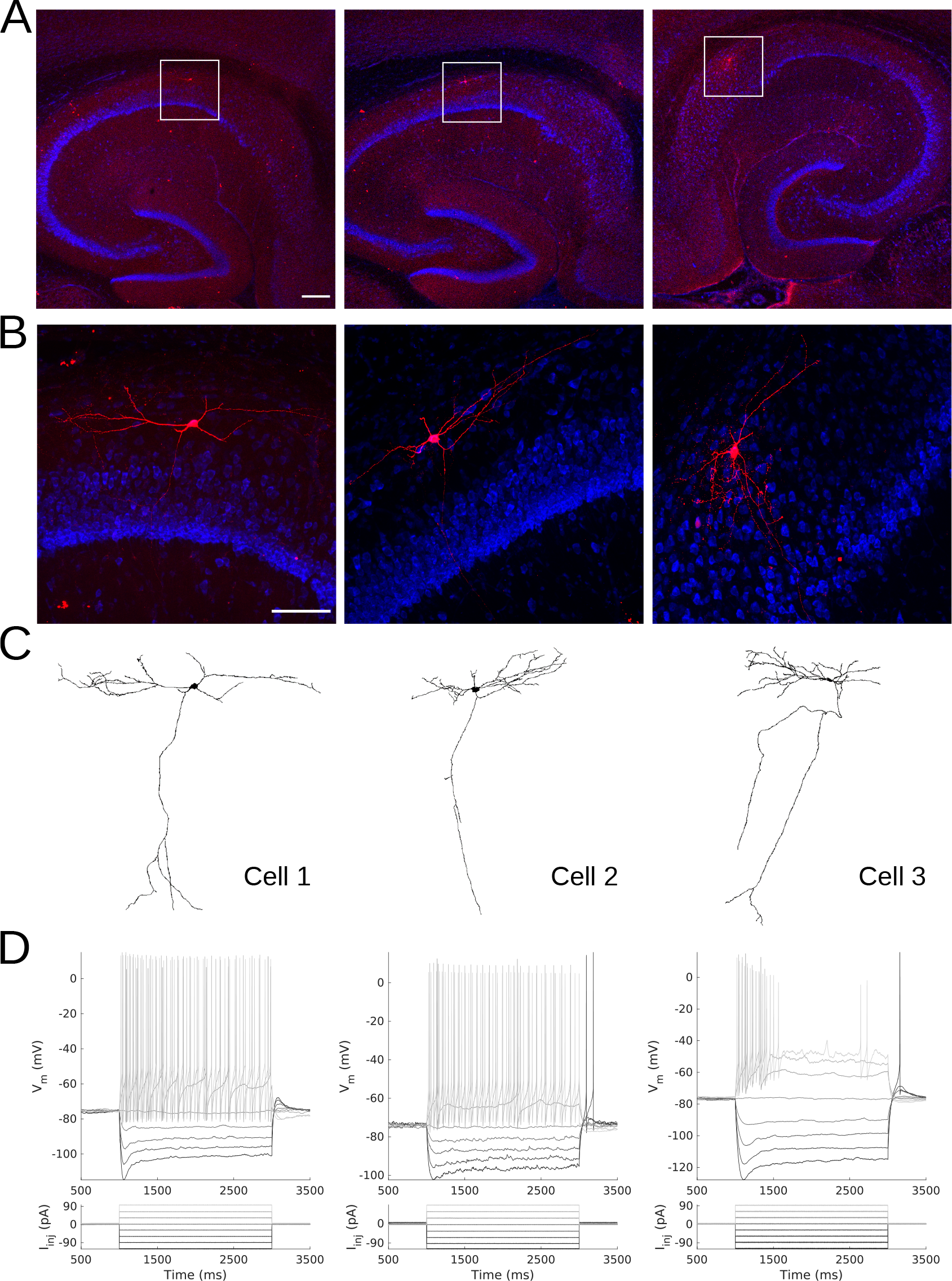
Morphology and Electrophysiology of OLM interneurons. Confocal images of OLM cells at **A.** 4X and **B.** 25X magnification, corresponding to *Cell 1, Cell 2,* and *Cell 3* (respectively, left, middle, and right - ordering of cells are the same for remainder of figure). Squares in **A.** indicate boundaries of zoomed images in **B.** Scale bars: 200 *μ* and 100 *μ* for **A.** and **B.**, respectively. **C.** Reconstructed morphologies for the three cells. **D.** Membrane potential (*V*_*m*_) traces in response to current clamp injections in the presence of synaptic blockers showing intrinsic electrophysiological properties.

**Figure 2.**
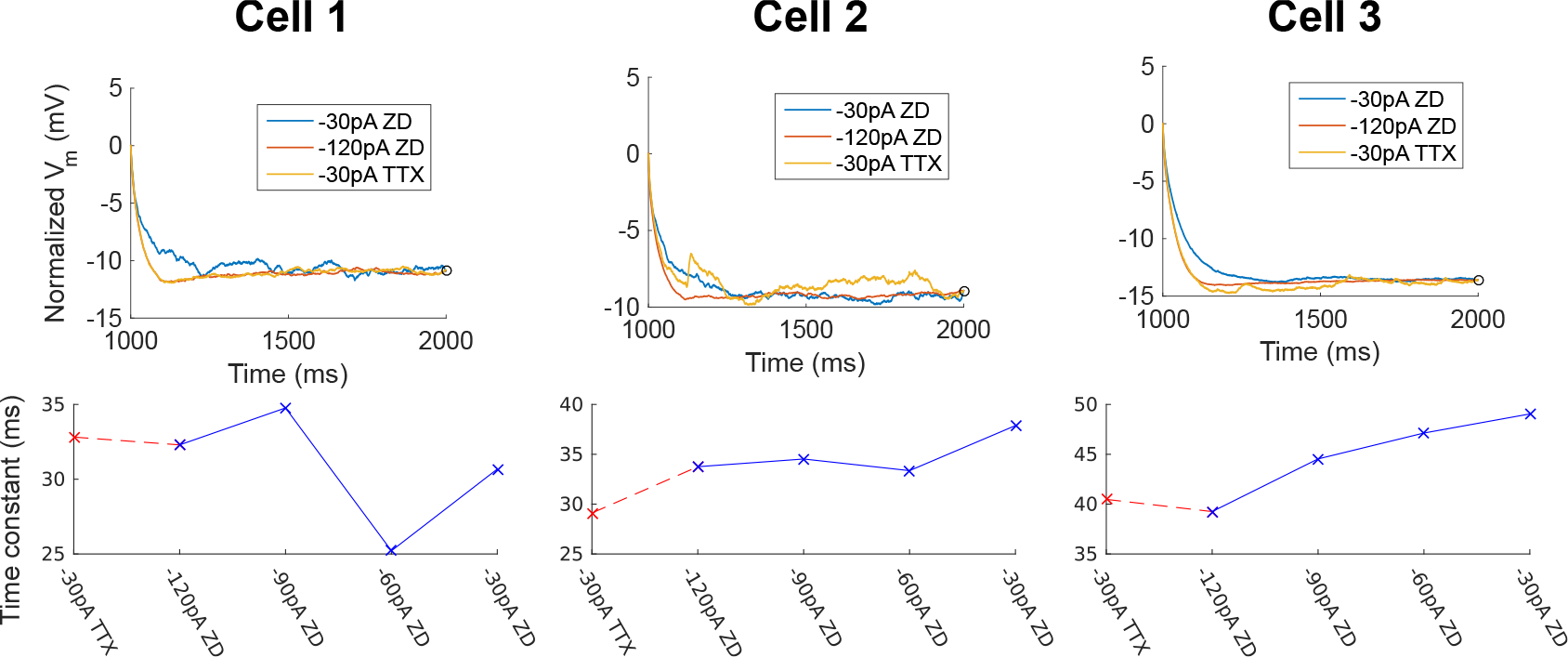
Passive Properties, Membrane Time Constants. (Top) Membrane potential (*V*_*m*_) normalized at steady-state, showing noisier responses of −30pA ZD trace (from protocol #7 in *Table 5*) compared to the −120pA ZD trace. (Bottom) Fitted membrane time constants (*τ*_*m*_) for all current clamp steps with ZD7288 application, as well as the −30pA TTX trace (from protocol #4 in *Table 5*), i.e.,current clamp step without ZD7288 application, but with other channel blockers present: TTX/4-AP/TEA.

To capture the passive response of the three cells we used long −120 pA current clamp traces in which all synaptic and voltage-gated channels were blocked. This choice was made because we found that the −30 pA traces were noisier in general (see *Figure 2*, top panels), and the −120 pA traces best captured the passive response of the cells. This can be seen from a comparison of the membrane time constants (*τ*_*m*_) for different current clamp steps (see *Figure 2*, bottom panels) and consideration of the protocol ordering of the recording session. Full details are provided in the Methods. The resulting fitted passive parameters of axial resistivity (*R*_*a*_), specific capacitance (*C*_*m*_), leak conductance (*G*_*pas*_) and leak reversal potential (*E*_*pas*_) (see *Table 6* in Methods, top of *Table 2* and as summarized in *Table 4* later) in conjunction with the respective cell morphologies form the “backbone” of the OLM cell models.

### Low specific capacitances are a robust feature of OLM cells

From our model fits we found that the *C*_*m*_’s obtained were much lower than the ≈ 0.9-1*μ*F/cm^2^ that have been previously reported as a “standard” value in mammalian neurons (*Gentet et al., 2000*). These values indicated the possibility of errors in the process of reconstructing the cell morphologies. In particular, we investigated our estimations of dendritic diameter, which is a particularly prominent source of possible errors in morphological reconstruction of neurons (*Jaeger, 2001*). For instance, although fine dendritic processes with diameters between 0.5 and 2*μ*m can be resolved using confocal microscopy, their apparent diameters will generally seem larger than their true size due to the point spread function of the optical system (*Jacobs et al., 2010*). To assess whether our low fitted *C*_*m*_ values re2ected compensation for overestimated dendritic diameters or whether they may reflect lower specific capacitance in biological OLM cells, we turned to cable theoretic considerations.

*Holmes et al. (2006)* showed that when either the diameter *d* or length *l* are multiplied by a constant factor *x*, the derived changes in membrane resistivity (*R*_*m*_), *R*_*a*_, and *C*_*m*_ needed to maintain an identical voltage response occur when *R*_*m*_ is scaled by *x*, *C*_*m*_ by 1/*x*, and *R*_*a*_ by *x*^2^. Note that since *G*_*pas*_ corresponds to the inverse of *R*_*m*_, the necessary change for *G*_*pas*_ is rather to scale it by 1/*x*. If we consider re-scaled dendritic diameters of our OLM models by scaling values (*d*_*s*_) of 0.5 and 1.5 to represent large changes, and 0.9 and 1.1 to represent small changes, the resulting values of *G*_*pas*_, *C*_*m*_, and *R*_*a*_ as predicted by cable theory are shown in *Table 2*. To confirm these predictions, we first re-scaled the original reconstructed diameters for each compartment in two of the models (*Cell 1* and *Cell 2*) using these four scaling values (0.5, 1.5, 0.9, 1.1). We used our unchanged passive properties previously obtained and examined the responses to a −120pA current clamp step. This served to assess the resulting changes in the *V*_*m*_ responses attributable to errors in electrotonic properties solely due to changing the diameters. We found that changes in diameters produced deviations in the *V*_*m*_ response roughly proportional to the magnitude of the scaled dendritic diameters (*Figure 2-Figure Supplement 2A*). The largest change was when the diameters were halved across all compartments (*d*_*s*_ = 0.5). We then re-fitted the passive properties of the two models under each case of re-scaled dendritic diameters, obtaining nearly identical model *V*_*m*_ responses to the −120pA current clamp step (*Figure 2-Figure Supplement 2B*). We found that the refitted values of *R*_*m*_, *C*_*m*_, and *R*_*a*_ for the various cases of scaled dendritic diameters, *d*_*s*_, were in excellent agreement with the values predicted by cable theory (*Table 2*). Thus, we were able to use the cable theoretic predictions to provide implicit limits on how much the fitted parameters could be expected to deviate solely due to errors in morphological considerations.

**Table 2.** Re-fitted passive properties for models with scaled dendritic diameters.

|  |  | **fitted**<br>**values** |  |  |  |  | **cable**<br>**theory** |  |  |
| --- | --- | --- | --- | --- | --- | --- | --- | --- | --- |
| | $d_s$ | RMSE<br>(mV) | $R_a$<br>(Ωcm) | $C_m$<br>(μF/cm<sup>2</sup>) | $G_{pas}$<br>(S/cm<sup>2</sup>) | $E_{pas}$<br>(mV) | $R_a^*$<br>(Ωcm) | $C_m^*$<br>(μF/cm<sup>2</sup>) | $G_{pas}^*$<br>(S/cm<sup>2</sup>) |
| **Cell 1** | **1.0** | 0.3602 | 141.85 | 0.2698 | $7.933 \times 10^{-6}$ | 49.05 | | | |
| | 0.5 | 0.3665 | 36.87 | 0.5278 | $1.556 \times 10^{-5}$ | 49.09 | 35.46 | 0.5396 | $1.586 \times 10^{-5}$ |
| | 1.5 | 0.3959 | 386.51 | 0.1592 | $4.768 \times 10^{-6}$ | 49.28 | 319.16 | 0.1799 | $5.288 \times 10^{-6}$ |
| | 0.9 | 0.3766 | 121.35 | 0.2802 | $8.342 \times 10^{-6}$ | 49.18 | 114.89 | 0.2998 | $8.814 \times 10^{-6}$ |
| | 1.1 | 0.3827 | 195.82 | 0.2269 | $6.750 \times 10^{-6}$ | 49.20 | 171.63 | 0.2453 | $7.211 \times 10^{-6}$ |
| **Cell 2** | **1.0** | 0.9818 | 285.78 | 0.2799 | $9.242 \times 10^{-6}$ | 54.74 | | | |
| | 0.5 | 0.9104 | 64.78 | 0.6484 | $2.085 \times 10^{-5}$ | 54.60 | 72.44 | 0.5598 | $1.848 \times 10^{-5}$ |
| | 1.5 | 1.0486 | 707.21 | 0.1624 | $5.480 \times 10^{-6}$ | 54.85 | 643.01 | 0.1866 | $6.161 \times 10^{-6}$ |
| | 0.9 | 0.9676 | 226.97 | 0.3201 | $1.051 \times 10^{-5}$ | 54.71 | 231.48 | 0.3110 | $1.029 \times 10^{-5}$ |
| | 1.1 | 0.9958 | 350.50 | 0.2470 | $8.198 \times 10^{-6}$ | 54.77 | 345.79 | 0.2545 | $8.401 \times 10^{-6}$ |

$d_s$ is the scaled diameter. $R_a^*$ , $C_m^*$ and $G_{pas}^*$ are the values as predicted from cable theory and are given by

All of our models were found to have consistent but low *C*_*m*_ values as shown in *Table 6* (0.27 for *Cell 1*, 0.28 for *Cell 2*, and 0.31 for *Cell 3*, units of *μ*F/cm^2^). Let us suppose that the diameter estimates in our morphological reconstructions were nevertheless overestimates of the true diameters. Then, the cable theoretic predictions, as confirmed by our simulations, would suggest that even if we halved the average diameters, the expected *C*_*m*_ in the cells would only increase to about 0.58*μ*F/cm^2^, taking the average of the re-fitted *C*_*m*_ of *Cell 1* and *Cell 2*. We considered it unlikely that our diameters were this much in error. That is, a scaling factor of 0.5 implies that the minimum diameter, which is 0.35 *μ*m in *Cell 1* and 0.32 *μ*m in *Cell 2*, would had to have been reduced to 0.17 *μ*m and 0.16 *μ*m for *Cell 1* and *Cell 2*, respectively, which are unreasonably small. If we nevertheless consider these scaled diameter values as possible ranges, we are led to the consideration that *C*_*m*_ in OLM cells may be within the range of 0.2-0.6*μ*F/cm^2^. We also note that if we compute *C* for these three cells directly from experimental capacitance and surface area values, then they are also low, as shown in *Table 3*. Due to these analyses, we decided to keep the values of *C*_*m*_ obtained from the passive property fitting procedure for each model as we proceeded to build upon this passive backbone.

**Table 3.** Specific capacitance computed directly from experiment.

| *Parameter* | *Cell 1* | *Cell 2* | *Cell 3* |
| --- | --- | --- | --- |
| Capacitance (pF) | 62.8 | 123.7 | 79.6 |
| Surface area ($\mu\text{m}^2$) | 29,378.1 | 35,158.5 | 21,990.3 |
| $C_m$ ($\mu\text{F}/\text{cm}^2$) | 0.21 | 0.35 | 0.36 |

*Capacitance properties were extracted at a current injection of -60 pA.*

### H-channels inserted into OLM cell models with constrained passive properties do not yield ideal fits

Using h-channel current (*I*_*h*_) parameter values estimated from the experimental data (see Methods), we inserted h-channels into our OLM cell multi-compartment models’ ‘backbone’ of morphological reconstructions and passive property fits. The h-channel parameters extracted from each of the OLM cells in this work included the *I*_*h*_ reversal potential (*E*_*h*_), the time constants of activation and deactivation of *I*_*h*_ (*τ*_*h*_), the steady-state activation curve of *I*_*h*_ (*r*_∞_), and the *I*_*h*_ maximum conductance density (*G*_*h*_) (*Figure 3*).

We demonstrated in our previous work that OLM cell models exhibited a tradeoff between total membrane *G*_*h*_ and the dendritic distribution of h-channels so that if the total *G*_*h*_ was conserved, the resulting model output would be appropriate (*Sekulić et al., 2015*). Now, for the first time, we have a measure of total *G*_*h*_. Thus, a key prediction for the resulting multi-compartment models is that the total *G*_*h*_ will constrain the distribution of h-channels to allow the models to appropriately capture the OLM cell electrophysiological characteristics. To consider this, we added an additional parameter to our models termed *H*_*dist*_, which is defined as the centripetal extent for which h-channels are inserted in the dendrites. It is defined by a real-valued number in the range of [0, 1] and represents the fraction of maximum dendritic path length from the soma on a per-cell basis. Compartments with a path length from the soma that was smaller than any given *H*_*dist*_ value were included when subsequently inserting h-channels, whereas those compartments whose distance from soma exceeded *H*_*dist*_ were excluded. The boundary condition of *H*_*dist*_ = 0 is defined as the case where h-channels are only present in somatic compartments and not present in the dendrites. A non-zero value for *H*_*dist*_ meant that the amount of dendrite specified by *H*_*dist*_ itself had h-channels in addition to the somatic compartments. *H*_*dist*_ = 1 refers to full somatodendritic presence of h-channels, i.e., uniform distribution in the dendrites and soma. The per-cell *I*_*h*_ parameters were inserted into each of the three models and two cases of *H*_*dist*_ were initially considered to test the boundary cases: either no dendritic h-channels (*H*_*dist*_ = 0) or full, uniform distribution of h-channels in the dendrites (*H*_*dist*_ = 1). The resulting h-channel conductance density was calculated by dividing total *G*_*h*_ by the resulting surface area of only somatic or somatodendritic compartments. These values and the other h-channel parameters are given in *Table 8* in the Methods. The *V*_*m*_ output of each of the three models being developed here, with boundary conditions of *H*_*dist*_ = 0 or 1 for the case of −120pA current clamp injection, compared to the experimental TTX trace for each cell, is shown in *Figure 4A*. These models do not fully match the experimental traces. Although we did explore *H*_*dist*_ values that were between 0 and 1 (not shown), it is clear that given the fits shown in *Figure 4A*, it is unlikely that changing *H*_*dist*_ to a value between 0 (somatic expression only) to 1 (full somato-dendritic expression) would improve the fits to the experimental data.

**Figure 3.**
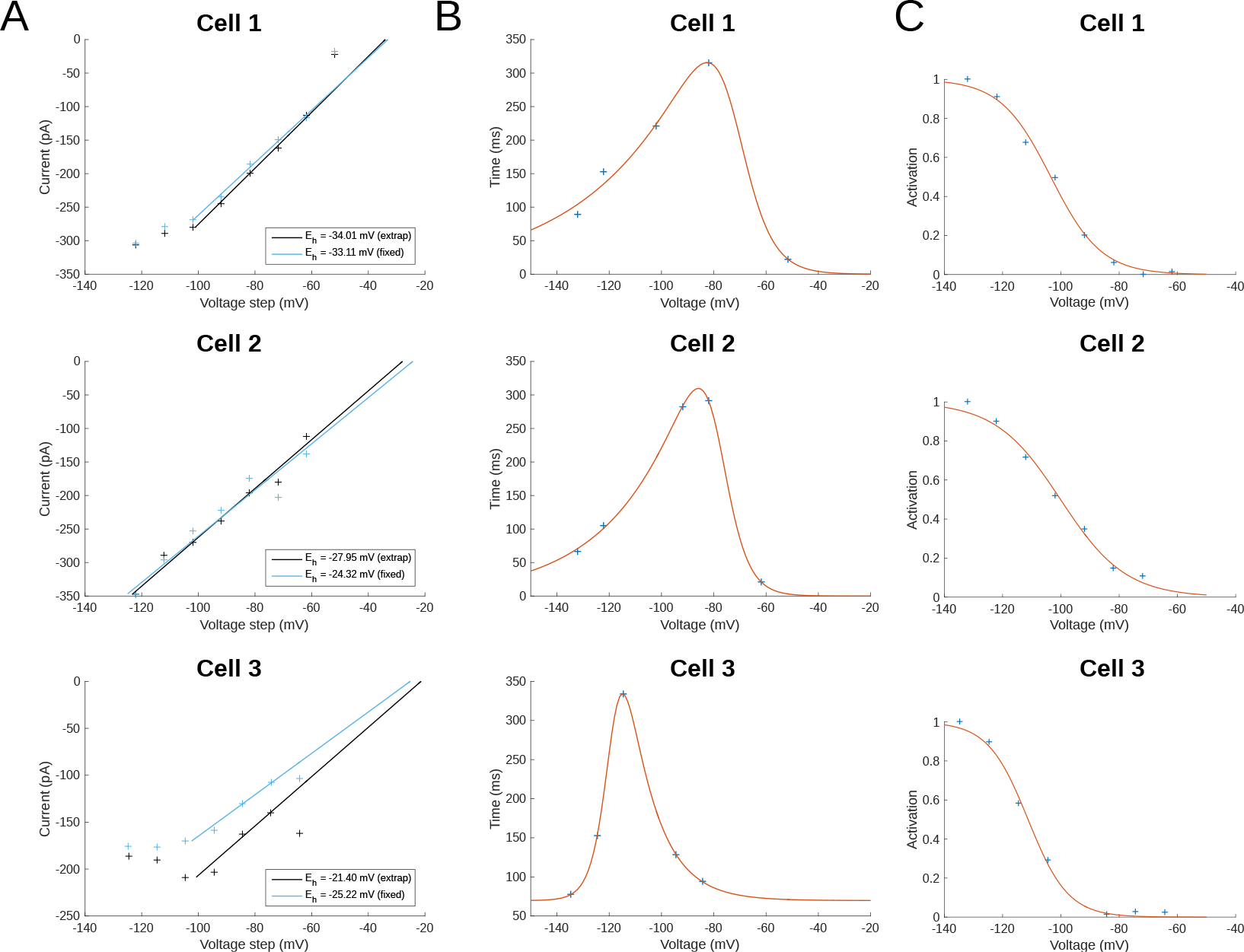
*I*_*h*_ Properties from Fitting to Experimental Data. **A.** *I*_*h*_ reversal potential as determined from current-voltage obtained from tail currents. See text in Methods for difference between ‘extrap’ and ‘fixed’ values in plots. **B.** Time constants (*τ*_*h*_) or kinetics of activation and deactivation. **C.** Steady-state activation curves (*r*_∞_).

**Figure 4.**
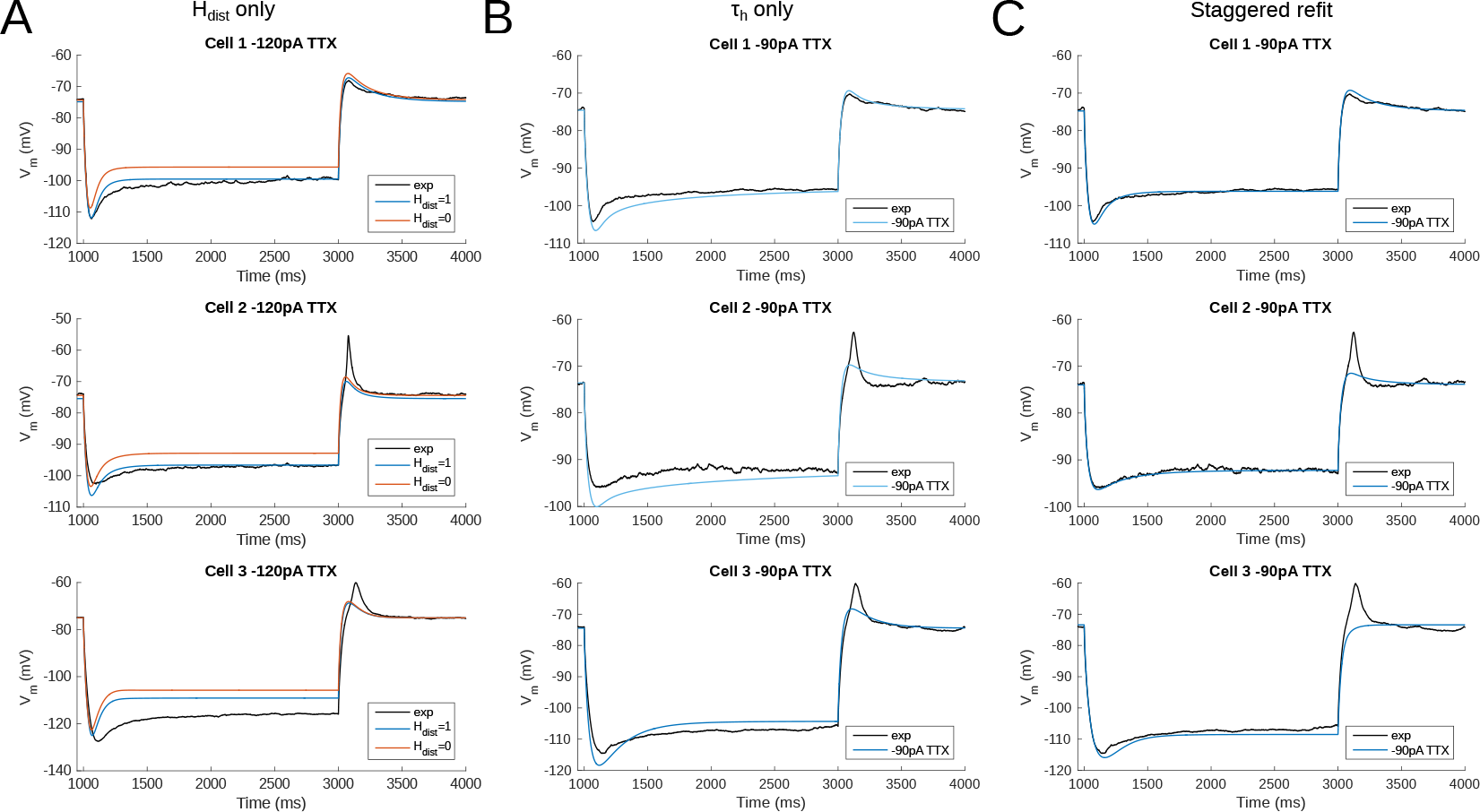
Fitting and Staggered Re-fitting of *I*_*h*_ and Passive Property Parameters Show That Only OLM Cell Models with Dendritic H-channels Reproduce Experimental Traces. **A.** Boundary conditions of *H*_*dist*_ parameter [0, 1] showing inappropriate fits when putting all experimentally derived parameters together. **B.** Re-fitting only *τ*_*h*_ does not provide good fits (*H*_*dist*_=1). **C.** Staggered re-fitting of parameters results in good fits. Shown in B and C are −90pA TTX traces which are “test” traces not used for fitting (−120pA TTX trace was used for fitting).

### A staggered re-fitting procedure yields consistent and generalized model fits for OLM cells with dendritic h-channels

Given the suboptimal match of our models with the experimental data, even with model parameters determined from experiment on a per-cell basis, we considered the possibility that one or more of the parameters were mismatched between the experimental cells and the parameter values derived from the recordings. We considered re-fitting the various parameters in the model to ensure that *I*_*h*_ and passive parameters resulted in correct output for each cell. However, due to the sheer number of parameters present in the model, care needed to be taken in how the parameters were adjusted as there are many interdependencies between the fitted parameters. For instance, when *I*_*h*_ is present, the trajectory of the *V*_*m*_ response upon a step of hyperpolarizing current in a cell depends not just on *C*_*m*_ and *R*_*a*_, but also on the time constants of activation and deactivation of h-channels (*τ*_*h*_) and, to a degree, the h-channel steady-state activation curve (*r*_∞_). Therefore, if there is error in the model *V*_*m*_ response compared to the experimental trace in this portion of the trace (*Figure 4A*), the mismatch between model and experiment may have been either due to the passive parameters, or due to *τ*_*h*_ or *G*_*h*_, which gated by the activation, determines the amount of *I*_*h*_. The problem, then, is how to attribute errors in any particular portion of a *V*_*m*_ trace to any given parameter in the model.

We noted that the initial mismatch in the case of *H*_*dist*_=1 and for *Cell 1* and *Cell 2* seem primarily to be located in the initial hyperpolarizing phase and the sag portion. Because the *τ*_*h*_ functions were constructed using a limited set of data, it was reasonable to suppose that a large source of mismatch in this portion of *V*_*m*_ could be due to errors in the *τ*_*h*_ function itself. We thus re-fitted the parameters for *τ*_*h*_, namely *t*_1_, *t*_2_, *t*_3_, *t*_4_, *t*_5_ for all three cells, against each respective −120pA trace and then compared the models’ responses to the other current clamp steps to see how much of the error could be accounted for by re-fitting *τ*_*h*_ alone (*Figure 4B*). This re-fitting of *τ*_*h*_ alone could not address the mismatch in *V*_*m*_ between model and experiment although it may have played some role, as evidenced by improving the match in *V*_*m*_ in some cases. Thus other parameter re-fitting needed to be considered. Detailed considerations including “overfitting” are given in the Methods.

We adopted the following approach and rationale. Since the passive properties were not as tightly constrained as the *I*_*h*_ properties, and could account for some of the mismatch in both the transient and steady state portions of the traces, we re-fitted them first. That is, *R*_*a*_, *C*_*m*_, *G*_*pas*_, *E*_*pas*_. Turning to the *I*_*h*_ properties, we first re-fitted the total *G*_*h*_, which determines the per-compartment conductance density, as well as the steady-state activation curve *r*_∞_ since it determines the voltage dependency of how many channels are open. We could not fit *G*_*h*_ and *r*_∞_ in the reverse order because any error in *G*_*h*_ – that is, how *I*_*h*_ scales with voltage when all channels are opened – could be accounted for by refitting *r*_∞_ by “flattening” it, thus lowering the total number of channels that are open at any given voltage. This would not be physiologically correct since the model would then imply that *I*_*h*_ is never fully activated, i.e., *r*_∞_ does not reach 1. Thus, by re-fitting *G*_*h*_ first, followed by *r*_∞_, we increased the likelihood that *r*_∞_ did not diverge too much from the experimental data points obtained from the protocol for *I*_*h*_ activation. Finally, we re-fitted *τ*_*h*_. If the passive properties and steady-state *I*_*h*_ due to *G*_*h*_ and *r*_∞_ accounted for much of the mismatch in *V*_*m*_, then the last step of re-fitting *τ*_*h*_ should allow for any mismatch due to *τ*_*h*_ to be corrected for.

Using this approach, which we termed a “staggered” re-fitting, we show the model outputs in *Figure 4C* where only the −120 pA TTX traces were used for fitting the parameters, with the −90pA TTX traces provided test data to validate the fits. We note that the results with this approach were more successful than the previous approaches. By fitting the parameters in such a way that the ones most likely to be responsible for errors in particular portions of the mismatched model *V*_*m*_ traces were fitted first, the resulting fits were more generalizable than the case of either re-fitting the passive properties alone, or re-fitting *τ*_*h*_ alone, since the staggered re-fitted values were able to match all of the other current clamp traces that were not used for fitting. This is shown in *Figure 4-Figure Supplement 2* for *Cell 1*, *Figure 4-Figure Supplement 3* for *Cell 2,* and *Figure 4-Figure Supplement 4* for *Cell 3*. As shown in *Figure 4B* and *Figure 4-Figure Supplement 1*, the other approaches clearly cannot capture other current clamp traces not used for the fitting. All of these staggered re-fitted values are shown in *Table 4*, *H*_*dist*_=1 column.

**Table 4.** Comparison of final fitted model parameters using either *H*_*dist*_=0 or *H*_*dist*_=1, and with model parameters before staggered re-fitting (*Table 6* and *Table 8*), starting fits, shown here for easy reference.

| **Parameter** | **Cell 1**<br>**Starting**<br>**Fits** | $H_{dist}=0$ | $H_{dist}=1$ | **Cell 2**<br>**Starting**<br>**Fits** | $H_{dist}=0$ | $H_{dist}=1$ | **Cell 3**<br>**Starting**<br>**Fits** | $H_{dist}=0$ | $H_{dist}=1$ |
| --- | --- | --- | --- | --- | --- | --- | --- | --- | --- |
| $R_a$ ($\Omega\text{cm}$) | 141.85 | 34.4 | 125.2 | 285.78 | 285.4 | 348.1 | 94.83 | 211.8 | 317.9 |
| $C_m$ ($\mu\text{F}/\text{cm}^2$) | 0.27 | 0.20 | 0.27 | 0.27 | 0.37 | 0.38 | 0.30 | 0.52 | 0.58 |
| $G_{pas}$ ($\text{S}/\text{cm}^2$) | $7.93 \times 10^{-6}$ | $7.36 \times 10^{-6}$ | $7.58 \times 10^{-6}$ | $9.24 \times 10^{-6}$ | $1.16 \times 10^{-5}$ | $1.19 \times 10^{-5}$ | $7.48 \times 10^{-6}$ | $8.26 \times 10^{-6}$ | $8.68 \times 10^{-6}$ |
| $E_{pas}$ (mV) | -49.0 | -64.0 | -64.6 | -54.7 | -61.5 | -61.8 | -69.1 | -75.7 | -76.1 |
| $E_h$ (mV) | -34.0 | -34.0 | -34.0 | -27.9 | -27.9 | -27.9 | -25.2 | -25.2 | -25.2 |
| $V_{1/2}$ (mV) | -103.4 | -103.1 | -103.7 | -100.1 | -100.6 | -99.6 | -111.3 | -108.4 | -113.8 |
| $k$ (mV) | 8.63 | 9.99 | 9.99 | 11.16 | 9.93 | 9.99 | 6.88 | 9.39 | 9.99 |
| $t_1$ (ms) | 8.03 | 12.30 | 8.56 | 8.98 | 12.73 | 11.28 | 35.09 | 36.77 | 41.84 |
| $t_2$ (ms) | 0.025 | 0.063 | 0.029 | 0.035 | 0.071 | 0.056 | 0.24 | 0.25 | 0.29 |
| $t_3$ (ms) | -4.40 | -22.87 | -6.91 | -8.49 | -20.38 | -19.28 | -4.28 | -3.22 | -4.03 |
| $t_4$ (ms) | 0.15 | 0.39 | 0.18 | 0.19 | 0.36 | 0.34 | 0.088 | 0.084 | 0.089 |
| $t_5$ (ms) | $7.32 \times 10^{-6}$ | 0.026 | $4.35 \times 10^{-5}$ | $3.57 \times 10^{-7}$ | $3.54 \times 10^{-6}$ | 0.006 | 69.72 | 5.69 | 4.29 |
| **Total $G_h$** | **4.17** | **1.91** | **3.12** | **3.64** | **0.75** | **2.14** | **2.20** | **0.77** | **1.82** |

All the models in the staggered re-fit were done with *H*_*dist*_=1, because that value was the one that provided the closest fit to the experimental traces (*Figure 4A*) when only passive properties were 1t to the *V*_*m*_ traces and *I*_*h*_ parameter values were obtained from the voltage clamp protocols. We examined whether using *H*_*dist*_=0 and applying our staggered re-fitting approach could also produce good, generalizable fits to the experimental data. The models with *H*_*dist*_=0 fitted the experimental *V*_*m*_ traces well in all four current clamp steps as it did for *H*_*dist*_=1, and we show the comparison to the −90pA TTX trace for *H*_*dist*_ values of both 0 and 1 in *Figure 5A*, noting that the −120 pA TTX trace was used for the fitting. The staggered re-fitted values for *H*_*dist*_=0 are also shown in *Table 4*. From a comparison across *Table 4* of parameter values for *H*_*dist*_= 0, 1 and original *I*_*h*_ parameter values 1t to the experimental data, it is clear that the re-fitted parameter values using *H*_*dist*_=0 are inappropriate. specifically, the total *G*_*h*_ (shown in bold in *Table 4*) for the case of *H*_*dist*_=1 was reasonably close to what was measured directly from the I-V plot of the reversal potential experimental protocol, unlike *H*_*dist*_=0, which exhibited *G*_*h*_ values that were much less than half of the experimentally-derived values. Given that this parameter was taken from the slope of the I-V plot, the values from *H*_*dist*_=0, if correct, would imply that the recorded current values were double the “true” values in the cell. This is graphically depicted in *Figure 5B*. We deemed this unlikely, and concluded that the relatively small divergence in the re-fitted *G*_*h*_ with *H*_*dist*_=1 compared to the experimental case indicated a much more reasonable error. Hence, the fact that it was possible to match the experimental *V*_*m*_ traces using both *H*_*dist*_=0 and *H*_*dist*_=1 did not mean that they were equally valid. The benefit of having directly measured experimental values representing *G*_*h*_, *τ*_*h*_, *r*_∞_ from the *same* cell meant that we could confidently state that models with *H*_*dist*_=0, though they fitted the *V*_*m*_ traces, were not appropriate models because they did not match the experimentally-derived values. Thus, only when h-channels were spread into the dendrites did we find models whose *V*_*m*_ responses matched the experimental traces and whose total *G*_*h*_ and other parameter values were in reasonable agreement with the experimentally measured values. We thus predict that the experimental cells in the dataset used here have h-channels expressed in their dendrites, with biophysical characteristics as given in *Table 4*, *H*_*dist*_=1.

**Figure 5.**
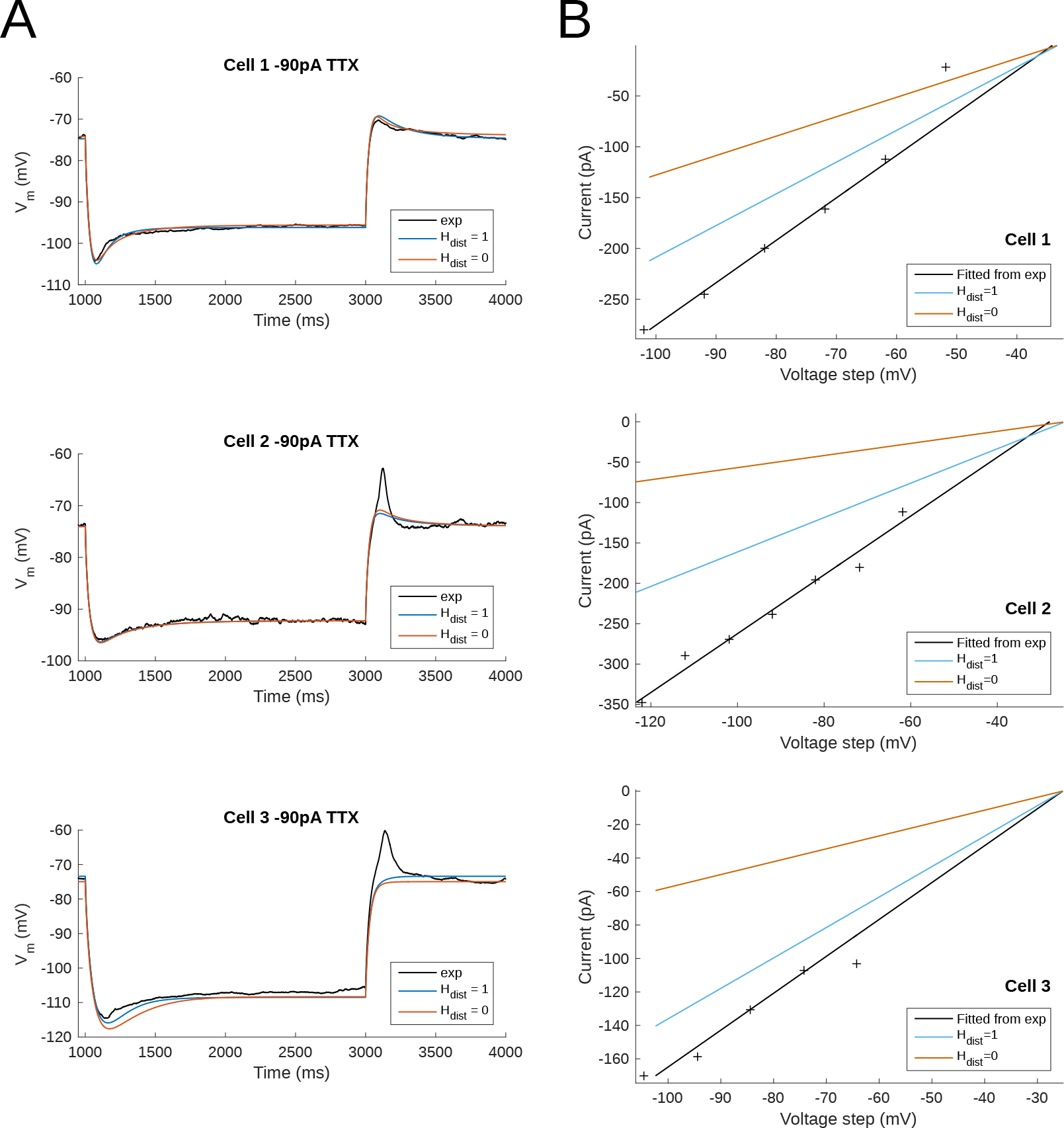
Only OLM Cell Models with Dendritic H-channels are Experimental Data Appropriate. **A.** Using a staggered re-fitting, both *H*_*dist*_=0 or 1 are good fits to the experimental data. Note that the −90pA TTX traces are “test” traces and were not used for fitting (−120pA TTX used for fitting), *H*_*dist*_= 0 or 1. **B.** *H*_*dist*_=1 is clearly more appropriate than *H*_*dist*_=0 relative to the experimental data as shown in plotting I-V curves.

### Immunohistochemistry reveals dendritic HCN2 channel expression in OLM cells

H-channels are encoded in HCN genes (HCN1-4) (*Santoro et al., 2000*; *Santoro and Baram, 2003*). For all three cells where the h-channel experimental data was analyzed, the slowest component of the time constant activation function is around 300 ms, whereas the fast component is less than 100 ms for all three cells. These values coincide with the fast component of the time constant of activation of *I*_*h*_ measured in CA1 pyramidal neurons and oriens/alveus interneurons (of which OLM cells are a population); in both cases, about 100 ms was the fast component and is indicative of heteromeric isoforms of HCN1 and HCN2 subtypes (*Santoro et al., 2000*). Thus, the biophysical properties of *I*_*h*_ were most consistent with the expression of HCN1 and HCN2 channels.

Transgenic deletion of HCN1 and/or HCN2 using CRE/loxP technology has provided additional evidence that HCN1 and HCN2 subunits are the dominant HCN subunits expressed in hippocampal stratum oriens interneurons (*Matt et al., 2011*). HCN1 and HCN2 subunits likely form heteromers that exhibit intermediate biophysical properties and cAMP sensitivity (*Ulens and Tytgat, 2001*). Given the context that HCN1 channel subunits are localized to dendrites of CA1 pyramidal cells (*Lörincz et al., 2002*), whether similar trafficking principles apply to other cell types motivated our studies. Matt and colleagues detected HCN2 immunoreactivity on the somata of somatostatin-positive neurons in the stratum oriens layer. Because somatostatin-immunoreactivity was confined to the somata, it was not clear whether HCN2 immunoreactivity extended to dendritic regions (*Matt et al., 2011*). Moreover, a previous study demonstrated that HCN4 immunoreactivity is present in somatodendritic regions of parvalbumin-positive subpopulations in statum oriens, but it is not clear whether OLM cells were included in this HCN4-containing subpopulation of stratum oriens interneurons (*Hughes et al., 2013*). To investigate the subcellular localization of HCN channels, we employed HCN2 immunocytochemistry in Chrna2-CRE:tdTomato mice. Unlike somatostatin, which labels several hippocampal interneuron subpopulations, the Chrna2 gene encodes for the *α*2 nicotinic acetylcholine receptor, which has an expression pattern that is highly restricted to hippocampal OLM cells (*Mikulovic et al., 2015*; *Urban-Ciecko and Barth, 2016*). Using this transgenic mouse as a tool for visualizing OLM dendrites, we investigated whether HCN2 immunoreactivity was present in the somata only, or extended to dendritic regions of OLM cells.

Although there is 100% HCN2 sequence identity between rat and mouse HCN2, no publication exists that tests the specificity of the HCN2 antibody in mouse. Therefore, we investigated the specificity of the HCN2 antibody using Western blot (see *Figure 6-Figure Supplement 1*). Using protein homogenate isolated from mouse hippocampus, we detected a band at 95 kD, which is the expected molecular weight for HCN2, confirming specificity of the HCN2 antibody. In accordance with previous studies (*Mikulovic et al., 2015*; *Leão et al., 2012*), tdTomato-positive neurons were observed in the stratum oriens layer of hippocampus (*Figure 6*), consistent with the expected anatomical localization of OLM cells. HCN2 immunoreactivity was detected to be co-localized to OLM cells in the stratum oriens, as observed previously (*Matt et al., 2011*). At higher magnification (*Figure 6D,E,H,I*), we observed HCN2 immunoreactivity on proximal dendrites of OLM cells (as indicated by arrows). These data were taken from ventral slices of hippocampus, but we also observed HCN2 immunoreactivity in OLM cell dendrites using dorsal slices, but they were of lower abundance (compare *Figure 6* with *Figure 6-Figure Supplement 2*). We also obtained HCN1 immunoreactivity in OLM cells from dorsal and ventral hippocampal slices (see *Figure 6-Figure Supplement 3*). In summary, these immunohistochemical observations independently corroborate our modelling results that HCN-containing channels are present on both somata and dendrites of OLM cells.

### Optimized full spiking models of OLM cells recapitulate both depolarizing and hyperpolarizing responses to current step stimuli

We have so far developed three multi-compartment models of OLM cells with fitted passive and *I*_*h*_ parameter values. H-channels are present in the dendrites of these models as found to be the most appropriate distribution given the experimental data, and now confirmed with immunohisto-chemical studies (*Figure 6*). We now focus on two of the OLM cell models - *Cell 1* and *Cell 2* - and move forward to include a full repertoire of ion channel types as used in previous OLM cell models (Lawrence et al., 2006b), thus creating full spiking models available for use in further studies.

**Figure 6.**
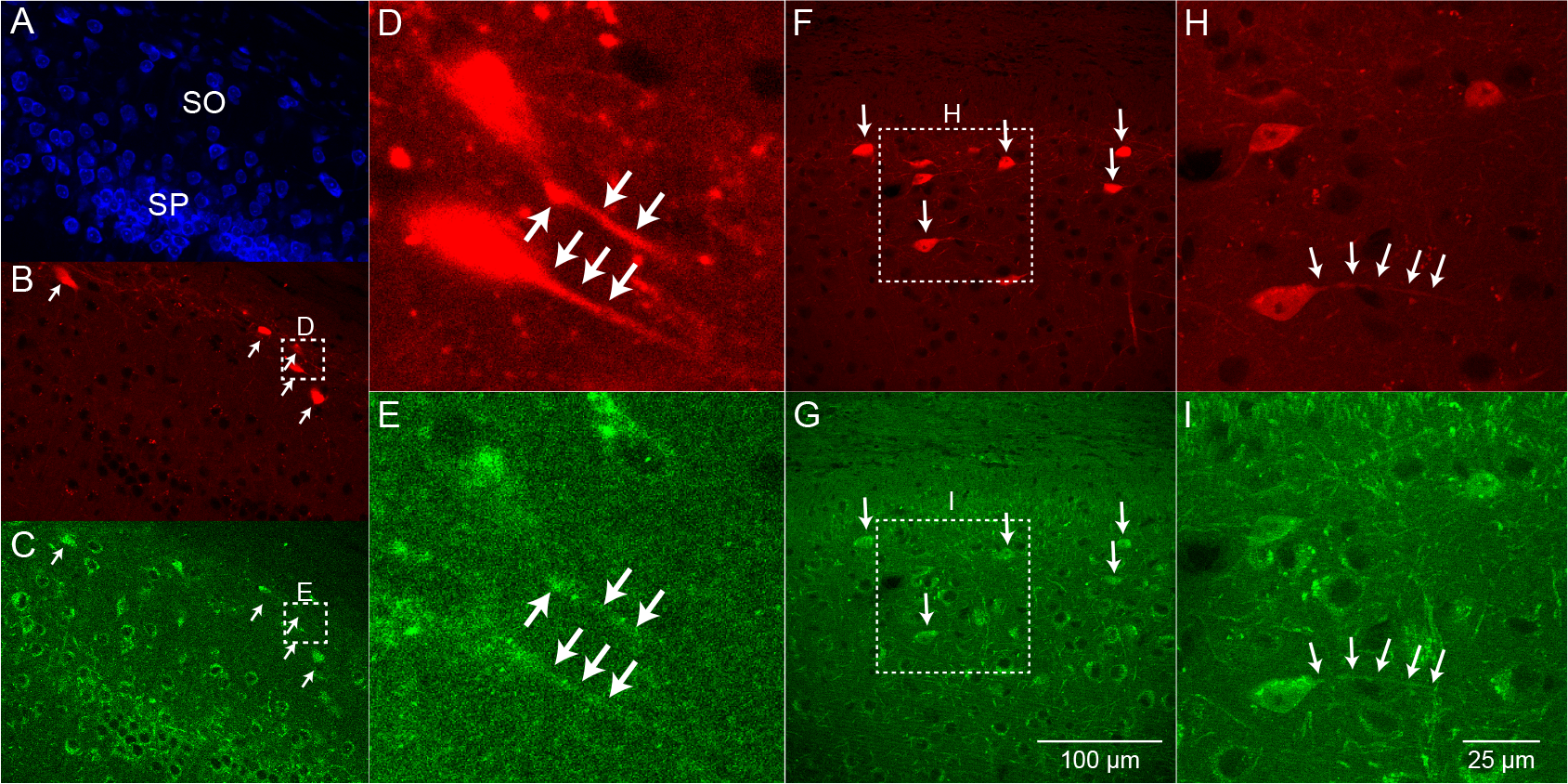
Evidence for HCN2-containing Channels on Dendrites of OLM Cells. **A.** Neurotrace 435, **B.** endogenous tdTomato, and **C.** HCN2 immunofluorescence were imaged in hippocampal slices from Chrna2-CRE:tdTomato mice. D. Expanded view of two OLM cells illustrating E. HCN2 immunofluorescence on OLM cell proximal dendrites. **F., H.** tdTomato images of additional OLM cells with **G., I.** dendritic HCN2 immunofluorescence. Slices obtained from ventral hippocampal CA1. Scale bar in **G.** applies to zoomed out images in **A., B., C., F., G.**. Scale bar in **I.** applies to zoomed in images in **D., E., H., I.**.

To do this, we optimized the parameter values to depolarizing steps of the particular cell, where most voltage-gated ion channels were expected to be activated. Details of the approach used is given in the Methods. In brief, we used BluePyOpt (*Van Geit et al., 2016*) to perform multi-objective optimizations that provided sets of parameter values which generated appropriate OLM cell voltage output at +30 pA, +60 pA and +90 pA depolarizing steps (protocol #2 in *Table 5* in the Methods), and we did our fitting using holding currents in line with the experimental data (4 pA for *Cell 1* and −5 pA for *Cell 2*). We note that our fits were done using the specific experimental data sets and not to a set of experimental data with variances associated with electrophysiological features. The resulting best fits are shown in *Figure 7*, and the next four top fits are shown in *Figure 7-Figure Supplement 3*. The optimized spiking model features relative to the experimental data are shown in *Figure 7-Figure Supplement 4A*, and the optimized parameter values are given in *Figure 7-Figure Supplement 4B,C* and the Methods. Similar outputs were obtained in these top five ranked optimized models and all performed well in terms of capturing electrophysiological feature measurements (*Figure 7-Figure Supplement 4A*). *Cell 2* in particular had more difficulty with the “AHP_depth efeature”, which is likely because the model failed to attain a high-enough spike threshold, and thus the resulting “AHP_depths” were too low. While we tried to encourage the models to reach higher spike thresholds by allowing the sodium voltage-dependencies to vary as free parameters in the optimizations (*Figure 7-Figure Supplement 4C*), in the end, the models could not fully capture the adaptation in spike threshold that was seen experimentally (i.e. the spike threshold appeared to increase during spiking at higher frequencies). These top models also had similar optimized parameter values (*Figure 7-Figure Supplement 4B,C*), though this may be a result of over-constraining the optimizations (see approach and parameter ranges in the Methods).

To ensure that the full spiking models did not affect the *I*_*h*_ fits, we applied hyperpolarizing steps to the full spiking models as done experimentally, and found that they were in full agreement with the experimental data, as shown in *Figure 7-Figure Supplement 1*. It was expected that adding the full set of ion channel mechanisms would not affect the model’s ability to match the hyperpolarizing steps since the additional currents are not active at these hyperpolarized values. This can be appreciated by looking at the contributions from the different currents at the different current steps using “currentscapes”, a novel visualization technique (*Alonso and Marder, 2019*). In *Figure 7-Figure Supplement 2*, it is clear that only *I*_*h*_ and the leak current are active during the hyperpolarization steps, and not other ionic currents. In fact, contributions from all other currents during these hyperpolarization steps were minimized beyond being able to see them on the plots and outward current can become non-existent since the reversal potential for potassium is passed.

**Figure 7.**
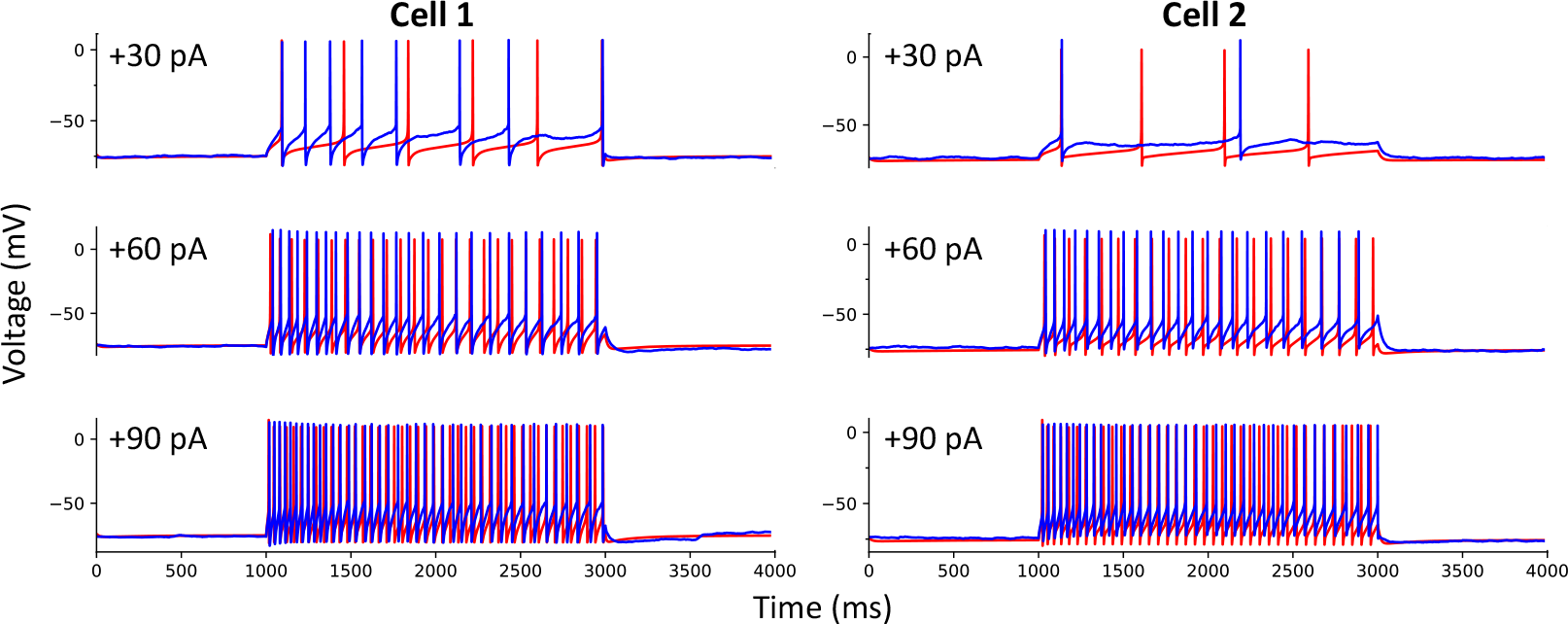
Full Spiking Optimized OLM Cell Models. The most highly ranked optimized models for *Cell 1* and *Cell 2* are plotted in red, and the experimental data is plotted in blue. Model parameters were optimized using depolarizing +30 pA, +60 pA, and +90 pA current step recordings from protocol #2 in *Table 5* specific for *Cell 1* and *Cell 2*.

Taking advantage of our generated currentscapes (*Figure 7-Figure Supplement 2*), we were able to easily observe several features in our optimized models. A prominent feature was the large contributions from A-type potassium currents during both the baseline periods as well as during spiking regime activities. Since we minimized slow delayed rectifier potassium conductance on purpose in order to achieve better fits (see Methods and *Figure 7-Figure Supplement 4*), it was not surprising that the major contribution of outward currents during spikes was from fast delayed rectifier potassium. However, it was perhaps surprising that M-type and calcium-activated potassium currents provided such large contributions to outward currents, despite having considerably smaller conductances relative to the other outward ion channel types (*Figure 7, Figure Supplement 4*). Particularly, M-type exhibited larger current contributions during the after-hyperpolarization periods (AHP) at higher spike rates. In terms of inward current contributions, we did not see any observable contributions from the L-type and T-type calcium channel types. Mostly, inward current contributions in the spiking regimes were from sodium channels. However, *I*_*h*_ provided some observable contributions during the spike recovery periods, and also provided a larger contribution leading up to the first spike.

We note that our goal was to obtain spiking models that could adequately recapitulate the data for the particular cell, that is, starting idealized “base” models of OLM cells. These base models should be further explored for degeneracy and can be leveraged for additional insights and hypothesis generation moving forward (see Discussion). However, they represent the most comprehensive multi-compartment models of OLM cells to date, having been produced using morphologies and electrophysiological recordings obtained from the same biological OLM cells.

## Discussion

In this work, we obtained a set of recordings from OLM cells in hippocampal CA1 that allowed us to explicitly link morphological, passive and h-channel biophysical parameters within the same cell. From this set, we constructed three “next generation” experimentally-constrained multi-compartment models of CA1 OLM cells. The models developed here are considered next generation in that, unlike all previous computational models of OLM cells (*Skinner and Ferguson, 2018*), we have here for the first time characterized the h-channels on an individual cell basis, and further, matched morphology and electrophysiology to constrain two full spiking models of OLM cells. Our models robustly predicted that h-channels are expressed on the dendrites of OLM cells. Moreover, through an independent line of experiments, we confirmed this to be the case using immunohistochemical staining of HCN2-containing subunits. These lines of evidence converge on the conclusion that h-channel expression extends into the dendrites of OLM cells. Our models can be used in future studies to explore the synaptic and network consequences of dendritic HCN channel expression in OLM cells within the context of hippocampal microcircuit function.

### Computational and experimental investigations in OLM cells converge on somato-dendritic HCN channel expression

The existence of h-channels, mixed cation channels that activate with hyperpolarization, has long been known since first discovered as “funny” currents in the heart (*Brown et al., 1979*), and in the CNS, they contribute to maintenance of the RMP, pacemaking ability, and synaptic integration (*Biel et al., 2009*; *Lörincz et al., 2002*; *Magee, 1998*). The contribution of h-channels in pyramidal cells to subthreshold resonance and spiking output features in hippocampus and cortex has been much studied (*Biel et al., 2009*; *Narayanan and Johnston, 2012*; *Santoro and Baram, 2003*; *Zemankovics et al., 2010*). In particular, it is known that the distribution of HCN1-containing channels increases from soma to distal dendrite and as such, have been shown to control the temporal summation of synaptic inputs from dendrites to soma (*Magee, 1998*; *Vaidya and Johnston, 2013*). However, h-channels in cerebellar Purkinje neurons are uniformly distributed in their dendrites and do not strongly affect temporal summation to the soma (*Angelo et al., 2007*). For interneurons, and OLM cells in particular, it is known that they express HCN channels, as seen by a large sag upon hyperpolarization (*Maccaferri and McBain, 1996*). However, prior to this work, it has been entirely unknown whether OLM cells express h-channels on their dendrites.

H-channels in OLM cells have been implicated in pacemaking and oscillatory activities of the hippocampus (*Gloveli et al., 2005*; *Maccaferri and McBain, 1996*), and theta (4-12 Hz) rhythms in particular (*Maccaferri and Lacaille, 2003*; *Rotstein et al., 2005*). Subsequent experimental studies found that OLM cells did not have any preferred spiking frequency response to broadband arti1cial synaptic inputs (*Kispersky et al., 2012*). *Kispersky et al. (2012)* did find, however, that OLM cells exhibited a phase-locked spiking preference to theta frequency modulated inputs, but this spike resonance did not depend on h-channels. However, these frequency modulated synaptic inputs were delivered exclusively to the soma of OLM cells via dynamic clamp technology. Using computational model databases of OLM cells in the absence or presence of h-channels in dendritic compartments, our previous studies revealed that OLM cells modelled to be in a simplified *in vivo*-like scenario could exhibit a theta frequency spiking resonance when inputs were delivered to their dendrites (*Sekulić and Skinner, 2017*). We further found that a high or low theta frequency spike resonance was possible and is respectively dependent on whether h-channels were present in the dendrites or not of the OLM cell models, reminiscent of Type 1 and 2 theta rhythms in the behaving animal (*Kramis et al., 1975*). Our modeling work examining dendritic distributions of h-channels in OLM cells found that the distributions could vary so long as total conductance was conserved (*Sekulić et al., 2015*), as was also found in Purkinje cells (*Angelo et al., 2007*). Thus a motivating factor in the present study was to constrain this extra ‘free parameter’ by obtaining direct measurements of the total conductance in OLM cells. In doing this, we were able to show that our OLM cell models best matched the experimental data if h-channels are present in the dendrites. Interestingly, while the total h-channel conductance ranged from 2.2-4.2 nS in the three cells that were fully analyzed (*Table 4*), the conductance density in each of the three cells is about 0.1 pS/*μ*m^2^, which is the value found in highly ranked OLM cell models from our previously developed model databases (*Sekulić et al., 2014*; *Sekulić and Skinner, 2017*). *Zemankovics et al. (2010)* obtained total conductance values averaging approximately 4 nS, which are near the upper limit of our measurements. However, total h-channel conductance values were obtained in OLM cells in rat, which had a two-fold larger measured capacitance (208 pF) than our mouse OLM cells (107 pF). Therefore, given the difference in measured surface area between rat and mouse, our data is in accordance with this previous study. Compared to *Maccaferri and McBain (1996)* who obtained a mean reversal potential of −32.9 mV, an activation curve with a half-activation voltage (*V*_1/2_) of −84.1 mV, and slope factor (*k*) of −10.2 mV, our reversal potentials ranged from −25.2 to −34 mV, with model *V*_1/2_ fits of −99.6 to −113.8 mV, and *k* of −9.99 mV (*Table 4*). The voltage-dependence of the time constant yielded fits that were different but with overlapping values for the three cells (*Figure 3B*).

This model prediction was borne out by our immunohistochemistry studies showing HCN2 in the dendrites of OLM cells. specifically, we found expression of HCN2 in dendrites of ventral (*Figure 6*) and dorsal (*Figure 6-Figure Supplement 2*) OLM cells. Although we attempted to obtain evidence of HCN1 localization, the low signal-to-noise of the antibody staining did not permit an unambiguous determination of HCN1 channels on OLM cell dendrites of either ventral or dorsal populations (*Figure 6-Figure Supplement 3*). A previous study found HCN2 expression in somatic regions of OLM cells, but did not examine dendritic expression (*Matt et al., 2011*). As far as we know, our study is the first to present immunohistochemical evidence for the expression of h-channels in the dendrites of a significant population of OLM cells. Moreover, given the evidence that HCN1 subunits are expressed in OLM cells (*Matt et al., 2011*), dendritic localization of HCN2 subunits, formation of heterodimers with HCN2 subunits (*Ulens and Tytgat, 2001*), and biophysical properties of OLM cell h-channels consistent with HCN1/2 expression (*Figure 3*), it is possible that HCN1 subunits are co-expressed with HCN2 subunits on OLM cell dendrites. As such, there may be common cellular rules for subcellular trafficking of HCN1/HCN2 subunits in OLM and CA1 pyramidal cells (*Lörincz et al., 2002*).

### Cellular and synaptic consequences of somatodendritic HCN channels in OLM cells on hippocampal microcircuit operations

In building our next generation OLM cell models using morphological and electrophysiological data from the same cell, we were able to robustly show, and thus predict, the presence of h-channels in the dendrites of OLM cells. In doing this, it was critically important that the experimental data came from the same cell. Recently OLM cells were discovered to be comprised of parvalbumin- and 5HT3 subtypes (*Chittajallu et al., 2013*). With the advent of sophisticated genetic sequencing techniques (*Cembrowski and Spruston, 2019*; *Harris et al., 2018*), additional OLM cell subtypes can be recognized. Whether Chrna2-expressing OLM cells are a distinct OLM subpopulation or preferentially fall into an existing subclass is not yet known. Moreover, it was noted that CA1 pyramidal cells have a continuous, rather than discrete, variation on the longitudinal axis of the hippocampus, indicating this as an organizational principle (*Cembrowski et al., 2016*), and structural-functional correlations are apparent for ventral, intermediate and dorsal regions of the long axis (*Fanselow and Dong, 2010*). Given this observation, it is interesting to note that *Cell 1* and *Cell 2* from an intermediate CA1 region have more similar characteristics than *Cell 3* which is from a more ventral CA1 region (see *Figure 3*).

It has been proposed that OLM cells play a gating role (*Leão et al., 2012*), akin to earlier work by *Blasco-Ibáñez and Freund (1995)* who showed that “horizontal SOM+ interneurons” (i.e., putative OLM cells) could act as a switch controlling activation of local pyramidal cells via Schaeffer collaterals or perforant path input from entorhinal cortex. Further work has shown that OLM cells in intermediate regions of CA1 exert a bidirectional control on learning and memory (*Siwani et al., 2018*), and ventral OLM cells control Type 2 theta rhythms and are associated with increased risk-taking (*Mikulovic et al., 2018*). In a recent modeling study, OLM cells were shown to be critical in producing a robust intrinsic theta output (*Chatzikalymniou and Skinner, 2018*), which suggests that their neuromodulation may be key to the maintenance of theta rhythms.

OLM cells receive inputs from various sources that include local excitatory and inhibitory CA1 neurons, and medial septal/diagonal band of Broca (MS-DBB) inputs (*Chamberland et al., 2010*; Lawrence et al., 2006a; *Leão et al., 2012*; *Lovett-Barron and Losonczy, 2014*). Excitatory inputs from MS-DBB glutamatergic neurons have activity that precedes and controls the initiation of both hippocampal theta activity as well as animal locomotion (*Fuhrmann et al., 2015*). OLM cells also receive cholinergic inputs from a separate population of MS-DBB cells, whose activation onto muscarinic receptors promotes excitability and enhances oscillatory activity of OLM cells at theta frequencies (Lawrence et al., 2006a,c). Still further, OLM cells receive GABAergic inputs from a third population of MS-DBB neurons that express GABA, thought to rhythmically inhibit and thus pace OLM cells at theta frequencies (*Borhegyi et al., 2004*; *Gulyás et al., 1990*). The interpretation of rhythmic inhibition by MS-DBB GABAergic cells of OLM cell activity at theta frequencies nevertheless leading to recruitment of OLM cell theta frequency firing is supported by our modeling work (*Sekulić and Skinner, 2017*) where we showed that modulation of inhibitory synaptic input preferentially enhances spike recruitment of OLM cell models at theta frequencies. At a behavioural level, recent work has shown that selective optogenetic silencing of MS-GABAergic neurons to reduce theta did not affect memory if it occurred outside of REM sleep (*Boyce et al., 2016*). The exact mechanisms of theta transmission from the MS-DBB to the hippocampus are likely to be complex and depend on the specific timing of the incoming excitatory and inhibitory pathways. It is possible that spatial as well as temporal patterning of synaptic inputs onto the dendritic trees of OLM cells are precisely arranged to be able to recruit OLM cell activity during theta rhythms. Thus, the dendritic distributions of voltage-gated channels, including but perhaps not limited to HCN1/2 channels, may be arranged in conjunction with the spatial pattern of synaptic inputs to allow OLM cells to participate in hippocampal theta activity in a behavioural context-dependent way. The cAMP-dependent neuromodulation of somatodendritic HCN1/2 channels in OLM cells by synaptically released norepinephrine onto beta-adrenergic receptors (*Maccaferri and McBain, 1996*; *Sekulić and Skinner, 2017*) will increase cellular excitability. Combined with increased activation of MS-DBB GABAergic afferents, this neuromodulation would promote rebound spiking in OLM cells and facilitate hippocampal encoding and novelty detection operations.

### Exposing and exploiting limitations in experiments and multi-compartment model development, and a ‘cycling’ strategy

It was initially unexpected that a model with fitted passive properties and morphologies obtained in conjunction with h-channel parameters extracted from the same cell did not produce voltage recordings that matched those from experiment when all channel types except h-channels were blocked (*Figure 4A*). To explain why this may be the case, some general issues in building multi-compartment models directly from limited experimental data need to be considered.

The experimental data obtained from the OLM cells here, used to extract both passive and h-channel characteristics, were not perfectly optimal. In an attempt to constrain as many distinct parameters within the same cell as possible, we deliberately sacrificed depth for breadth so that practical choices were inevitable in the distribution of efforts. There are inherent limitations to cell stability that require rapid succession through a sequence of experimental protocols (*Table 5*). In our hands, the limit of stability was approximately thirty minutes. In this time, we were able to obtain recordings, bath changes, and biocytin fills that allowed us to do reconstructions, and obtain passive property and h-channel biophysical properties, but having several protocols prevented multiple sweeps of any given protocol. The I-V relation for determining maximum conductance and reversal potential was not always linear across all voltage steps, as required from theoretical perspectives in the mathematical model formulations. Furthermore, there was some error associated with fitting the Boltzmann function describing the steady-state activation curves to all data points obtained from the h-channel activation protocol. Indeed, due to inherent biological variability and experimental constraints, some measure of error is expected whenever experimental data is fitted to theoretical or mathematical models, such as a Boltzmann function for the activation curves, or a dual exponential function for the time constant of activation. Accordingly, although we obtained the requisite experimental data for fully characterizing h-channels and fitting to mathematical models of them, we should not expect that the resulting parameters will necessarily result in fully appropriate cellular output when initially used. That is, even when inserted into multi-compartment models built of the same cells from which the h-channel characteristics were obtained, there may be error in the resulting model’s *V*_*m*_’s output compared to that of the experimental recordings. In essence, this is due to the accumulation of errors in estimating the various parameters used, and is compounded with increasing number of experimentally-constrained parameters in the model.

To overcome this, we found that an approach of a staggered re-fitting of the parameters in the model was able to produce generalizable results so that the *V*_*m*_ output could match all of the experimental traces including those for which it was not specifically optimized. This procedure can be thought of as correcting for errors in the procedures for extracting the parameter values from the experimental data, using the same principles for correcting for errors in the morphological reconstructions by fitting the passive properties. Having many recordings from the same cell allowed us to do a staggered re-fitting of model parameters that avoided overfitting and allowed validation, as well as consideration of the voltage dependence of h-channel activation time constants. It may be possible to use more sophisticated optimization schemes to obtain generalized fits, but the challenge of fitting detailed multi-compartment models with many parameters to experimental data is recognized, and has led to use of two-stage fitting processes (*Hay et al., 2011*; *Roth and Bahl, 2009*). We note that our staggered re-fitting can be considered as a form of two-stage fitting where in our situation, we determined how to proceed with the re-fitting stages based on how robust the experimental recordings were considered to be. A further limitation of the electrophysiological data was that our recordings were somatic. Due to the relatively compact nature of OLM cell dendrites, this is not a major limitation unlike what it may be for pyramidal cells which have extended dendritic trees. However, in the end, we obtained a strong prediction of h-channels being present in the dendrites since all three individual model fits supported this interpretation, indicating the robustness of our staggered re-fitting procedure.

We obtained specific capacitance values that were lower than the ‘typical’ values of 1.0 *μ*F/cm^2^ (*Gentet et al., 2000*). This intriguing result requires experimental validation in the form of nucleated patch recordings as done previously for directly measuring the specific capacitance in other neurons (*Eyal et al., 2016*; *Gentet et al., 2000*). It is unlikely that our low specific capacitance values are due to surface area estimation errors, as even if we had overestimated our surface areas by double, specific capacitance values would still be ≈0.5 *μ*F/cm^2^ (see *Table 2*). It is interesting to note that in the case of human neurons where values of ≈0.5 *μ*F/cm^2^ were reported (*Eyal et al., 2016*), another group has reported values of ≈0.9 *μ*F/cm^2^ (*Beaulieu-Laroche et al., 2018*).

Clearly, it is important to keep in mind what one’s goal(s) are in the building of a multi-compartment model in the first place. Without making some simplifying assumptions, such as uniform passive properties, and having constraining experimental data, we are necessarily faced with the curse of dimensionality (*Almog and Korngreen, 2016*). In our original multi-compartment models of OLM cells (*Saraga et al., 2003*), we were motivated to include dendrites because of clear evidence of highly active dendrites (*Martina et al., 2000*) in OLM cells. Moving forward, we expanded the extent of ion channels present in the models when recordings specific to M-channels in OLM cells were performed (Lawrence et al., 2006b). A key notion in experimentally-constrained computational modelling is that the models are never complete. There should always be a reciprocal transfer of knowledge between model and experiment where experimental data is used to constrain models which, in turn, both point out gaps in our understanding of the underlying cellular neurophysiology as well as generate hypotheses, refine protocols, and consider additional measurable parameters that can then be incorporated into future model revisions.

A particular conceptualization of the role of computational modelling in neuroscience is to help resolve, or at least reframe, these basic concerns of how “realistic” detailed models can be. Rather than the idea of obtaining a detailed model as a crystallized end point of any given study, we consider the role of the detailed modelling as an integral component of a cyclical process of knowledge generation in neuroscience. We have expressed this as the experiment-modelling cycling approach (*Sekulić and Skinner, 2018*). Although the approach was initially formulated in the context of population or database modelling, it can be generalized for any computational model whose goal is to explain experimental data, develop hypotheses and make predictions. This conceptualization states from the outset that the goal of modelling is not to find optimal or realistic models per se, but rather to develop models in such a way that a specific physiological question is raised and can lead to experimental examinations. In our initial studies, we asked the question of whether OLM cells expressed h-channels in their dendrites and thus built OLM cell model databases that either did or did not have h-channels in their dendrites (*Sekulić et al., 2014*). The most relevant aspect of this approach in terms of answering the question of whether models are realistic or not is the recognition that the process is cyclical. Thus, we consider that an essential goal in multi-compartment modeling is the back-and-forth cycling between experiments and models that leads to continual refinement of the model relative to the biological cell, thus allowing for the generation of predictions for further experiments, and thus knowledge generation in neuroscience.

### Limitations and future work

Although doing more than three full reconstructions, analysis and multi-compartment model building may be desirable, we felt that consistently obtaining best matches with dendritic h-channels in all three of our models when 1t with data from the same cell was enough to allow for conclusions as to dendritic expression of h-channels in OLM cells. Also, we focused on uniform h-channel distribution in the dendrites since our starting models using either no h-channels or h-channels fully and uniformly distributed in the dendrites did not match the experimental data (*Figure 4A*). Considering distributions that were not uniformly distributed (e.g., distributed only in proximal dendrites) would be unlikely to capture the data given that the total h-channel conductance would remain the same.

Our development of full spiking OLM cell models here, as based on *Cell 1* and *Cell 2*, are available for future use and provided online (see Methods). In particular, it would be interesting to use currentscape visualization analyses (*Alonso and Marder, 2019*) to help disentangle the interacting dynamics, perhaps using it to direct how one might best reduce the model complexity to allow dynamical system analyses to be applied, as well as applying sensitivity analysis techniques such as uncertainpy (*Tennøe et al., 2018*). In turn, this could help decipher how OLM cells preferentially respond to different theta frequencies based on their biophysical profile as shown in our previous computational models (*Sekulić and Skinner, 2017*). Further, these models now provide a foundation or canonical start for the creation of new databases designed to address specific biophysical questions as done in our original database that was developed to ask whether h-channels were present in the dendrites (*Sekulić et al., 2014*). Interestingly, co-regulation of h-channels and A-type channels are apparent in the currentscapes (see *Figure 7-Figure Supplement 2*) as was observed in our original OLM cell databases.

We have previously shown that using virtual networks, or creating *in vivo*-like representations with multi-compartment cellular models, as done with our earlier OLM cell models (*Sekulić and Skinner, 2017*), can lead to insights of circuit function from cellular specifics. We have also created *in vivo*-like states with interneuron-specific interneuron models (*Guet-McCreight and Skinner, 2019*), and used them to make links between *in vitro* and *in vivo* studies (*Luo et al., 2018*). In essence, it seems possible that an understanding of the contribution of biophysical cellular details to circuits in the behaving animal can emerge by using virtual networks.

In a review, *Almog and Korngreen (2016)* demonstrate the limitations associated with the re-usability of layer 5 pyramidal cell models, and also state that there is a need for proving that multi-compartment models are valid within the context of network simulations. These are challenging issues to consider but an important step that they suggest is to ensure that models are linked with the experimental data. Along these lines, neuroinformatic tool developments (e.g., Nexus - https://bluebrainnexus.io) can help reduce the workload.

In conclusion, we have achieved our goal and our work has shown that if the development of multi-compartment models are done for a specific cell type in which ion channel characterization and morphological and passive data can be obtained from the same cell for several cells, it is possible to determine their ion channel distribution and biophysical characterization.

## Methods and Materials

### Ethics statement

All procedures were performed in accordance with the University of Montana (Animal Use Protocols 026-11 and 017-14) and Texas Tech University Health Sciences Center (Animal Use Protocols 15025, 15031 and 16037) Institutional Animal Care and Use Committees.

### Brain slice preparation

Transverse hippocampal slices were prepared as described previously *Yi et al. (2014)*. Briefly, SOM-CRE^+/−^:Rosa26YFP^+/−^ (SOM-YFP) mice of both genders (9-10 weeks) were anesthetized with isoflurane and then transcardially perfused with ice-cold partial sucrose solution (PSS) containing (mM): 80 NaCl, 2.5 KCl, 24 NaHCO_3_, 0.5 CaCl_2_, 4 MgCl_2_, 1.25 NaH_2_PO_4_, 25 glucose, 75 sucrose, 1 ascorbic acid, 3 sodium pyruvate, saturated with 95% O2/5% CO_2_, pH 7.4 *Bischofberger et al. (2006)*. After carefully extracting, blocking, and mounting the brain, transverse hippocampal slices (300 μm) were cut in ice-cold oxygenated PSS with a 1200 S Vibratome (with Vibrocheck accessory; Leica Microsystems, Bannockburn, IL, USA), and then were incubated in warm (36°C) oxygenated PSS at least 30 min before use.

### Chemical reagents

DL-APV was purchased from R&D Systems (Minneapolis, MN, USA). Tetrodotoxin (cat# 5651), TEA (cat# 2265), 4-AP (cat# A78403), DNQX (cat# D0540), SR-95531 (cat# S106), and ZD7288 hydrate (cat# Z3777) were purchased from Sigma-Aldrich, Inc. (Saint Louis, MO, USA). Salts and chemicals for saline solutions, including biocytin, were also purchased from Sigma-Aldrich, Inc.

### Western Blot methods

Mouse hippocampi and the whole brains (without hippocampi) were collected separately, sonicated and lysed in Radioprecipitation Assay Lysis buffer (5M NaCl, 0.5M EDTA pH 8.0, 1M Tris pH 8.0, 1% NP-40, 10% sodium deoxycholate, 10% SDS). Halt™ Protease inhibitor cocktail 100x (ThermoFisher, cat# 87786) was added before use. The denatured samples were resolved on a 4-20% Mini-PROTEAN TGX Stain-Free precast protein gel (Bio-Rad, cat# 4568094) and transferred to a nitrocellulose membrane (0.45 μm, Bio-Rad, cat# 1620115). The membrane was then blocked in 6% milk (diluted in TBS) for 45 min at room temperature, incubated with primary antibody (1:200, rabbit anti-HCN2, Alomone, cat# APC-030) in 2% milk overnight at 4°C, followed by goat-anti-rabbit secondary antibody (1:10,000, IRDye 800CW, LI-COR) in 2% milk for 1 hour at room temperature. TBST was used for all the washes after antibody incubation (5 x 5 minutes). The near infrared immunoreactivity bands were imaged using Odyssey CLx Imaging System (LI-COR) and converted to grayscale with Photoshop.

### Electrophysiological recordings and analyses

Hippocampal slices were transferred to a recording chamber and submerged in artificial cerebrospinal fluid (ACSF) solution containing (mM): 125 NaCl, 2.5 KCl, 25 NaHCO_3_, 2 CaCl_2_, 1 MgCl_2_, 1.25 NaH_2_PO_4_ and 20 glucose, saturated with 95% O_2_/5% CO_2_, pH7.4, at 34–35°C. SOM-YFP cells in the CA1 stratum oriens layer of hippocampus were visualized using IR-Dodt contrast and fluorescence video-microscopy (Zeiss Axiovision 4.7) on either a Patch Pro 2000 (Scientifica Ltd, Uckfield, East Sussex, UK) or Infrapatch (Luigs and Neumann, Ratingen, Germany) on an upright Zeiss microscope (Axio Examiner; Carl Zeiss Microscopy, LLC, Thornwood, NY, USA). On the Patch Pro 2000, live YFP-positive cells were visualized with a 505 nm LED (LED4C11-SP; Thorlabs) driven by a four-channel LED driver (DC4100; Thorlabs). On the Infrapatch rig, a 505 nm LED was controlled by the Colibri LED illumination system (Carl Zeiss Microscopy). Patch pipettes (2-4 MΩ) were fabricated using a two-step vertical electrode puller (PC-10; Narishige, East Meadow, NY, USA) and filled with internal solution containing (mM): 110 potassium gluconate, 40 KCl, 10 HEPES, 0.1 EGTA, 4 MgATP, 0.3 Na_2_GTP, 10 phosphocreatine and biocytin 0.2%, titrated to pH 7.2 with KOH, osmolarity 295-305 mOsm/L. Whole cell recordings were made using a Multiclamp 700B amplifier (Molecular Devices, Union City, CA, USA), filtered at 4 kHz, and digitized at 20 kHz (Digidata 1440A; Molecular Devices). Current and voltage traces were acquired on a PC running Axograph X (Axograph Scientific, Sydney, Australia). Solutions were heated to 34-35°C with an inline solution heater (HPT-2, Scientifica; SH-27B/TC-324B, Warner, Hamden, CT, USA). Access resistance (R_s_) was monitored during recording. Cells with initial R_s_ less than 20 MΩ were recorded. If R_s_ changed more than 20% during the course of the whole cell recording, the data were excluded from further analyses. In all recordings, the AMPA receptor antagonist DNQX (25 *μ*M), the NMDA receptor antagonist DL-APV (*μ*M), and the GABA_A_ receptor antagonist SR-95531 (gabazine; 5 *μ*M) were included in the ACSF. For blocking intrinsic voltage-gated channels to obtain I_h_, TEA (10 mM), 4-AP (5 mM), and TTX (1 *μ*M) were applied. The I_h_-specific blocker ZD7288 (10 *μ*M) was used to obtain the I_h_-sensitive current and to constrain I_h_ parameters on a per-cell basis.

The order of protocols is important to consider during the subsequent procedures of obtaining OLM cell passive properties in light of varying stages of cell health and deterioration as the recordings progressed. The chronological order of current clamp and voltage clamp experimental protocols performed are shown in *Table 5*. The approximate length of experiment for a given cell patched was at most 30 min. At the end of the recording, pipettes were withdrawn to outside-out patch configuration. Slices were kept on the rig for several minutes to facilitate diffusion of biocytin to distant subcellular compartments. Electrophysiological data were analyzed with Axograph X. The junction potential was calculated to be 11.88 mV and was subtracted from all experimentally recorded voltage values prior to use in subsequent data analysis and creation of multi-compartment computational models.

### Visualization of biocytin-filled cells and confocal imaging

During electrophysiological experiments, recorded SOM-YFP cells were filled with biocytin for post hoc morphological reconstruction. After recording, slices were fixed overnight at 4°C in 0.1 M phosphate-buffered saline (PBS) containing 4% paraformaldehyde. After several washes in PBS, and 2 hours permeabilization with 0.3% Triton X-100 in PBS at room temperature, slices were incubated overnight at 16°C in PBS with Alexa 633-conjugated streptavidin (final concentration 1 μg/mL, catalogue no. S-21375; Invitrogen). Slices were cryopreserved in 30% sucrose containing PBS and then re-sectioned at 100-150 *μ*m thickness using a sliding freezing microtome (HM430; Thermo Scientific, Waltham, MA, USA). After staining with Neurotrace 435/455 (1:100 in PBS) and mounting on gelatin-coated slides in Vectashield (catalogue no. H-1400; Vector Laboratories), sections were imaged with a Fluoview FV-1000 confocal imaging system (Olympus) with a 60× objective. Confocal stacks (800 × 800 pixels; 0.2 *μ*m z-step) of SOM-YFP cells were flat projected, rotated and cropped in PhotoShop 13.0 or ImageJ for display.

### Morphological reconstruction of OLM cells

ImageJ was used as a general purpose image processor including greyscale conversion and bleach correction (*Schneider et al., 2012*). XuvTools was used for stitching of confocal images (Emmen-lauer et al., 2009). Bitplane Imaris was used for viewing reconstructions in 3D and for validating the z-stack. Finally, volume reconstruction and specification of geometric models of neuronal morphologies was performed using Neuromantic (*Myatt et al., 2012*). Confocal microscope images at 60X magnification were acquired for the cells used in this work. The field of view of each image was restricted to 200×200 μm, resulting in 2-11 image “stacks” per cell. The microscope step size was 0.2 *μ*m in the Z-plane, resulting in 150-200 images per stack. Variation in contrast between stacks were likely due to photobleaching, as stacks acquired later in the image acquisition process for each cell were more apparently bleached than the ones acquired earlier. Bleach correction was performed using ImageJ by normalizing the contrast of all stacks for each cell according to the average intensity value across all stacks per cell. Stacks were then stitched together to recover the volume information for the entire cell. Stitching was performed using the XuvTools software package (*Emmenlauer et al., 2009*). The algorithm implemented in XuvTools utilizes an approach that calculates the correlation between transformations of successive pairs of 3D images while performing a translation or displacement operation of each image. The transformations consist of phase information extracted from discrete Fourier transforms of the images. When displaced images are compared using the phase information from the Fourier transform – referred to as the phase-only correlation (POC) – there is a precise peak in the POC function that corresponds to the estimate of the magnitude of displacement and can thus serve as a marker for how much displacement is needed to allow for maximum overlap between the images (*Emmenlauer et al., 2009*; *Ito et al., 2006*). An efficient implementation in XuvTools takes advantage of the finding that even when downscaling the quality of the image when performing comparisons in order to save on computational runtime, the resulting POC still exhibits a clearly defined peak at the location of displacement with no local maxima close to the peak (*Emmenlauer et al., 2009*). Accordingly, the algorithm is supposed to be robust to noise and is invariant to linear changes in gray values. In terms of practical application, however, we found that the success of the XuvTools algorithm varied depending on whether bleach correction was performed or not. In the case of several cells, the confocal stacks could not be stitched prior to bleach correction. A possible explanation for this is that the bleaching may not result in a strictly linear shift in contrast but rather skews the histogram of intensity values towards brighter values. Thus, XuvTools would not be expected to find a well defined peak in the POC function between the images because the two images are then not simply shifted versions of one another but in fact decorrelated with respect to the phase of image intensity extracted by the Fourier transform. This would result in an attenuated peak in the POC function, leading to less certainty in how much displacement is needed to find the optimal overlap between image pairs. We next performed volumetric reconstruction of the soma, dendrites, and axons. This was done using the freely-available Neuromantic software package that implements semi-automated tracing (*Myatt et al., 2012*). The semi-automated tracing procedure resulted in successful tracing of several cells in the experimental dataset used here, but with a fundamental limitation being that 1 *μ*m was the minimum possible diameter hard-coded in Neuromantic. The resulting surface areas of the traced cells were too large, and distal dendritic diameters were clearly overestimated. Inhibitory interneurons may possess dendrites with thickness less than a micron, e.g., 0.4 *μ*m in cerebellar interneurons (*Abrahamsson et al., 2012*). Accordingly, we performed full manual reconstructions using Neuromantic.

**Table 5.**
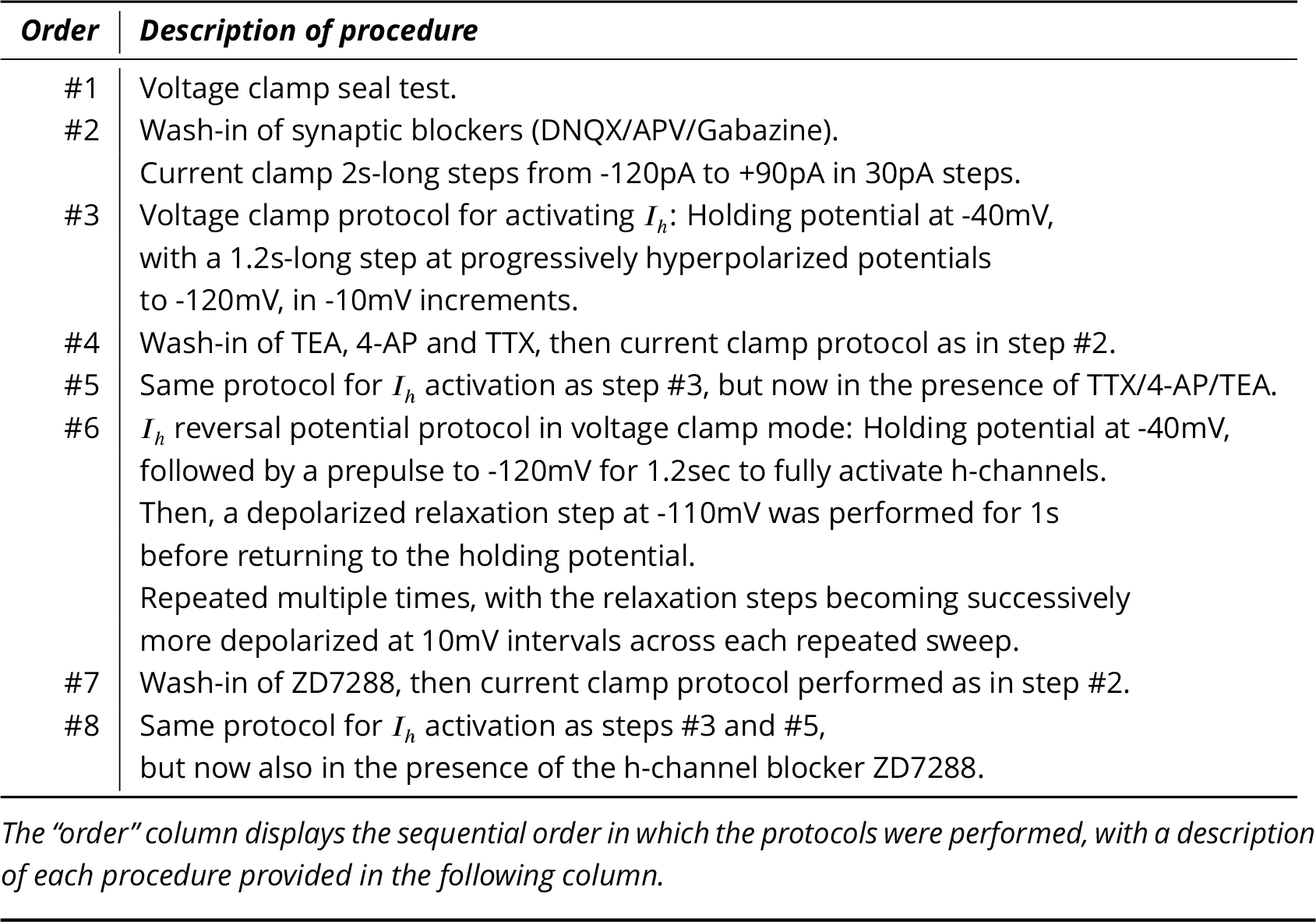
Order of experimental protocols performed on OLM cells.

### Immunohistochemistry and confocal imaging

Heterozygous Chrna2-CRE^+/−^:td-Tomato^+/−^ mice (*Mikulovic et al., 2015*; *Leão et al., 2012*) were transcardially perfused with fresh 4% paraformaldehyde (PFA) in 0.1 M phosphate-buffered saline (PBS). The brains were post-fixed with the same fixative overnight and stored in 30% sucrose in PBS at 4°C for at least 48 hours or until fully submerged in 30% sucrose. Brains were then frozen with Fisher Healthcare™ Tissue-Plus™ O.C.T Compound 4585 (Fisher Scientific, cat# 23-730-571) using dry ice in 100% ethanol, and sectioned coronally at 45 *μ*m thickness using a HM430 sliding microtome (ThermoFisher). Free floating brain slices containing hippocampus were washed with PBS (3 × 10 min) and permeablized with 0.3% Triton™ X-100 (Sigma-Aldrich, cat # X100) in PBS. Blocking was performed using a 30%/70% ratio of antibody diluent solution (ADS; 1% bovine serum albumin and 0.3% Triton™ X-100 in PBS) to 10% goat serum, respectively, for 2 hours at room temperature. Sections were then washed in PBS (3 × 10 min) and incubated with rabbit anti-HCN2 primary antibody (0.8 mg/ml in ADS, Alomone Labs, cat# APC-030) overnight at 4°C using an orbital shaker (Multi-Purpose Rotator, ThermoScientific). The following day, sections were washed in PBS (3 × 10 min) and incubated in goat anti-rabbit AlexaFluor 488 secondary antibody (1:500, ThermoFisher, cat# A-11008) for 2 hours in PBS on an orbital shaker. Finally, following PBS washes (3 × 10 min), sections were counterstained with Neurotrace 435/455 Blue Fluorescent Nissl Stain (1:100, ThermoFisher, cat# N21479) for 30 minutes at room temperature in a light-protected box (*Yi et al., 2014*). Following PBS washes (3 × 10 min), sections were then mounted with VECTASHIELD® Hardset™ Antifade Mounting Medium (Vector Laboratories, cat# H-1400), dried overnight, and sealed with clear nail polish. Confocal images were acquired using a T1-E microscope with A1 confocal (Nikon Instruments) at 10x, 40x, and 100x magnification.

### Criteria for selection of OLM cells

Patch clamp recordings were performed on a total of 45 cells from the stratum oriens of YFP-positive cells from heterozygous Chrna2-CRE+/−:td-Tomato+/− mice. After histological processing was complete, neurons were classified as OLM cells if they possessed a horizontally-oriented cell body and dendrites within the oriens layer and a major axon projecting perpendicularly with ramifications in the lacunosum/moleculare layer. Additional criteria were developed for stability of access and input resistance, completeness of electrophysiological protocols, and signal-to-noise level in both current and voltage clamp recordings. Only those cells that exhibited <20% change in input resistance over the course of the experiment were considered for further modelling. The full suite of electrophysiological protocols, including wash-in of ZD7288 blocker to be able to determine *I*_*h*_ currents, was required to fulfill selection criteria. Of the 45 cells recorded from in total, 11 OLM cells met electrophysiological criteria for stability, completeness, and noise level. An analysis of these 11 cells is given in *Table 1* in which the experimental data analysis was performed as in (*Yi et al., 2014*)).

Of these 11 OLM cells, three (***Cell 1***, ***Cell 2***, ***Cell 3***) were advanced for subsequent detailed experimental analyses and multi-compartment computational model development. Over the course of the recordings, the input resistances as determined from seal test recordings changed from: 260.5 to 216.9 *M* Ω (−16.7%) for *Cell 1*; 147.3 to 175.1 *M* Ω (+15.8%) for *Cell 2*; and 458.7 to 390.6 *M* Ω (−8.7%) for *Cell 3*. The sources of these modest changes in input resistance are not clear, but mechanical drift, activity-dependence (execution of many protocols), and intracellular dialysis are suspected to be contributing factors.

### Fitting passive properties in multi-compartment models

Simulations of multi-compartment models of neurons were performed using the NEURON simulation environment (*Hines and Carnevale, 2001*). We selected long current clamp steps for the fitting of passive membrane properties rather than shorter voltage clamp “seal test” protocols due to the incomplete clamping of the membrane by short voltage clamp steps (*Holmes, 2010*). Furthermore, these voltage traces minimize the contribution of active conductances. Recordings were performed with synaptic- and voltage-gated channels blocked, and was initially preferable for passive membrane property fitting in the models. Recordings obtained in the presence of h-channel blocker ZD7288 are referred to as *“ZD traces”* and are given by #7 in *Table 5*. Due to the possibility of changes in membrane responses as a function of the length of the recording session, we compared the membrane time constants (*τ*_*m*_) during the charging portion of the current clamp step for the voltage traces obtained across recordings with synaptic- and voltage-gated blockers applied. We found that the −30pA ZD traces, being the last traces recorded in the session, showed noisier membrane responses compared to the −120pA ZD traces obtained earlier. This manifested as an “undershoot” of the −30pA ZD traces after normalization of the traces was done, so that the −30pA ZD traces showed a marked slowing of *τ*_*m*_ compared to both the −120pA ZD as well as −30pA *“TTX traces”* (i.e., #4 in *Table 5*, referred to as such due to TTX application, in addition to potassium and synaptic blockers), the latter two being largely overlapping (*Figure 2-Figure Supplement 1A*). The noisier charging portion of the −30pA ZD traces could be seen more clearly if the time point of normalization of the traces occurred later, at 1000ms after the onset of the hyperpolarization current clamp step (*Figure 2-Figure Supplement 1A,b*). This demonstrated that in the case of the −30pA TTX current injection, few or no h-channels were activated as the *V*_*m*_ response was nearly identical to that of the ZD traces. This could be further confirmed quantitatively by fitting single exponential equations to the *V*_*m*_ responses from the time point of the onset of the hyperpolarization current clamp step (1000ms) to the point at which the steady-state of the *V*_*m*_ response was approximately achieved (*Figure 2*). For all cells, *τ*_*m*_ for the −30pA TTX trace was most closely matched by the −120pA ZD trace (dashed red line), with the subsequent ZD traces (−90pA, −60pA, and −30pA) exhibiting an increased, and hence slower, membrane response (*Figure 2*, bottom panels). The fact that the −120pA ZD trace exhibited a similar response as a current injection of one quarter magnitude and without h-channels being blocked indicated that in both cases, the response of the membrane was mostly passive.

Thus, given the lower signal-to-noise of the −30pA ZD traces, we considered that the passive properties obtained using the −120pA ZD traces would be better representations of the electrotonic features of the experimental cells. We thus fitted the passive membrane properties of multi-compartment models using a virtual current clamp and the Multiple Run Fitter (MRF) of the NEURON simulation environment (*Hines and Carnevale, 2001*). The −120pA ZD traces for each cell were used as the experimental recording for which the models’ *V*_*m*_ trajectories needed to match in response to −120pA virtual current. To confirm that the −120pA ZD traces led to better fits of the cells’ passive properties, we compared the fits obtained using −120pA and −30pA ZD traces. The resulting fitted passive parameters of axial resistivity (*R*_*a*_), specific capacitance (*C*_*m*_), leak conductance (*G*_*pas*_) and leak reversal potential (*E*_*pas*_) are displayed in *Table 6*. For each cell, the cumulative root-mean-square error (RMSE) across all traces used for each fit was lower when the −120pA ZD trace was used for fitting the passive properties (*Table 6*, left column for each cell).

During use of the MRF in NEURON for the passive property fitting procedure, certain regions of the traces were discounted from fitting, such as the first 500ms portion so that initial model transients did not affect the fitting. Furthermore, because the charging portion of *V*_*m*_ was very short – on the order of 100ms – it was given a greater weight value (10X) compared to the rest of the trace, in the MRF. Only the initial portion of the steady-state response after the hyperpolarizing current clamp step was used for fitting. This was because for some cells, a small depolarization was present even under ZD7288 block, which could have been due to noise or the presence of another, unidentified inward current that was not blocked.

**Table 6.** Fitted passive parameters and resulting goodness of fit of models to various current clamp step traces.

|  | **Cell 1** | **Cell 1** | **Cell 2** | **Cell 2** | **Cell 3** | **Cell 3** |
| --- | --- | --- | --- | --- | --- | --- |
| *"ZD trace"* |  |  |  |  |  |  |
| used for fitting | -120 pA | -30 pA | -120 pA | -30 pA | -120 pA | -30 pA |
| $R_a$ ($\Omega\text{cm}$) | 141.85 | 258.21 | 285.78 | 474.78 | 94.83 | 329.99 |
| $C_m$ ($\mu\text{F}/\text{cm}^2$) | 0.2698 | 0.2552 | 0.2799 | 0.4721 | 0.3057 | 0.4744 |
| $G_{pas}$ ($\text{S}/\text{cm}^2$) | $7.93 \times 10^{-6}$ | $9.62 \times 10^{-6}$ | $9.24 \times 10^{-6}$ | $9.39 \times 10^{-6}$ | $7.48 \times 10^{-6}$ | $7.27 \times 10^{-6}$ |
| $E_{pas}$ (mV) | -49.0 | -48.3 | -54.7 | -56.6 | -69.1 | -68.1 |
| RMSE (-30 pA ZD) | 0.5058 | 0.3289 | 0.6594 | 0.1893 | 1.079 | 0.0456 |
| RMSE (-60 pA ZD) | 7.458 | 5.742 | 0.9828 | 0.7092 | 2.410 | 1.398 |
| RMSE (-90 pA ZD) | 1.945 | 3.393 | 0.6155 | 3.829 | 2.194 | 16.89 |
| RMSE (-120 pA ZD) | 0.3602 | 3.440 | 0.9818 | 9.875 | 0.5199 | 50.38 |
| RMSE (-30 pA TTX) | 0.6596 | 0.2048 | 0.3108 | 0.6234 | 1.679 | 9.021 |
| Cumul. RMSE (mV) | 10.92 | 13.10 | 3.550 | 15.22 | 7.881 | 77.73 |

### Passive membrane model and experiment comparisons

Input resistance (*R*_*in*_) in the passive models computed using a current clamp protocol of −120 pA, i.e., the same protocol used to fit the passive properties, is given by values of *V*_*m*_ taken at the start and end of the current clamp step: *R*_*in*_ = (*V*_*start*_ − *V*_*end*_)/(120*pA*). Using experimental −120pA ZD traces, the input resistance is also computed. These input resistance values are shown in *Table 7*.

**Table 7.** Computed surface areas, model discretizations, input resistances and time constants.

| **Parameter** | **Cell 1** | **Cell 2** | **Cell 3** |
| --- | --- | --- | --- |
| Soma surface area ($\mu\text{m}^2$) | 7,651 | 13,035 | 6,911 |
| Somatodendritic surface area ($\mu\text{m}^2$) | 29,378 | 35,159 | 21,990 |
| Number of compartments (model with $I_h$) | 303 | 632 | 837 |
| $R_{in}$ ($\text{M}\Omega$) (passive model) | 411 | 332 | 550 |
| $R_{in}$ ($\text{M}\Omega$) (expt with ZD7288) | 363 | 326 | 531 |
| $\tau_m$ (ms) (passive model) | 30.4 | 27.4 | 36.4 |
| $\tau_m$ (ms) (using -30pA TTX trace) | 32.8 | 29.1 | 40.5 |
| $\tau_m$ (ms) (using -120pA ZD trace) | 32.3 | 33.8 | 39.3 |

We note that for the comparison of membrane time constants of the OLM cells used, we fitted exponential curves to the charging portion of *V*_*m*_ for each cell at various time points of the recording session using a nonlinear least squares regression (*Figure 2*). The amplitude of the traces were normalized at the time point at which depolarizing responses in the *“TTX traces”* (i.e., #4 in *Table 5* when TTX/4-AP/TEA applied), due to the h-channel current (*I*_*h*_) cause the membrane potential to deviate from the (putatively) passive response under the *“ZD traces”* (i.e., #7 in *Table 5* when ZD7288 also applied). For each cell, both the −30pA and −120pA ZD traces were used to compare to the TTX traces, as these should both reflect largely passive membrane responses. We note that for most cells, the −30pA TTX trace followed the −30pA ZD trace (left) as well as the −120pA ZD trace (right), *Figure 2*. *Cell 1* in particular exhibited a very good match between the −30pA TTX and −120pA ZD traces. The fitted *τ*_*m*_ values for the −120pA ZD trace and −30pA TTX trace are given in *Table 7*. We also fitted the membrane time constant for the models, using a −120pA current clamp step in the models without *I*_*h*_ included (see *Table 7*). Resulting *V*_*m*_ traces were 1t in the same way as the experimental traces, except that the *V*_*m*_ data points were weighted by the relative time step of integration in the NEURON simulations such that data points in the *V*_*m*_ vector closely spaced in time would be weighed less. This ensured that the 1t was not disproportionately weighed by the early, rapidly changing charging portion with many more data points.

Compartmentalization of the models was done in NEURON using the *d*_*λ*_ rule where compartment lengths are set to a fraction of the length constant *λ*_*f*_, where *f*=100Hz. We set the fraction of *d*_*λ*_ to be 0.1 for all models. *Table 7* gives the resulting number of compartments in each of the cells, along with their surface areas in the finalized models, that is, after staggered re-fitting.

### Mathematical equations for h-channels

The specification of the current for h-channels, *I*_*h*_, was taken from our previous work (Lawrence et al., 2006b; *Sekulić et al., 2014*). However, the kinetics for activation and deactivation, the steady-state activation curves, and the conductance densities were defined on a per-cell basis in the present work (see Results). This required moving the relevant variables in the *I*_*h*_ MOD-file into the PARAMETER block to allow per-cell configuration in the NEURON code.

The conductance-based mathematical formulation used to represent current flow through h-channels is given by:

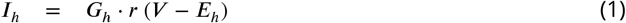

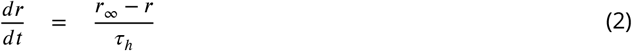

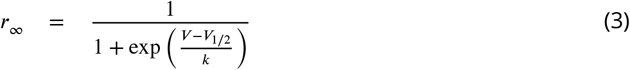

where *G*_*h*_ is the maximal synaptic conductance for the h-channels, *r* is the activation variable, *E*_*h*_ is the h-channel reversal potential, *r*_∞_ is the steady-state activation, *k* is the slope of activation and *V*_1/2_ is the potential of half-maximal activation of *I*_*h*_, *τ*_*h*_ is the time constant of activation, *V* is the membrane potential, and *t* is time. The voltage dependence of *τ*_*h*_ is given by a double exponential expression with parameters *t*_1_, *t*_2_, *t*_3_, *t*_4_, *t*_5_ as follows:

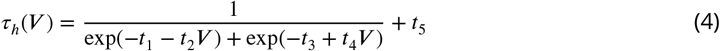

### Extraction of h-channel characteristics from voltage and current clamp traces

Given our experimental protocol (see *Table 5*), we were able to obtain h-channel current (*I*_*h*_) reversal potentials, activation kinetics, and steady-state activation for each of the three chosen cells.

#### Reversal potential

To obtain the reversal potential for *I*_*h*_, we first removed the leak components and capacitive transients from the voltage clamp recordings in order to isolate the *I*_*h*_ components. This was done by taking the traces obtained by the reversal potential protocol (#6 in *Table 5*) and subtracting from them the capacitive response generated by an equivalent magnitude voltage clamp deflection from the *I*_*h*_ activation protocol with ZD7288 application (#7 in *Table 5*), resulting in *I*_*h*_ tail currents *Figure 3-Figure Supplement 1A*). The traces were then smoothed using the *rloess* smoothing function in MATLAB, which performs local linear regression over a window, using weighted linear least squares. The smoothing window was set to 25ms so that only noise in the recordings was removed, and not time-dependent changes attributable to ion channel currents.

To create the current-voltage (I-V) plot, a fixed time point after the capacitive transient ended was determined by eye, which allowed us to obtain the time point of maximum deflection after the voltage clamp step (*Magee, 1998*; *Molleman, 2003*). We refer to this as the “fixed” time point for determining the I-V plot. The validity of this technique relies on the assumption that the maximal number of h-channels are still open by the time the capacitive transient is abolished, so that the resulting current does not depend on changes in the conductance, only on the driving force. The fact that *I*_*h*_ deactivates slowly means that this assumption is likely to be a safe one. However, to account for the possibility of early channel closure, a second method for extracting the current values and constructing an I-V plot was used. This consisted of fitting single exponential functions to the time course of the decay of current upon the step relaxation of the voltage clamp, which is used primarily to determine the voltage-dependent time constants of deactivation. The fitted exponential functions were then evaluated at the time of the relaxation of the voltage clamp step. In this way, we could deduce the amount of current that is masked by the capacitive transient by extrapolating the value from the exponential functions that were fitted on the non-capacitive portions of the current trace. That is, the functions were fitted to a window corresponding to the fixed point as the start time, and the end of the voltage clamp step as the end time. We refer to the current values and resulting I-V plot as the “extrapolated” method. We note that the smoothed traces were only used for the fixed method so that noisy fluctuations in the current traces did not unduly influence the resulting I-V plot; however, for the extrapolated method the exponential functions were fitted using the original, non-smoothed subtracted traces. The current traces with fitted exponentials are shown in *Figure 3-Figure Supplement 1A*, and I-V plots for both fixed and extrapolated methods are shown in *Figure 3A*. The resulting reversal potential (*E*_*h*_) values for each cell were determined by fitting a first-order polynomial to the linear portion of the I-V curve only. For *Cell 1* and *Cell 2*, the linear portion of the extrapolated I-V curve overlapped with the fixed I-V curve, and the resulting *E*_*h*_ values were similar between the two methods. For these cells, we therefore took *E*_*h*_ from the extrapolated I-V curves. For *Cell 3*, however, the capacitive transients disrupted the response and affected the fitting so that the extrapolated I-V values did not exhibit as strong of a linear relationship as the fixed I-V values. One possible explanation for the distorted (non-linear) measurements of I-V values with *Cell 3* is that the current traces for *I*_*h*_ deactivation, from which the reversal potential I-V plots were determined, did not match fully between the control case (with only TTX/4-AP/TEA blockers) and the later protocol with the *I*_*h*_ blocker ZD7288, due to the effects of noise. Thus, the subtraction of the two to remove the leak components introduced some distortion in the resulting current traces. As a result, although the resulting *E*_*h*_ values between extrapolated and fixed points were similar, we took *E*_*h*_ for *Cell 3* from the fixed I-V curve instead, to minimize possible error from using the line fitted with only 4 out of the 8 possible I-V datapoints (*Figure 3A*, *Cell 3*). The resulting *E*_*h*_ values for all cells are given in *Table 8*. These values are in general agreement with literature values of *I*_*h*_ reversal potentials in OLM cells (*Maccaferri and McBain, 1996*).

#### Voltage-dependent time constant of activation and deactivation and steady-state activation

To obtain the time constants of activation/deactivation for *I*_*h*_ (*τ*_*h*_) we used the recordings where a voltage clamp protocol with an initial clamp at a holding potential was then stepped to various hyperpolarized potentials, measuring the resulting transmembrane current (#3 in *Table 5*). The identical protocol was then performed with the *I*_*h*_-specific blocker ZD7288 (#8 in *Table 5*). Using this data, we subtracted the ZD7288 traces from the control traces to isolate *I*_*h*_. Then, single exponential functions were fitted to the time-varying change in current upon each voltage step. *I*_*h*_ showed no voltage-dependent inactivation (*Figure 3-Figure Supplement 1B*). To construct the curve of voltage-dependent activation and deactivation kinetics, the time constants of activation were combined with the deactivation time constants obtained from the tail currents (*Figure 3-Figure Supplement 1A*). The time course of activation and deactivation was then described using a double exponential function of the form given by Equation (4) in the Methods, with parameters *t*_1_, *t*_2_, *t*_3_, *t*_4_, *t*_5_ to be fit. Fitting of the double exponential functions was done using the Curve Fitting toolbox in MATLAB. The resulting fitted values for the voltage-dependent time constant of activation and deactivation are given in *Table 8* and plotted in *Figure 3B*. We note that the shape of the time constant of activation function is roughly similar across the three cells, with particular overlap between *Cell 1* and *Cell 2*. In all three cases, the slowest component of the time constant activation function is around 300 ms, whereas the fast component is less than 100 ms for all three cells.

**Table 8.** *I*_*h*_ parameter values obtained from fits to experimental data of each cell and computed conductance densities for somatic or somatodendritic distributions.

| **Parameter** | **Cell 1** | **Cell 2** | **Cell 3** |
| --- | --- | --- | --- |
| $E_h$ (mV) | -34.0 | -27.9 | -25.2 |
| $V_{1/2}$ (mV) | -103.4 | -100.1 | -111.3 |
| $k$ (slope factor) | 8.63 | 11.16 | 6.88 |
| $t_1$ (ms) | 8.03 | 8.98 | 35.09 |
| $t_2$ (ms) | 0.025 | 0.035 | 0.24 |
| $t_3$ (ms) | -4.40 | -8.49 | -4.28 |
| $t_4$ (ms) | 0.15 | 0.19 | 0.088 |
| $t_5$ (ms) | $7.32 \times 10^{-6}$ | $3.57 \times 10^{-7}$ | 69.72 |
| Total $G_h$ (nS) | 4.17 | 3.64 | 2.20 |
| Soma surface area ($\mu\text{m}^2$) | 7,650.9 | 13,034.5 | 6,910.5 |
| Somatodendritic surface area ($\mu\text{m}^2$) | 29,378.1 | 35,158.5 | 21,990.3 |
| $G_h$ (pS/$\mu\text{m}^2$), $H_{dist}=0$ | 0.546 | 0.279 | 0.380 |
| $G_h$ (pS/$\mu\text{m}^2$), $H_{dist}=1$ | 0.142 | 0.104 | 0.120 |

#### Steady-state activation

The steady-state activation curves, *r*_∞_, for the OLM cells were constructed by measuring the current amplitude in the ZD7288-subtracted traces at the end of each step of the voltage clamp protocol for *I*_*h*_ activation (*Figure 3-Figure Supplement 1B*). The current at each voltage step was plotted and normalized to the greatest recorded current value which for h-channels is at the most hyperpolarized range. Then, a Boltzmann function for *r*_∞_ (Equation (3) in Methods) with parameters *V*_1/2_ for the voltage at half-activation and slope factor *k* for the steepness of the sigmoidal curve, was fitted to each cell’s voltage-dependent activation data. The fitted values are given in *Table 8*, and the resulting activation curves for the three cells are shown in *Figure 3C*.

#### Maximal conductances

To determine the maximal conductance for *I*_*h*_, *G*_*h*_, we used the tail currents from the reversal potential step protocol as this corresponded to the point in time when *I*_*h*_ was fully activated (*Dougherty et al., 2013*; *Magee, 1998*). These currents were thus measured when all h-channels are opened, and thus describe the ratio of maximum current to voltage needed to obtain I-V plots for determining *G*_*h*_ (*Molleman, 2003*). The slope of the linear portion of the I-V plot for the tail currents, with the reversal potential as origin (denoting zero current flow), was used as the measure of *G*_*h*_. As described above, a line was fitted to the linear portion of the I-V plots for all three cells to determine the reversal potential. The slope of the line gives *G*_*h*_ and the resulting values for the three cells are given in *Table 8*. When scaled by the surface area, we obtain an *G*_*h*_ as a conductance density that is used in the model code.

### Staggered model re-fitting considerations with h-channels

In consideration of a re-fitting procedure, we noted that if we wanted to judiciously tune individual parameters in such a manner that for a given portion of each *V*_*m*_ trace, the parameters that can be responsible for affecting that portion of the trace should be tuned in the order from those with the greatest uncertainty to those with the least uncertainty. Using as an example the (passive property) scenario of the initial charging portion of the membrane upon step hyperpolarizing current, although we would expect all of *C*_*m*_, *R*_*a*_, *τ*_*h*_ and *r*_∞_ to affect this portion of the trace to various degrees, it would have been a mistake to fit, say, *r*_∞_ prior to fitting *C*_*m*_. This is not only because *I*_*h*_ wouldn’t yet be fully activated but also because the fitted function for *r*_∞_ exhibits a very good match to the recorded steady state current values for multiple voltage steps (*Figure 3C*), whereas the fit for *C*_*m*_ has more uncertainty due to the issues of dendritic diameter estimation as well as cell rundown seen in the recordings used for fitting, as previously described. If we were to have fitted *r*_∞_ first, we would have attributed an undue source of error of the mismatch in *V*_*m*_ to *r*_∞_. The disproportionate change in *r*_∞_ curve would then have manifested in inappropriate *V*_*m*_ output elsewhere. Therefore, the criterion for whether we had selected an appropriate parameter for re-fitting was that if the re-fitted parameter resulted in a better fit to the portion of *V*_*m*_ trace under consideration but a worse fit elsewhere, then we had not selected the correct parameter for which the error in *V*_*m*_ mismatch should be attributed to. In practice we could perform this test by fitting to one set of current clamp traces and validating the model’s correctness by testing its output to another current clamp step trace without re-fitting the parameter.

The approach of inappropriately attributing errors to parameters, taken to its extreme, would be to allow all parameters to be adjusted simultaneously. We demonstrate the results of this “naïve” scheme by allowing the passive properties (4 parameters) and *I*_*h*_ properties (9 parameters) to simultaneously vary while using the PRAXIS fitting procedure in NEURON to minimize the error in *V*_*m*_ response between model and experiment. Let us use *Cell 3* as an example. When first fitted to the −120pA current clamp step TTX trace, the model exhibited a remarkably good 1t to the experimental *V*_*m*_ trace (*Figure 4-Figure Supplement 1A*, left) which, at first glance, would indicate that the OLM cell’s output had been captured. However, when we then injected a −90pA current clamp step in the model with these fitted parameters as a test and compared its output to that of the experimental cell, we saw that it was very poor at matching the −90pA TTX current trace response (*Figure 4-Figure Supplement 1A*, right). If we instead fitted to the −90pA current clamp TTX trace, we found that the model could also match the experimental *V*_*m*_ output quite well (*Figure 4-Figure Supplement 1B*, right) but then it failed to capture the −120pA output when used as a test (*Figure 4-Figure Supplement 1B*, left). It is important to note that in doing the fits and tests, we ensured that the holding current applied to the model was always in line with what was used for the particular cell at the given current step, although fitted parameters were not changed between fits and tests. This “overfitting” of the experimental data used for adjusting the parameters was thus inappropriate, and a more judicious procedure was required, where only a subset of parameters were considered at any given time and for any given feature of the *V*_*m*_ mismatch between model and experiment.

### Full spiking multi-compartment model optimizations

In creating full spiking models, we used the final passive model backbone with h-channels in the dendrites, and used the same complement of ion channel types that had been used in previous instantiations of the OLM cell model (Lawrence et al., 2006b; *Sekulić et al., 2014*). The equations used are all given in the Appendix of Lawrence et al. (2006b). They include transient sodium, fast and slow delayed rectifier potassium, A-type potassium, M-type, T- and L-type calcium, and calcium-dependent potassium channels. Their conductances in soma (*s*), axon (*a*) or dendrites (*d*) are represented respectively as *G*_*NaT*_, *G*_*Kdrf*_, *G*_*Kdrs*_, *G*_*KA*_, *G*_*M*_, *G*_*CaT*_, *G*_*CaL*_, *G*_*KCa*_ as given in *Table 9*.

In our optimizations, we allowed *G*_*NaT*_, *G*_*Kdrf*_, *G*_*Kdrs*_ to vary independently in the soma, dendrites, and axon, and we also allowed the sodium channel to have some flexibility by allowing alterations in its voltage dependency, i.e., introducing a free parameter, *V*_*shift*_ that could change by ± 7 mV. Note that soma, dendrites, and axon each have an independent *V*_*shift*_ parameter, but the *V*_*shift*_ value remains the same across forward and backward rate activations and inactivations such that activation and inactivation curves shift by the same amount and the “activation/inactivation window” stays constant. Except for the inclusion of *V*_*shift*_, the activation and inactivation equations underlying the sodium current are the same as used previously (Lawrence et al., 2006b), and as based on experimental data of *Martina et al. (2000)*. For completeness, the equations for the sodium current, *I*_*NaT*_, are shown below:

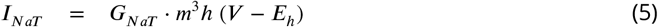

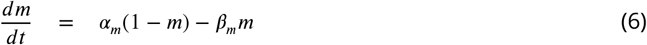

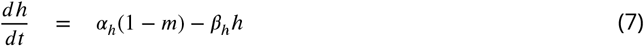

where, for somatic compartments,

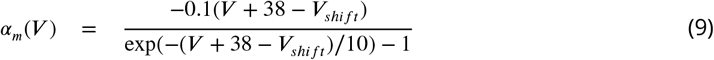

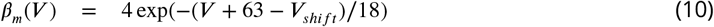

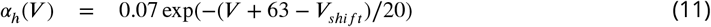

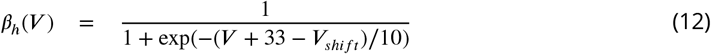

and for dendritic and axonal compartments,

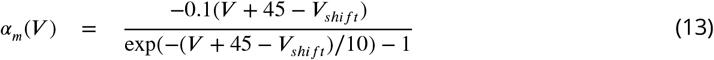

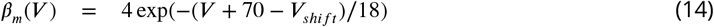

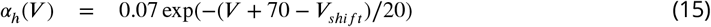

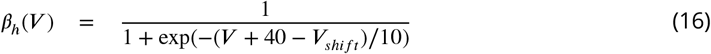

**Table 9.** Location and optimization ranges for ion channel types.

| **Conductance type** | **Distribution location** | **Cell 1 range (pS/$\mu\text{m}^2$)** | **Cell 2 range (pS/$\mu\text{m}^2$)** |
| --- | --- | --- | --- |
| $G_{NaT,s}$ | soma | 10-100 | 10-100 |
| $G_{NaT,d}$ | dendrites | 40-200 | 40-200 |
| $G_{NaT,a}$ | axon | 40-200 | 40-200 |
| $G_{Kdrf,s}$ | soma | 3-200 | 50-200 |
| $G_{Kdrf,d}$ | dendrites | 3-200 | 50-200 |
| $G_{Kdrf,a}$ | axon | 3-200 | 50-200 |
| $G_{Kdrs,s}$ | soma | 0-0.01 | 0-0.01 |
| $G_{Kdrs,d}$ | dendrites | 0-0.01 | 0-0.01 |
| $G_{Kdrs,a}$ | axon | 0-0.01 | 0-0.01 |
| $G_{KA}$ | soma, dendrites | 1.25-120 | 1.25-120 |
| $G_M$ | soma, dendrites | 0.05-1.5 | 0.05-1.5 |
| $G_{CaT}$ | dendrites | 0.625-5 | 0.625-5 |
| $G_{CaL}$ | dendrites | 6.25-50 | 6.25-50 |
| $G_{KCa}$ | dendrites | 1.375-11 | 1.375-11 |

### Optimization Approach and Parameter Details

For the optimizations, we did the following:

1. Performed multi-objective optimizations using the BluePyOpt module in Python (*Van Geit et al., 2016*) and high performance computing resources via the Neuroscience Gateway (*Sivagnanam et al., 2013*) to find ion channel conductances in order to minimize the error across multiple features in the electrophysiology - see *Table 10*.
2. Fine-tuned the parameter ranges and objectives to avoid areas of the parameter space that generate undesirable results and keep re-doing the optimizations using this approach until the top models consistently generate appropriate electrophysiologies. The parameter ranges used that produced the 1nal models are shown in *Table 9*.

**Table 10.**
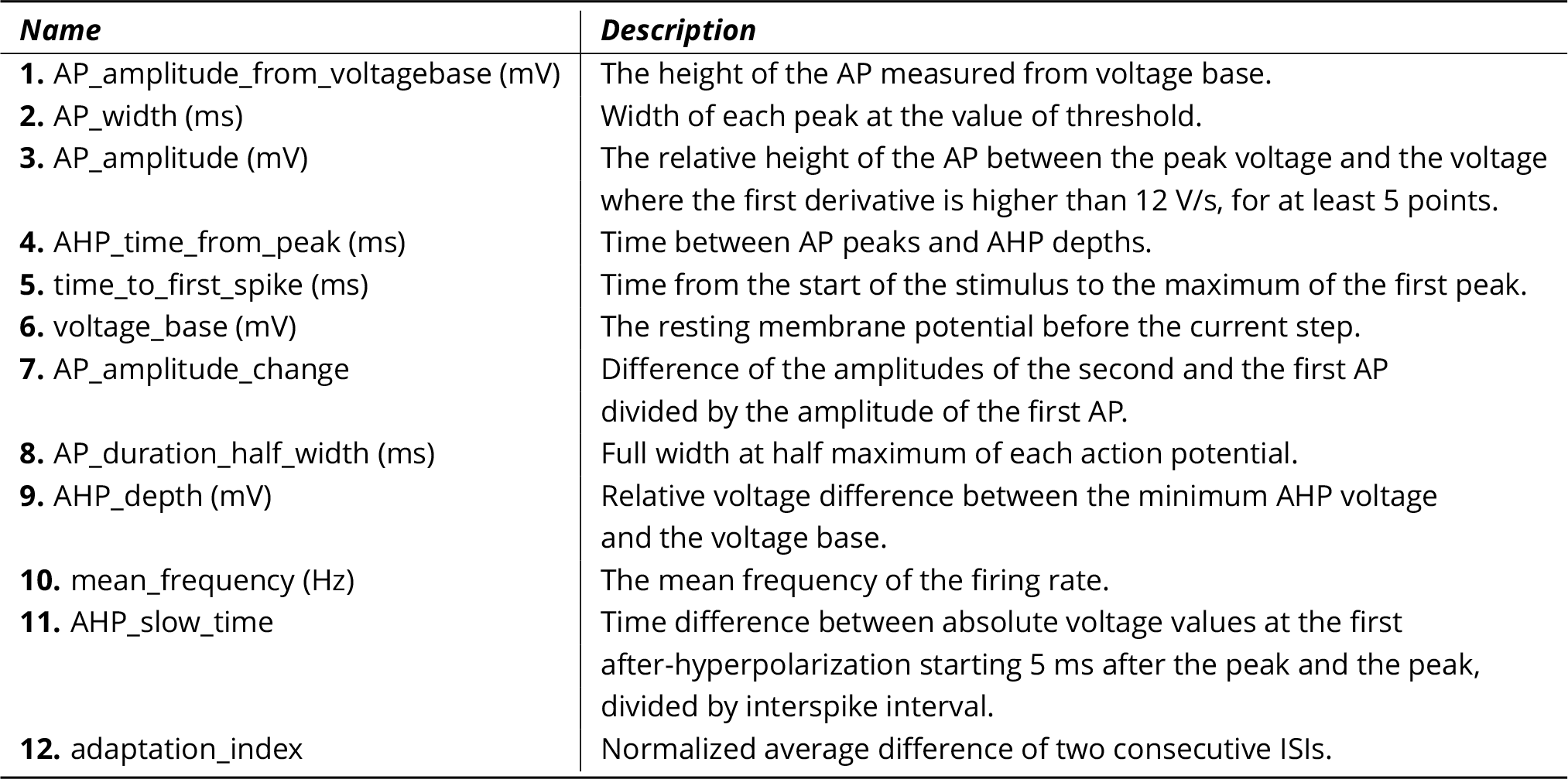
Descriptions of the eFEL measurements that were used as objective features in the spiking model optimizations. AP: Action Potential; AHP: After-Spike Hyperpolarization.Consult the eFEL manual for more details on these measurements: https://media.readthedocs.org/pdf/efel/latest/efel.pdf

The top five optimized models for *Cell 1* and *Cell 2* are presented in *Figure 7* and *Figure 7, Figure Supplement 3*. Their fitness values are: (*Cell 1:* 411.53, 425.38, 430.68, 430.93, 438.65; *Cell 2:* 660.96, 665.49, 669.06, 678.58, 684.60.).

In order of model rankings (i.e. [1^*st*^,…,5^*th*^]), the values below are the *V*_*shift*_ parameters (in mV) for the top five full spiking models (see *Figure 7, Figure Supplement 4C* to see the resulting voltage-dependencies).

*Cell 1: V*_*shift,s*_ = [−4.83, −6.55, −6.70, −6.46, −6.68], *V*_*shift,d*_ = [4.85, 6.37, 3.71, 2.86, 4.17], *V*_*shift,a*_ = [2.49, 2.82, 2.84, 5.70, 2.82].

*Cell 2: V*_*shift,s*_ = [−4.36, −4.69, −4.88, −4.36, −4.36], *V*_*shift,d*_ = [−1.26, −1.12, −0.54, −1.46, −1.41], *V*_*shift,a*_ = [6.57, 6.57, 6.30, 6.57, 6.57].

After performing several optimizations and adjusting the parameters to improve the optimization outputs, we used the following optimization parameters for both models: Number of Offspring = 100, Number of Generations = 200, Mutation Rate = 0.15, Crossover Rate = 0.85, Eta (i.e. learning rate) = 0.5, Optimizer = ‘IBEA’, Random Seed = 61 (*Cell 1*) and 9 (*Cell 2*)

All of the objective features that were used in the optimization are listed in *Table 10*, and the parameter ranges are given in *Table 9*. Features 1-10 were used for the +30 pA, +60 pA, and +90 pA current injection protocols. Features 11-12 were only used for the +60 pA and +90 pA current injection protocols, since the +30 pA current injection did not always generate a sufficient number of spikes for those features to be calculated. Since we were fitting the models to single current injection traces, standard deviation values were chosen manually for each objective feature, in order to weight each objective feature by hand. Since standard deviation is used in computing the fitness for each model (i.e. fitness is quantified as the sum of standard deviations away from the experimental target efeature values), manipulating these values offered a way to weight particular target measurements. More specifically, we initially chose standard deviations that were 1-2 order of magnitudes smaller than the largest significant digit for each measurement. For example, AP_duration_half_width in the somatic area of a neuron is usually a small value between 0.5-2 ms, and we used a standard deviation of 0.01 ms for this efeature. If the optimization ended up under-performing on any specific efeature measurements, we would sometimes attempt to improve it by using smaller standard deviation values for those measurements. Though this had some mild effects on improving the optimizations, constraining the free parameter ranges ended up showing much better improvements in the optimization results. We also added a heavy penalization on models that generated spikes during the baseline periods. Finally, in order to make BluePyOpt compatible with the OLM cell model compartmentalization, we adjusted BluePyOpt’s method for compartmentalization such that it uses the *d*_*λ*_ rule.

To check if axonal properties were appropriate for what is known experimentally (*Martina et al., 2000*), we performed simulations with our final optimized spiking models of *Cell 1* and *Cell 2* to investigate morphological sites of action potential (AP) initiation. specifically, *Martina et al. (2000)* previously showed that depending on whether a short high-intensity current or a long low-intensity current was injected into the soma, an AP would occur initially in the soma or axon-*bearing* dendrite, respectively. For both models of *Cell 1* and *Cell 2*, short high-intensity current evoked action potential initiation in the soma, but long low-intensity current evoked action potential initiation in axon-*lacking* dendrites. This suggests that specialized distributions of spike-initiating channels are missing in the axon of the model and are necessary for correctly setting the action potential initiation site. Given that OLM cell axonal channel properties are unknown, we did not venture further into specializing axonal properties in our models.

## Code and data availability

NEURON code for all the models are available on https://github.com/FKSkinnerLab/OLMng and associated experimental data available on https://osf.io/qvnu9/.

## Acknowledgments

Work on the Chrna2-CRE:tdTomato mice was performed under an established Material Transfers Agreement between Uppsala University and Texas Tech University Health Sciences University under Dr. Klas Kullander. We thank Drs. Sanja Mikulovic and Klas Kullander for their help at early stages of immunocytochemical experiments. Preliminary studies of this work have appeared in the form of several published Society for Neuroscience poster abstracts. FKS thanks Dr. Scott Rich for an overall reading of this work.

---

**Figure 2–Figure supplement 1.**
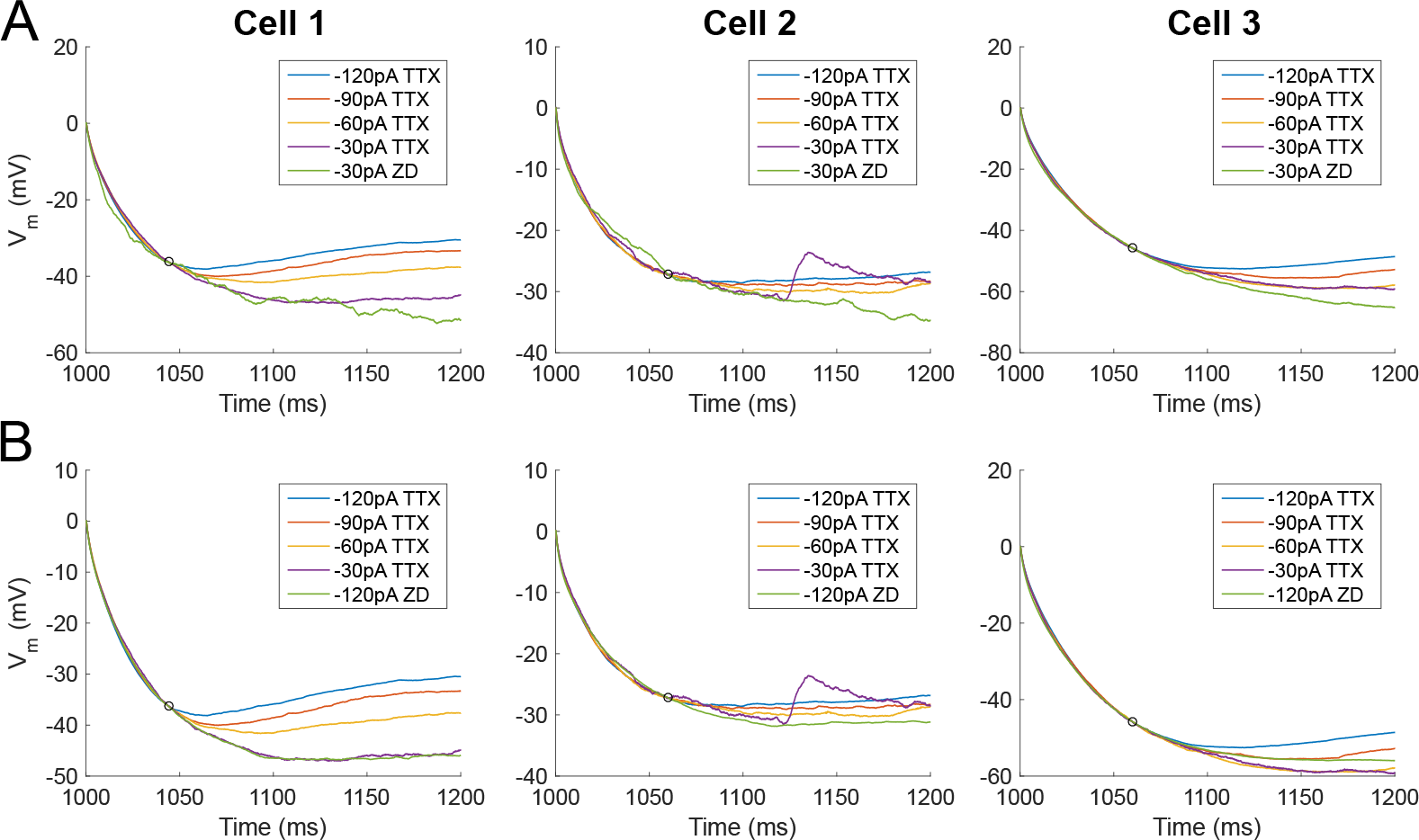
Comparison of charging portions of membrane potential (*V*_*m*_) for fitting model passive responses. **A.** Traces for all hyperpolarizing current injection steps with synaptic and voltage-gated channel blockers, except ZD7288 (“*TTX*”) and the −30pA trace with ZD7288 application (“*ZD*”) for *Cell 1, 2,* and *Cell 3* (respectively left, middle, right - ordering of cells are the same for remainder of figure). Small circles represent the time point at which all traces were normalized and were determined by eye as the point at which depolarization due to activted h-channels caused TTX traces to deviate from the “passive” ZD condition. This value is unique per cell and relative to the time of step current injection (1000ms) as follows: 44 ms (*Cell 1*), 60 ms (*Cell 2*), 60 ms (*Cell 3*). **B.** As in **A.**, except the −120pA ZD trace is shown instead of the −30pA ZD trace.

**Figure 2–Figure supplement 2.**
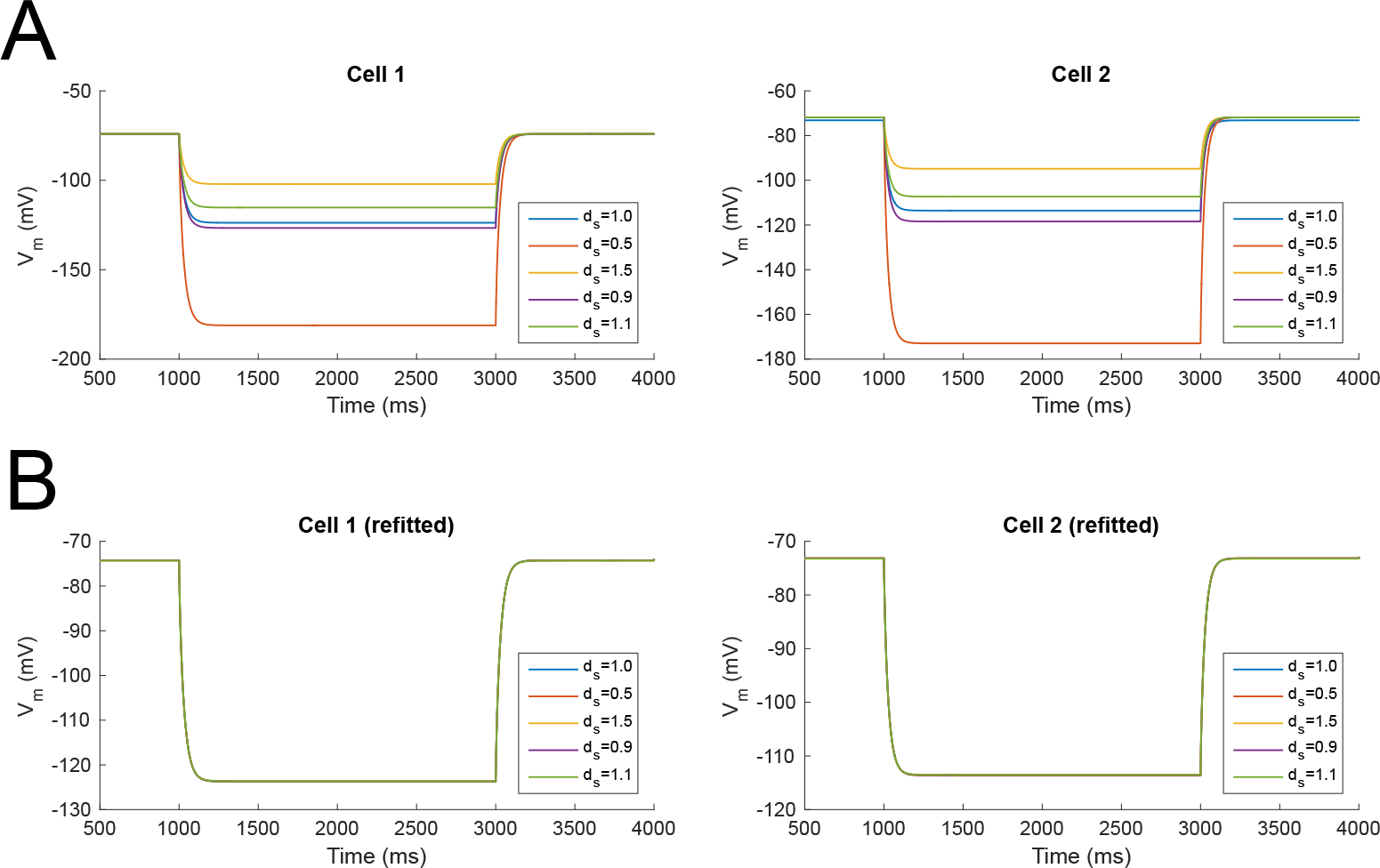
Model membrane potential (*V*_*m*_) responses with scaled dendritic diameters and both unchanged and refitted passive properties. Responses of two models (*Cell 1*, left and *Cell 2*, right) to −120pA current clamp step commands and a wide range of rescaled dendritic diameters and with **A.** original (fixed) and **B.** refitted passive properties. The scaling factor, *d*_*s*_, represents the constant scaling factor performed on each compartment’s diameter in the respective cell reconstruction. The response of the original reconstructed model – i.e., with no scaling – is shown as *d*_*s*_ =1.0 for all cases.

**Figure 3–Figure supplement 1.**
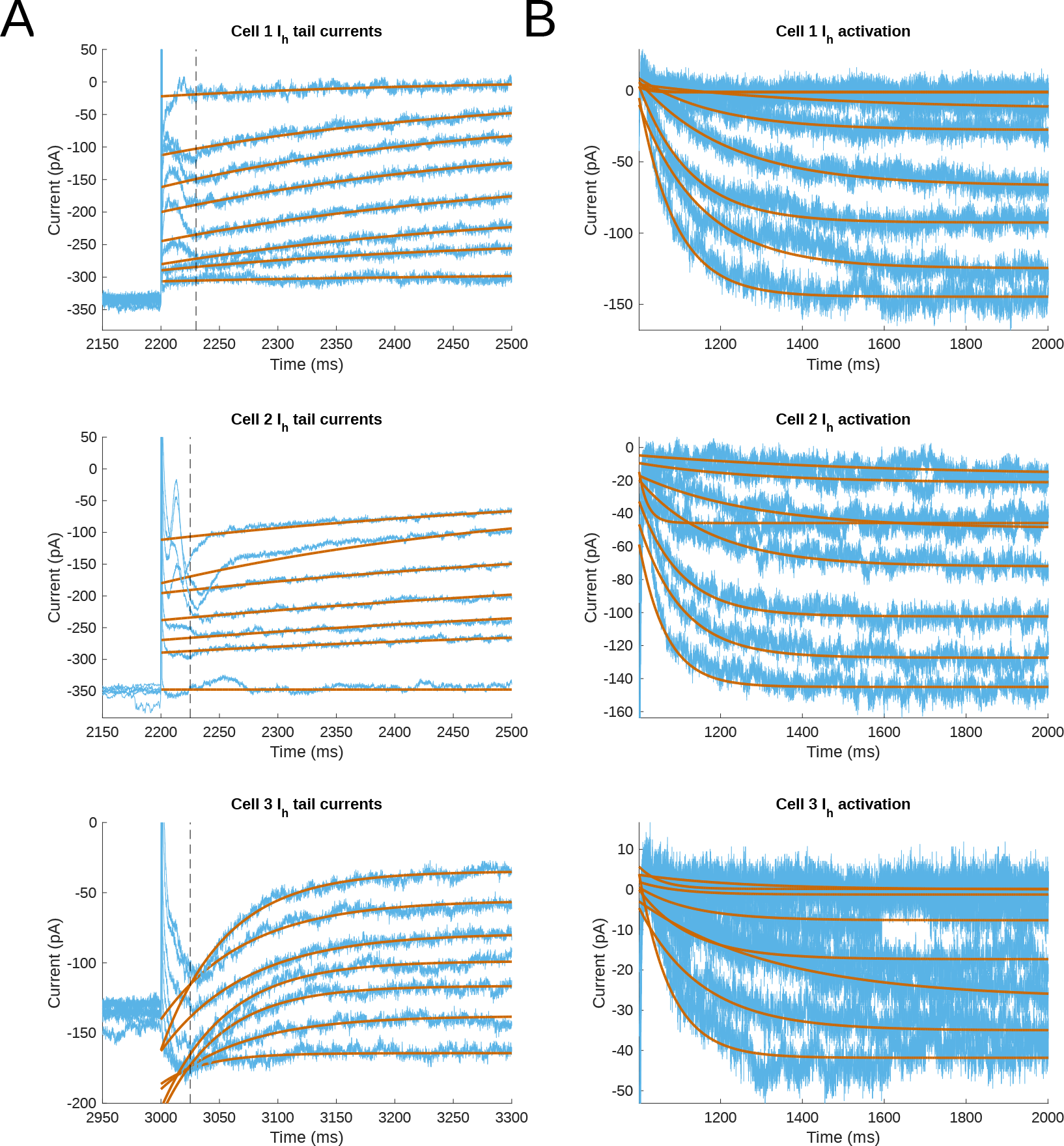
Experimental traces and fits for reversal potential and time constants of activation and deactivation. **A.** Tail current fits. **B.** *I*_*h*_ activation fits. Tail current protocol with leak subtraction, and ZD-subtracted traces from a voltage-clamp hyperpolarizing step protocol to reveal kinetics of *I*_*h*_ activation (B), for *Cell 1* (top), *Cell 2* (middle), and *Cell 3* (bottom). Fitted single exponential functions for each trace are shown in the tail current plots (A). The vertical dashed lines denote the approximate point of termination of the capacitive transient, and hence maximum deflection of the *I*_*h*_ current after the relaxation step in the voltage clamp protocol.

**Figure 4–Figure supplement 1.**
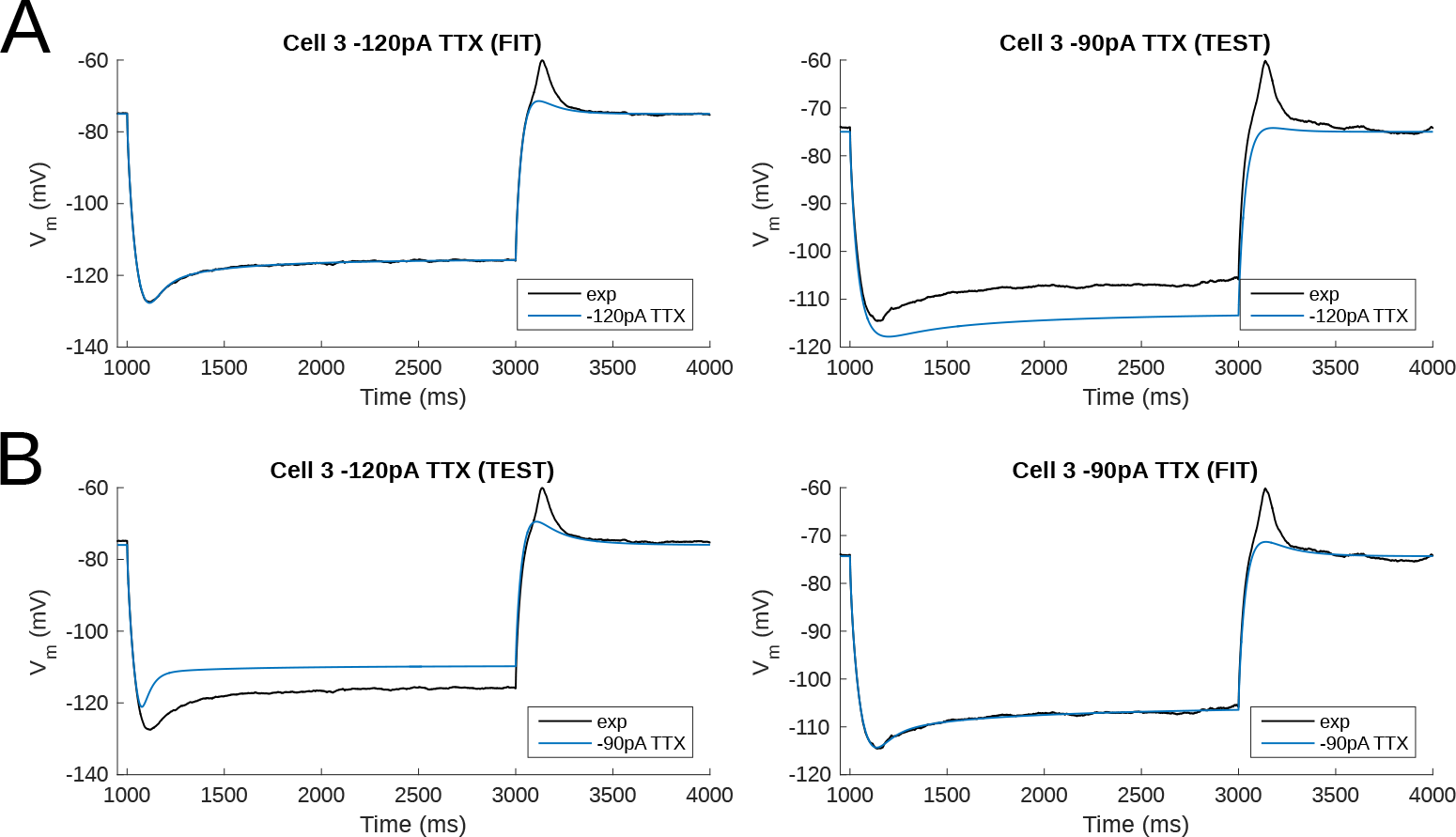
Simultaneous fitting of all passive and *I*_*h*_ parameters leads to overfitting of the experimental traces and poor model generalization - Cell 3 as an example. **A.** Model fitted to a −120pA TTX current clamp trace (left) and tested against a −90pA current clamp trace (right). **B.** The reverse case, with model fitted to a −90pA TTX current clamp trace (right) but tested against a −120pA trace (left). *H*_*dist*_=1. Holding current injections: 2.7 pA for −120pA step; 3.1 pA for −90pA step.

**Figure 4–Figure supplement 2.**
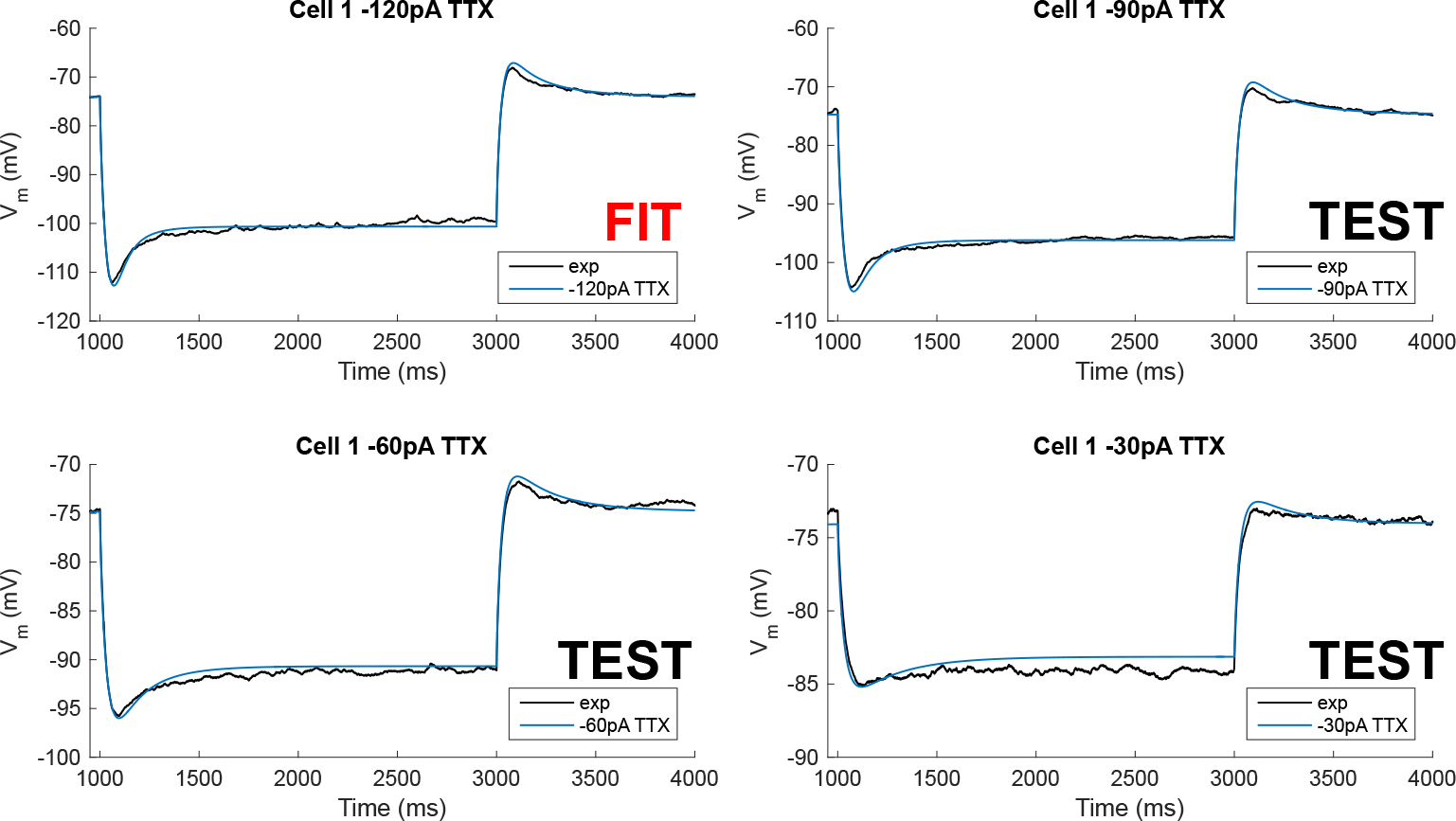
Traces for Cell 1 model compared to experiment after staggered re-fitting. Model *V*_*m*_ traces compared to experiment for *Cell 1* with staggered re-fitting procedure, where first passive properties are fitted, followed by total *G*_*h*_, *r*_∞_ and *τ*_*h*_. Only the −120pA TTX trace was used for fitting; the other traces show validation of the model’s parameters using different current clamp steps. *H*_*dist*_=1. Holding current injections: −28 pA for all four steps of −120, −90, −60, −30 pA.

**Figure 4–Figure supplement 3.**
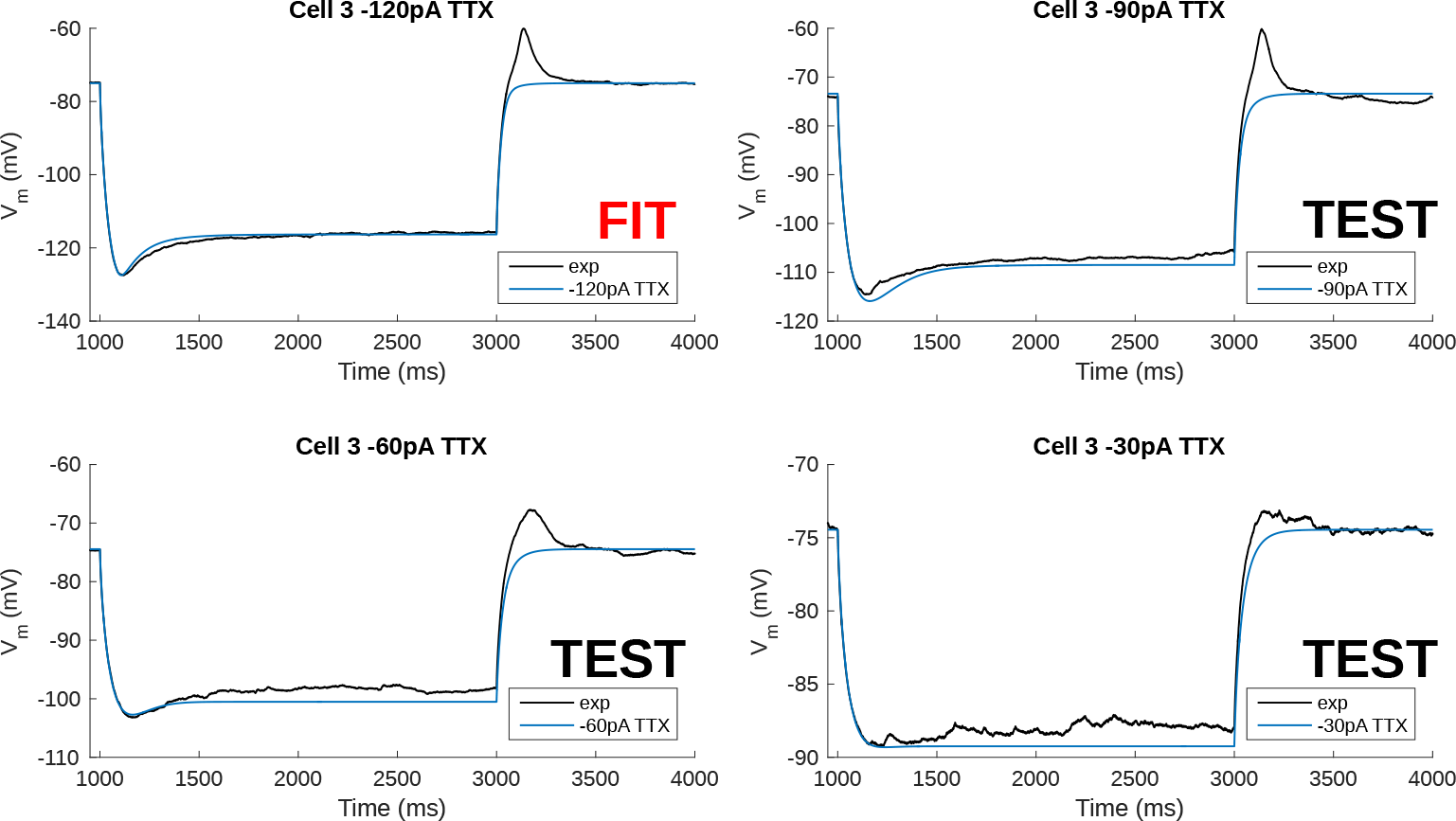
Traces for Cell 2 model compared to experiment after staggered re-fitting. Model *V*_*m*_ traces compared to experiment for *Cell 2* with staggered re-fitting procedure, where first passive properties are fitted, followed by total *G*_*h*_, *r*_∞_ and *r_h_*. Only the −120pA TTX trace was used for fitting; the other traces show validation of the model’s parameters using different current clamp steps. *H*_*dist*_=1. Holding current injections: −5.1 pA for all four steps of −120, −90, −60, −30pA.

**Figure 4–Figure supplement 4.**
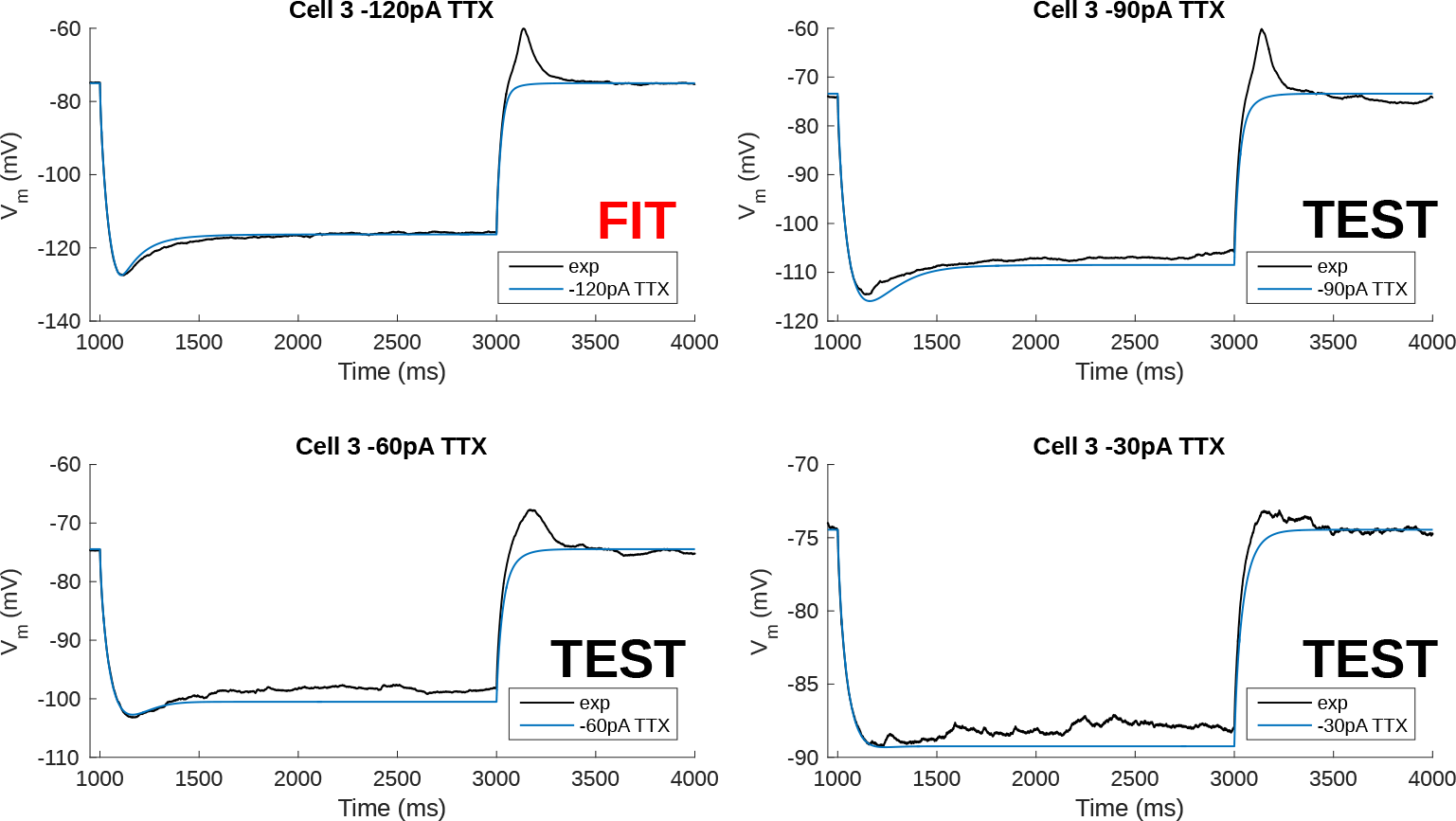
Traces for Cell 3 model compared to experiment after staggered re-fitting. Model *V*_*m*_ traces compared to experiment for *Cell 3* with staggered re-fitting procedure, where first passive properties are fitted, followed by total *G*_*h*_, *r*_∞_ and *τ*_*h*_. Only the −120pA TTX trace was used for fitting; the other traces show validation of the model’s parameters using different current clamp steps. *H*_*dist*_=1. Holding current injections: 2.7 pA for −120pA step; 3.1 pA for −90 and −60pA steps; 3.4 pA for −30pA step.

**Figure 6–Figure supplement 1.**
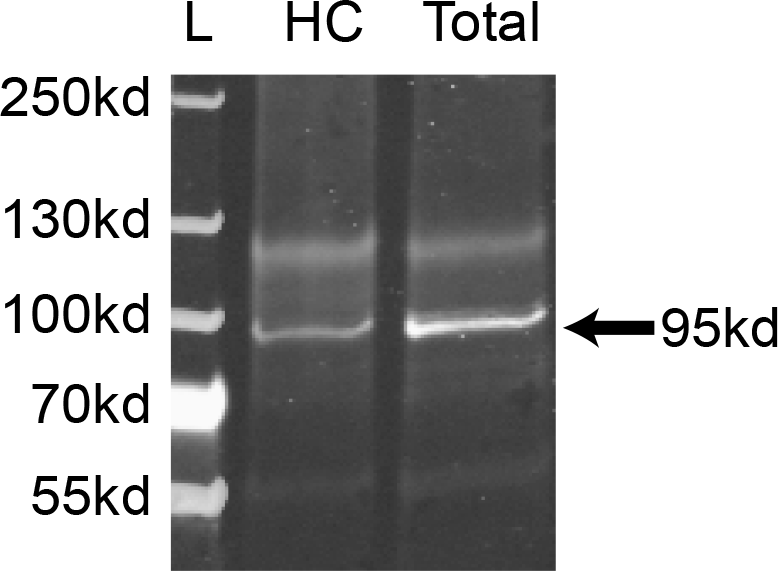
Western blots. A specific band is shown in both hippocampal and total brain lysates, which is consistent with the predicted HCN2 molecular weight (95 kd).

**Figure 6–Figure supplement 2.**
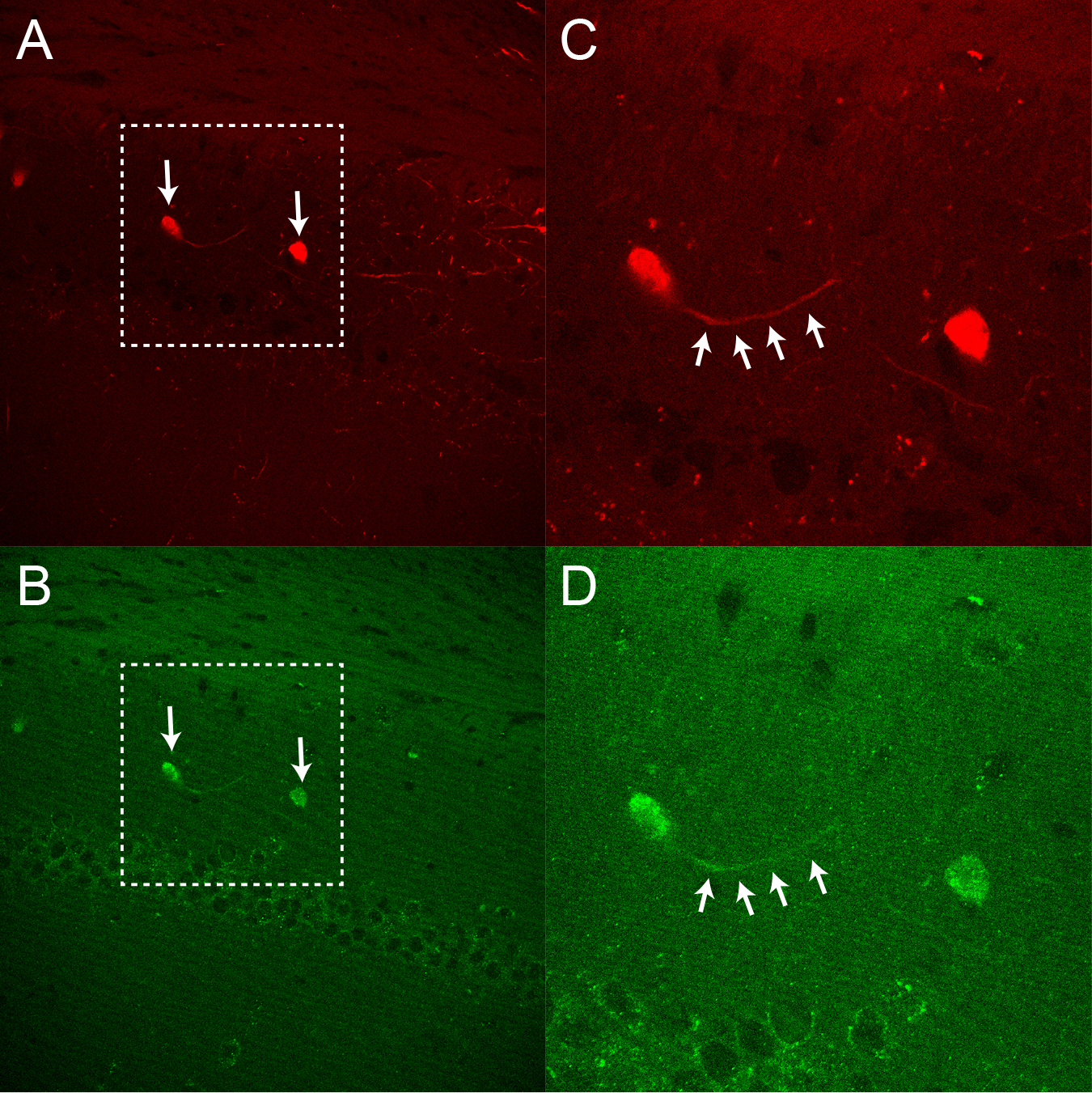
HCN2 in dorsal hippocampal OLM cells. **A.** Endogenous td-Tomato and **B.** HCN2 immunofluorescence imaged in dorsal hippocampal slices from Chrna2-CRE:tdTomato mice. **C., D.** Expanded view of panels **A., B.**. Fewer tdTomato cells were found in dorsal compared to ventral slices.

**Figure 6–Figure supplement 3.**
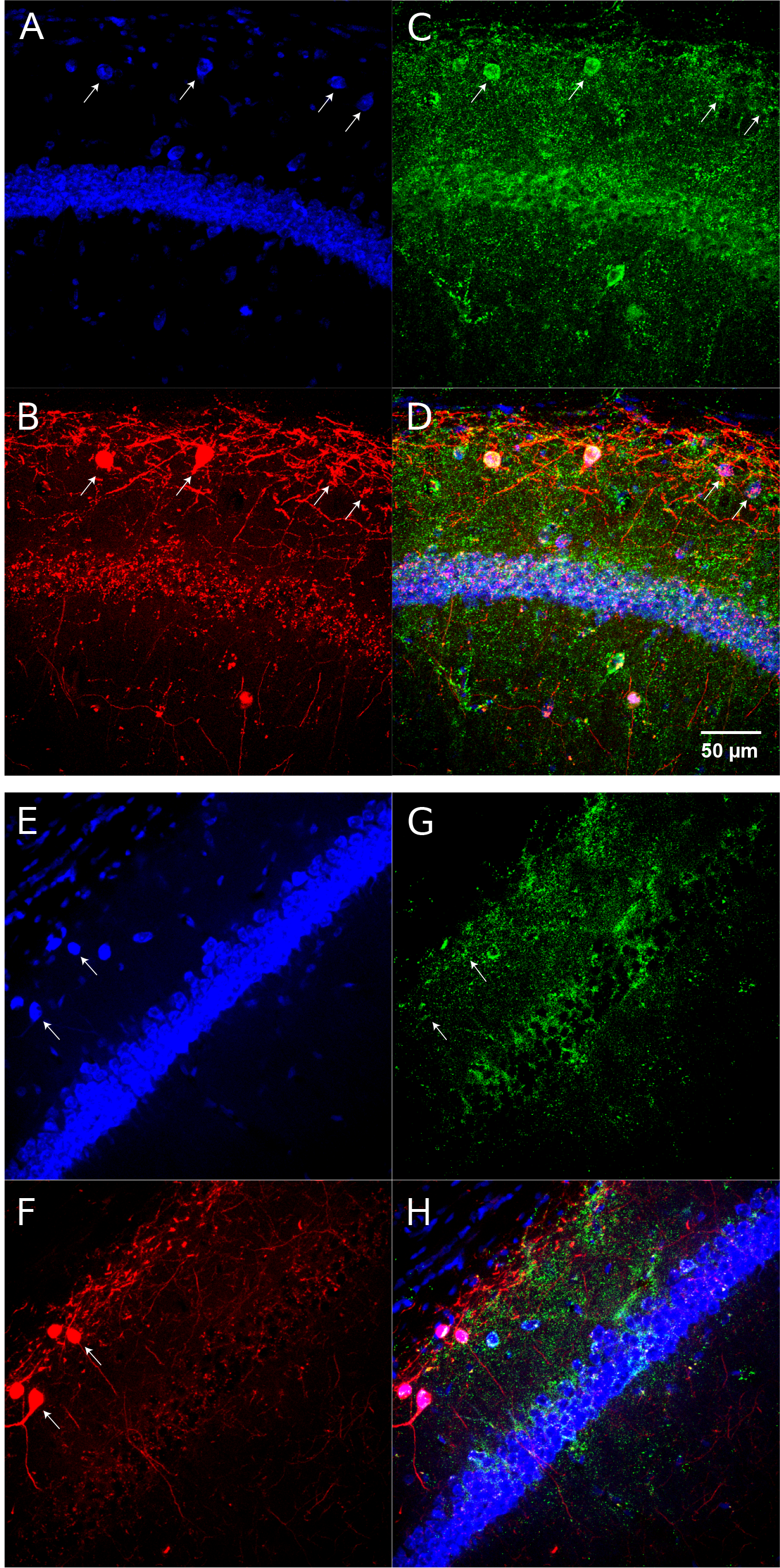
HCN1 immunofluorescence in hippocampal OLM cells. **A.** Neurotrace 435, **B.** Endogenous tdTomato, **C.** HCN1 immunofluroescence, and **D.** Merged view imaged from dorsal hippocampal slices. **E.-G.** as in **A.-D.** HCN1 from ventral hippocampus.

**Figure 7–Figure supplement 1.**
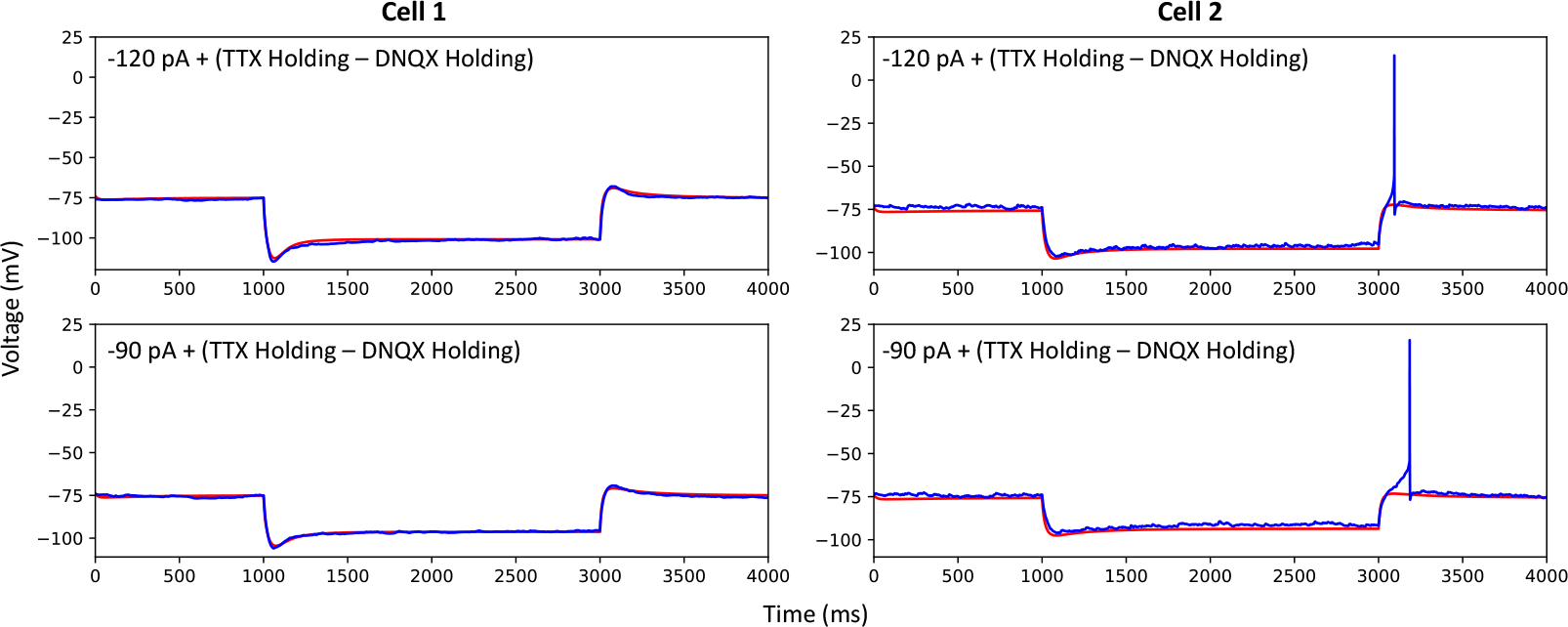
Adding spiking currents does not affect the fit to hyperpolarizing steps. When injecting −90 pA and −120 pA current injections to the top spiking models, we accounted for differences in holding currents during experiment with TTX traces or traces from protocol #2 in **Table 5**-‘DNQX current traces’. The spiking models still generate appropriate hyperpolarization responses. *Cell 1* holding current injections: −28 pA for −90 and −120pA steps, TTX traces, and 4pA for DNQX traces; *Cell 2* holding current injections: −5.1 for −90 and −120pA steps, TTX traces, and −5pA for DNQX traces.

**Figure 7–Figure supplement 2.**
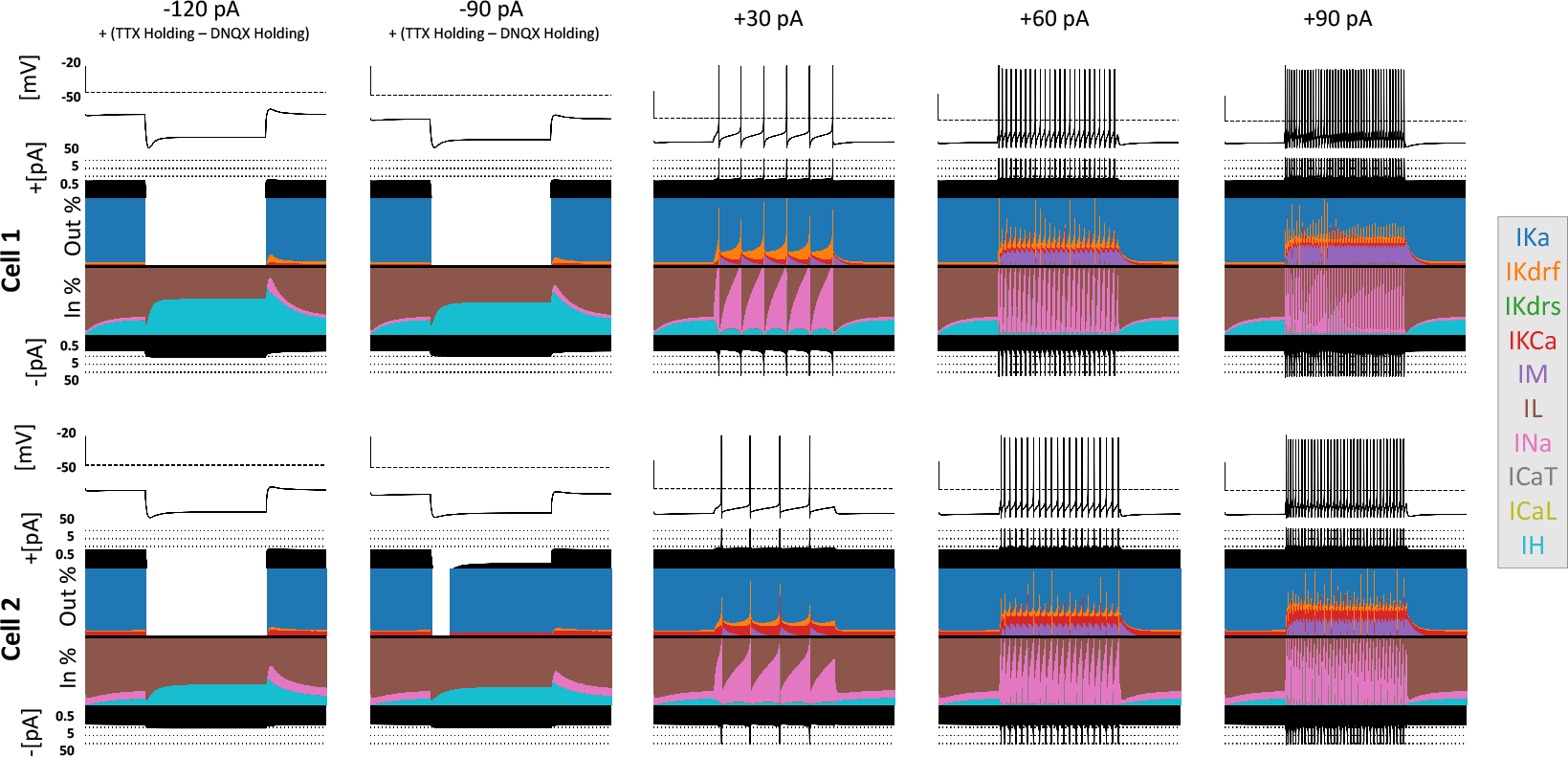
Currentscapes for top spiking models. Using −120 pA, −90 pA, +30 pA, +60 pA, and +90 pA current injection steps, the currentscape plots (*Alonso and Marder, 2019*) indicate the relative current contributions (i.e. the color areas in the plots) of the total inward or outward channel currents (i.e. black areas at the top and bottom of each currentscape plot). Note that the recordings shown here are from the first dendritic compartment adjacent to the soma since calcium channels are not present in the somatic compartment. During hyperpolarizing steps, it is evident from these plots that *I*_*h*_ (IH) and the leak current (IL) are the primary contributors to the electrophysiological output. For depolarizing steps, we see the largest contributions are from A-type potassium current (IKa), fast delayed-rectifier current (IKdrf), sodium current (INa), and calcium-dependent potassium current (IKCa), with increasing contributions from M-type current (IM) as the current step magnitude gets larger. Slow delayed-rectifier current (IKdrs) contributes minimally.

**Figure 7–Figure supplement 3.**
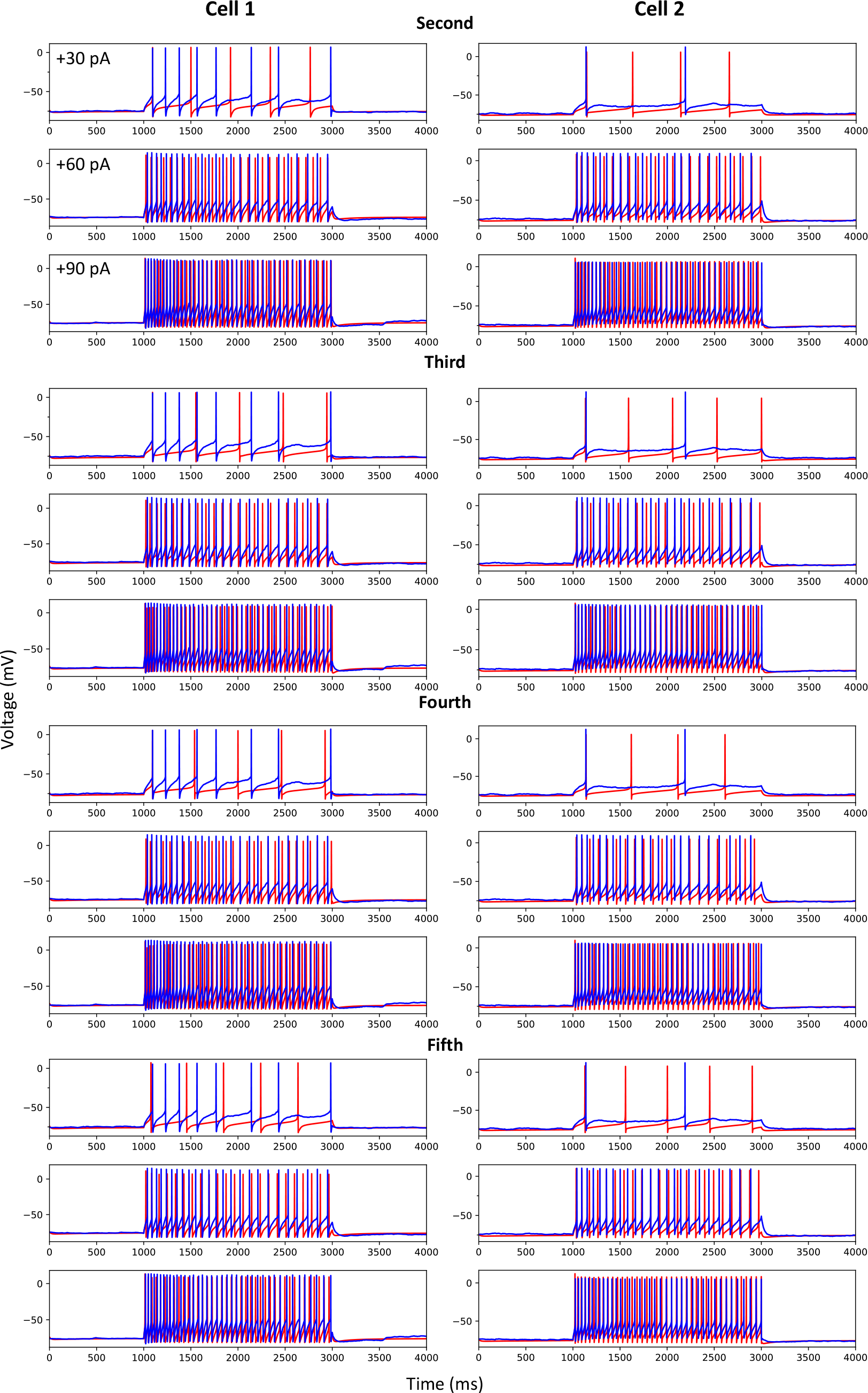
Other top optimized spiking models. As in *Figure 7*, we show the +30 pA, +60 pA, and +90 pA current injection steps for models (red) plotted against the corresponding experimental data (blue). Spiking models for *Cell 1* and *Cell 2* that were ranked second, third, fourth, and fifth are shown from top to bottom.

**Figure 7–Figure supplement 4.**
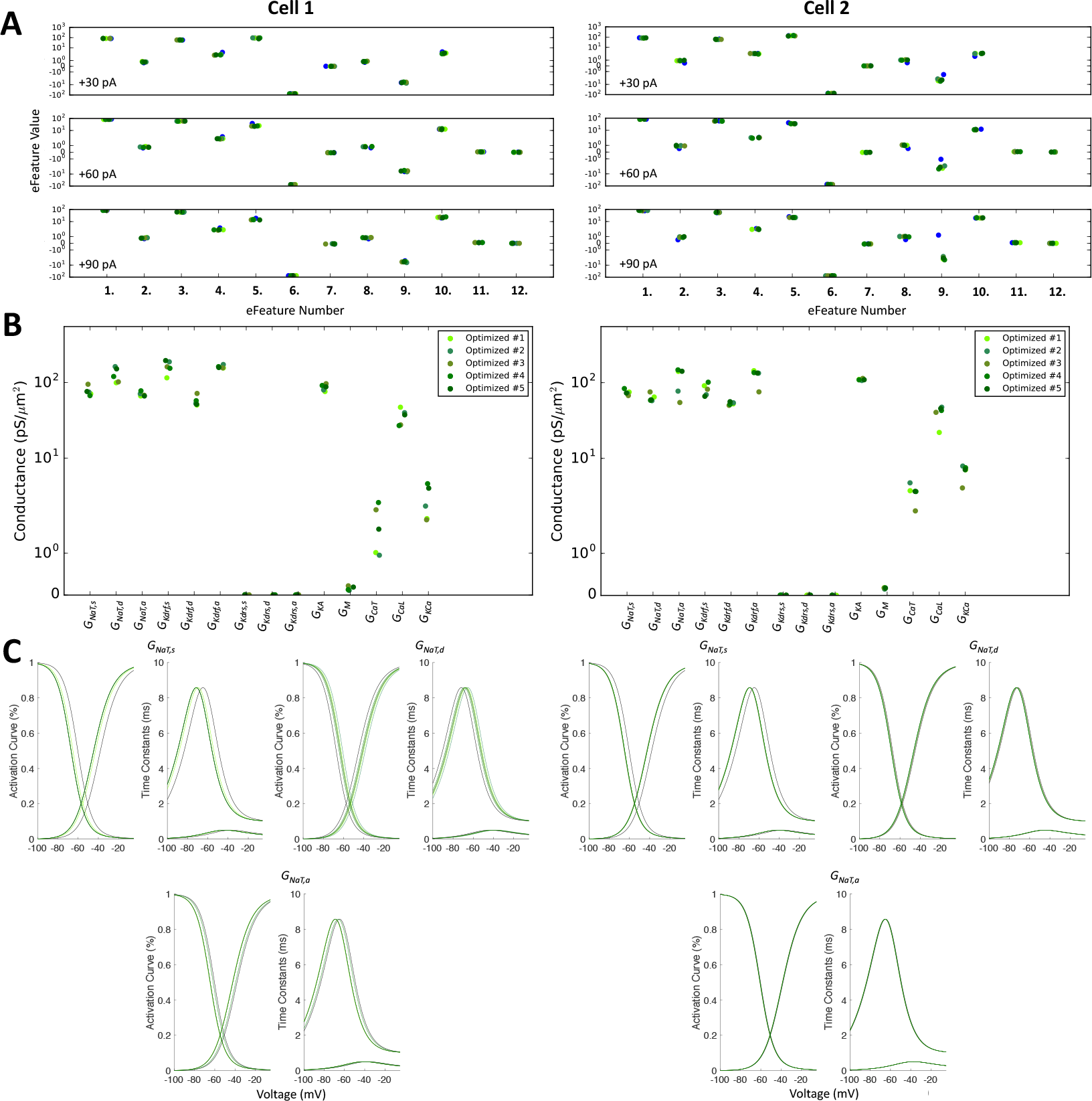
Optimized spiking model features and parameters. **A.** Measurements of objective e-features for each of the top five optimized models (shades of green) during the +30 pA, +60 pA, and +90 pA current injection steps. Each number on the x-axis corresponds to an e-feature. For corresponding e-feature names and descriptions, see *Table 10*. The corresponding target values obtained from the experimental data are shown as blue dots. **B.** Optimized conductance values in the top five spiking models. **C.** Voltage-dependency of somatic, dendritic, and axonal sodium channels were allowed to shift during the optimizations. Here we show the resulting voltage-dependent activation curves and time constants in the top five spiking models (shades of green) as compared to the activation curve used in previous instantiations of the OLM cell model (black curves). See Methods for specific numbers.

## References

Abbas AI, Sundiang MJM, Henoch B, Morton MP, Bolkan SS, Park A J, Harris AZ, Kellendonk C, Gordon JA. Somatostatin Interneurons Facilitate Hippocampal-Prefrontal Synchrony and Prefrontal Spatial Encoding. Neuron. 2018 Nov; 100(4):926–939.e3. doi: 10.1016/j.neuron.2018.09.029.

Abrahamsson T, Cathala L, Matsui K, Shigemoto R, DiGregorio D. Thin Dendrites of Cerebellar Interneurons Confer Sublinear Synaptic Integration and a Gradient of Short-Term Plasticity. Neuron. 2012 Mar; 73(6):1159–1172. doi: 10.1016/j.neuron.2012.01.027.

Almog M, Korngreen A. Is realistic neuronal modeling realistic? Journal of Neurophysiology. 2016 Aug; p. jn.00360.2016. doi: 10.1152/jn.00360.2016.

Alonso LM, Marder E. Visualization of currents in neural models with similar behavior and different conductance densities. eLife. 2019 Jan; 8. doi: 10.7554/eLife.42722.

Angelo K, London M, Christensen SR, Hausser M. Local and Global Effects of Ih Distribution in Dendrites of Mammalian Neurons. Journal of Neuroscience. 2007 Aug; 27(32):8643–8653. doi: 10.1523/JNEUROSCI.5284-06.2007.

Bassett DS, Zurn P, Gold JI. On the nature and use of models in network neuroscience. Nature Reviews Neuroscience. 2018 Sep; 19(9):566–578. doi: 10.1038/s41583-018-0038-8.

Beaulieu-Laroche L, Toloza EHS, van der Goes MS, Lafourcade M, Barnagian D, Williams ZM, Eskandar EN, Frosch MP, Cash SS, Harnett MT. Enhanced Dendritic Compartmentalization in Human Cortical Neurons. Cell. 2018 Oct; 175(3):643–651.e14. doi: 10.1016/j.cell.2018.08.045.

Biel M, Wahl-Schott C, Michalakis S, Zong X. Hyperpolarization-Activated Cation Channels: From Genes to Function. Physiological Reviews. 2009 Jul; 89(3):847–885. doi: 10.1152/physrev.00029.2008.

Bischofberger J, Engel D, Li L, Geiger JR, Jonas P. Patch-clamp recording from mossy fiber terminals in hippocampal slices. Nature Protocols. 2006; 1(4):2075–2081. http://www.nature.com/articles/nprot.2006.312, doi: 10.1038/nprot.2006.312.

Blasco-Ibáñez JM, Freund TF. Synaptic input of horizontal interneurons in stratum oriens of the hippocampal CA1 subfield: structural basis of feed-back activation. Eur J Neurosci. 1995; 7(10):2170–2180.

Borhegyi Z, Varga V, Szilágyi N, Fabo D, Freund TF. Phase segregation of medial septal GABAergic neurons during hippocampal theta activity. The Journal of Neuroscience: The Official Journal of the Society for Neuroscience. 2004 Sep; 24(39):8470–8479. doi: 10.1523/JNEUROSCI.1413-04.2004.

Boyce R, Glasgow SD, Williams S, Adamantidis A. Causal evidence for the role of REM sleep theta rhythm in contextual memory consolidation. Science. 2016 May; 352(6287):812–816. doi: 10.1126/science.aad5252.

Brown HF, Difrancesco D, Noble SJ. How does adrenaline accelerate the heart? Nature. 1979 Jul; 280(5719):235. doi: 10.1038/280235a0.

Bucher D, Prinz AA, Marder E. Animal-to-Animal Variability in Motor Pattern Production in Adults and during Growth. Journal of Neuroscience. 2005 Feb; 25(7):1611–1619. doi: 10.1523/JNEUROSCI.3679-04.2005.

Cardin JA. Inhibitory Interneurons Regulate Temporal Precision and Correlations in Cortical Circuits. Trends in Neurosciences. 2018 Oct; 41(10):689–700. doi: 10.1016/j.tins.2018.07.015.

Cembrowski MS, Spruston N. Heterogeneity within classical cell types is the rule: lessons from hippocampal pyramidal neurons. Nature Reviews Neuroscience. 2019 Feb; p. 1. doi: 10.1038/s41583-019-0125-5.

Cembrowski M, Bachman J, Wang L, Sugino K, Shields B, Spruston N. Spatial Gene-Expression Gradients Underlie Prominent Heterogeneity of CA1 Pyramidal Neurons. Neuron. 2016 Jan; 89(2):351–368. doi: 10.1016/j.neuron.2015.12.013.

Chamberland S, Salesse C, Topolnik D, Topolnik L. Synapse-specific inhibitory control of hippocampal feedback inhibitory circuit. Frontiers in Cellular Neuroscience. 2010; 4:130. doi: 10.3389/fncel.2010.00130.

Chatzikalymniou AP, Skinner FK. Deciphering the Contribution of Oriens-Lacunosum/Moleculare (OLM) Cells to Intrinsic *θ* Rhythms Using Biophysical Local Field Potential (LFP) Models. eNeuro. 2018 Jul; 5(4):ENEURO.0146-18.2018. doi: 10.1523/ENEURO.0146-18.2018.

Chittajallu R, Craig MT, Ashley M, Yuan X, Gerfen S, Tricoire L, Erkkila B, Barron SC, Lopez CM, Liang BJ, Jeffries BW, Pelkey KA, J Mc. Dual origins of functionally distinct O-LM interneurons revealed by differential 5-HT3AR expression.. 2013; 16(11):1598–1607. doi: 10.1038/nn.3538.

Cutsuridis V, Graham B, Cobb S, Vida I. Hippocampal Microcircuits: A Computational Modeler’s Resource Book. 1st edition. ed. Springer; 2010.

Dougherty KA, Nicholson DA, Diaz L, Buss EW, Neuman KM, Chetkovich DM, Johnston D. Differential expression of HCN subunits alters voltage-dependent gating of h-channels in CA1 pyramidal neurons from dorsal and ventral hippocampus. Journal of Neurophysiology. 2013 Apr; 109(7):1940–1953. doi: 10.1152/jn.00010.2013.

Ecker JR, Geschwind DH, Kriegstein AR, Ngai J, Osten P, Polioudakis D, Regev A, Sestan N, Wickersham IR, Zeng H. The BRAIN Initiative Cell Census Consortium: Lessons Learned toward Generating a Comprehensive Brain Cell Atlas. Neuron. 2017 Nov; 96(3):542–557. doi: 10.1016/j.neuron.2017.10.007.

Emmenlauer M, Ronneberger O, Ponti A, Schwarb P, Griffa A, Filippi A, Nitschke R, Driever W, Burkhardt H. XuvTools: free, fast and reliable stitching of large 3D datasets. Journal of Microscopy. 2009 Jan; 233(1):42–60. doi: 10.1111/j.1365-2818.2008.03094.x.

Eyal G, Verhoog MB, Testa-Silva G, Deitcher Y, Lodder JC, Benavides-Piccione R, Morales J, DeFelipe J, Kock CPd, Mansvelder HD, Segev I. Unique membrane properties and enhanced signal processing in human neocortical neurons. eLife. 2016 Oct; 5:e16553. doi: 10.7554/eLife.16553.

Fanselow MS, Dong HW. Are the Dorsal and Ventral Hippocampus Functionally Distinct Structures? Neuron. 2010 Jan; 65(1):7–19. doi: 10.1016/j.neuron.2009.11.031.

Fuhrmann F, Justus D, Sosulina L, Kaneko H, Beutel T, Friedrichs D, Schoch S, Schwarz MK, Fuhrmann M, Remy S. Locomotion, Theta Oscillations, and the Speed-Correlated Firing of Hippocampal Neurons Are Controlled by a Medial Septal Glutamatergic Circuit. Neuron. 2015 Mar; 86(5):1253–1264. doi: 10.1016/j.neuron.2015.05.001.

Gentet LJ, Stuart GJ, Clements JD. Direct Measurement of Specific Membrane Capacitance in Neurons. Biophysical Journal. 2000 Jul; 79(1):314–320. doi: 10.1016/S0006-3495(00)76293-X.

Gloveli T, Dugladze T, Rotstein HG, Traub RD, Monyer H, Heinemann U, Whittington MA, Kopell NJ. Orthogonal arrangement of rhythm-generating microcircuits in the hippocampus. Proceedings of the National Academy of Sciences of the United States of America. 2005 Sep; 102(37):13295–13300. doi: 10.1073/pnas.0506259102.

Goaillard JM, Taylor AL, Schulz DJ, Marder E. Functional consequences of animal-to-animal variation in circuit parameters. Nature Neuroscience. 2009 Nov; 12(11):1424–1430. doi: 10.1038/nn.2404.

Golowasch J, Goldman MS, Abbott LF, Marder E. Failure of Averaging in the Construction of a Conductance-Based Neuron Model. Journal of Neurophysiology. 2002 Feb; 87(2):1129–1131. doi: 10.1152/jn.00412.2001.

Guet-McCreight A, Skinner FK. Using computational models to predict in vivo synaptic inputs to interneuron specific 3 (IS3) cells of CA1 hippocampus that also allow their recruitment during rhythmic states. PLOS ONE. 2019 Jan; 14(1):e0209429. doi: 10.1371/journal.pone.0209429.

Gulyás AI, Görcs TJ, Freund TF. Innervation of different peptide-containing neurons in the hippocampus by GABAergic septal afferents. Neuroscience. 1990; 37(1):31–44.

Harris KD, Hochgerner H, Skene NG, Magno L, Katona L, Gonzales CB, Somogyi P, Kessaris N, Linnarsson S, Hjerling-Leffler J. Classes and continua of hippocampal CA1 inhibitory neurons revealed by single-cell transcriptomics. PLOS Biology. 2018 Jun; 16(6):e2006387. doi: 10.1371/journal.pbio.2006387.

Hay E, Hill S, Schürmann F, Markram H, Segev I. Models of Neocortical Layer 5b Pyramidal Cells Capturing a Wide Range of Dendritic and Perisomatic Active Properties. PLOS Computational Biology. 2011 Jul; 7(7):e1002107. doi: 10.1371/journal.pcbi.1002107.

Hines ML, Carnevale NT. NEURON: a tool for neuroscientists. The Neuroscientist: A Review Journal Bringing Neurobiology, Neurology and Psychiatry. 2001 Apr; 7(2):123–135.

Holmes WR, Ambros-Ingerson J, Grover LM. Fitting experimental data to models that use morphological data from public databases. Journal of Computational Neuroscience. 2006 Jun; 20(3):349–365. doi: 10.1007/s10827-006-7189-8.

Holmes W R. Passive Cable Modeling. In: De Schutter E, editor. Computational Modeling Methods for Neuroscientists Cambridge, MA: MIT Press; 2010.p. 233–258.

Hughes D, Boyle K, Kinnon C, Bilsland C, Quayle J, Callister R, Graham B. HCN4 subunit expression in fast-spiking interneurons of the rat spinal cord and hippocampus. Neuroscience. 2013; 237:7–18. doi: 10.1016/j.neuroscience.2013.01.028.

Ito K, Morita A, Aoki T, Nakajima H, Kobayashi K, Higuchi T. A fingerprint recognition algorithm combining phase-based image matching and feature-based matching. In: International Conference on Biometrics Springer; 2006. p. 316–325. http://link.springer.com/chapter/10.1007/11608288_43.

Jacobs G, Claiborne B, Harris K. Reconstruction of Neuronal Morphology. In: De Schutter E, editor. Computational Modeling Methods for Neuroscientists Cambridge, MA: MIT Press; 2010.p. 187–210.

Jaeger D. Accurate reconstruction of neuronal morphology. In: De Schutter E, editor. Computational Neuroscience: Realistic Modeling for Experimentalists Boca Raton, Fla: CRC Press; 2001.p. 159–178.

Katona L, Lapray D, Viney TJ, Oulhaj A, Borhegyi Z, Micklem BR, Klausberger T, Somogyi P. Sleep and Movement Differentiates Actions of Two Types of Somatostatin-Expressing GABAergic Interneuron in Rat Hippocampus. Neuron. 2014 Apr; doi: 10.1016/j.neuron.2014.04.007.

Kepecs A, Fishell G. Interneuron cell types are fit to function. Nature. 2014 Jan; 505(7483):318–326. http://www.nature.com.myaccess.library.utoronto.ca/nature/journal/v505/n7483/full/nature12983.html?WT.ec_id=NATURE-20140116#outlook, doi: 10.1038/nature12983.

Kispersky TJ, Fernandez FR, Economo MN, White JA. Spike Resonance Properties in Hippocampal O-LM Cells Are Dependent on Refractory Dynamics. Journal of Neuroscience. 2012 Mar; 32(11):3637–3651. doi: 10.1523/JNEUROSCI.1361-11.2012.

Klausberger T. GABAergic interneurons targeting dendrites of pyramidal cells in the CA1 area of the hippocampus. European Journal of Neuroscience. 2009; 30(6):947–957. doi: 10.1111/j.1460-9568.2009.06913.x.

Klausberger T, Magill PJ, Márton LF, Roberts JDB, Cobden PM, Buzsáki G, Somogyi P. Brain-state” and cell-type-specific firing of hippocampal interneurons in vivo. Nature. 2003 Feb; 421(6925):844–848. doi: 10.1038/nature01374.

Klausberger T, Somogyi P. Neuronal Diversity and Temporal Dynamics: The Unity of Hippocampal Circuit Operations. Science. 2008 Jul; 321(5885):53–57. http://www.sciencemag.org/content/321/5885/53.abstract, doi: 10.1126/science.1149381.

Kopell NJ, Gritton HJ, Whittington MA, Kramer MA. Beyond the Connectome: The Dynome. Neuron. 2014 Sep; 83(6):1319–1328. doi: 10.1016/j.neuron.2014.08.016.

Kramis R, Vanderwolf CH, Bland BH. Two types of hippocampal rhythmical slow activity in both the rabbit and the rat: Relations to behavior and effects of atropine, diethyl ether, urethane, and pentobarbital. Experimental Neurology. 1975 Oct; 49(1):58–85. doi: 10.1016/0014-4886(75)90195-8.

Lawrence JJ, Grinspan ZM, Statland JM, McBain CJ. Muscarinic receptor activation tunes mouse stratum oriens interneurones to amplify spike reliability. The Journal of Physiology. 2006 Mar; 571(Pt 3):555–562. http://www.ncbi.nlm.nih.gov/pubmed/16439425, doi: 10.1113/jphysiol.2005.103218.

Lawrence JJ, Saraga F, Churchill JF, Statland JM, Travis KE, Skinner FK, McBain CJ. Somatodendritic Kv7/KCNQ/M channels control interspike interval in hippocampal interneurons. The Journal of Neuroscience: The OZcial Journal of the Society for Neuroscience. 2006 Nov; 26(47):12325–12338. http://www.ncbi.nlm.nih.gov/pubmed/17122058, doi: 10.1523/JNEUROSCI.3521-06.2006.

Lawrence JJ, Statland JM, Grinspan ZM, McBain CJ. Cell type-specific dependence of muscarinic signalling in mouse hippocampal stratum oriens interneurones. The Journal of Physiology. 2006 Feb; 570(Pt 3):595–610. http://www.ncbi.nlm.nih.gov/pubmed/16322052, doi: 10.1113/jphysiol.2005.100875.

Leão RN, Mikulovic S, Leão KE, Munguba H, Gezelius H, Enjin A, Patra K, Eriksson A, Loew LM, Tort ABL, Kullander K. OLM interneurons differentially modulate CA3 and entorhinal inputs to hippocampal CA1 neurons. Nature Neuroscience. 2012; 15(11):1524–1530. doi: 10.1038/nn.3235.

Lörincz A, Notomi T, Tamás G, Shigemoto R, Nusser Z. Polarized and compartment-dependent distribution of HCN1 in pyramidal cell dendrites. . 2002; 5(11):1185–1193. doi: 10.1038/nn962.

Lovett-Barron M, Kaifosh P, Kheirbek MA, Danielson N, Zaremba JD, Reardon TR, Turi GF, Hen R, Zemelman BV, Losonczy A. Dendritic Inhibition in the Hippocampus Supports Fear Learning. Science. 2014 Feb; 343(6173):857–863. doi: 10.1126/science.1247485.

Lovett-Barron M, Losonczy A. Behavioral consequences of GABAergic neuronal diversity. Current Opinion in Neurobiology. 2014 Jun; 26:27–33. doi: 10.1016/j.conb.2013.11.002.

Luo L, Callaway EM, Svoboda K. Genetic Dissection of Neural Circuits: A Decade of Progress. Neuron. 2018 Apr; 98(2):256–281. doi: 10.1016/j.neuron.2018.03.040.

Maccaferri G, McBain CJ. The hyperpolarization-activated current (Ih) and its contribution to pacemaker activity in rat CA1 hippocampal stratum oriens-alveus interneurones. The Journal of Physiology. 1996 Nov; 497 (Pt 1):119–130.

Maccaferri G, David J, Roberts B, Szucs P, Cottingham CA, Somogyi P. Cell surface domain specific postsynaptic currents evoked by identified GABAergic neurones in rat hippocampus in vitro. The Journal of Physiology. 2000 Apr; 524(1):91–116. doi: 10.1111/j.1469-7793.2000.t01-3-00091.x.

Maccaferri G, Lacaille JC. Interneuron Diversity series: Hippocampal interneuron classifications – making things as simple as possible, not simpler. Trends in Neurosciences. 2003 Oct; 26(10):564–571. doi: 10.1016/j.tins.2003.08.002.

Magee JC. Dendritic Hyperpolarization-Activated Currents Modify the Integrative Properties of Hippocampal CA1 Pyramidal Neurons. Journal of Neuroscience. 1998 Oct; 18(19):7613–7624.

Marder E, Goaillard JM. Variability, compensation and homeostasis in neuron and network function. Nature Reviews Neuroscience. 2006 Jul; 7(7):563–574. http://www.ncbi.nlm.nih.gov/pubmed/16791145, doi: 10.1038/nrn1949.

Marder E, Taylor AL. Multiple models to capture the variability in biological neurons and networks. Nat Neurosci. 2011 Feb; 14(2):133–138. http://dx.doi.org.myaccess.library.utoronto.ca/10.1038/nn.2735, doi: 10.1038/nn.2735.

Martina M, Vida I, Jonas P. Distal initiation and active propagation of action potentials in interneuron dendrites. Science (New York, NY). 2000 Jan; 287(5451):295–300.

Marín O. Interneuron dysfunction in psychiatric disorders. Nature Reviews Neuroscience. 2012 Feb; 13(2):107–120. doi: 10.1038/nrn3155.

Matt L, Michalakis S, Hofmann F, Hammelmann V, Ludwig A, Biel M, Kleppisch T. HCN2 channels in local inhibitory interneurons constrain LTP in the hippocampal direct perforant path. Cellular and Molecular Life Sciences. 2011 Jan; 68(1):125–137. doi: 10.1007/s00018-010-0446-z.

Mikulovic S, Restrepo CE, Hilscher MM, Kullander K, Leão RN. Novel markers for OLM interneurons in the hippocampus. Frontiers in Cellular Neuroscience. 2015; 9:201. doi: 10.3389/fncel.2015.00201.

Mikulovic S, Restrepo CE, Siwani S, Bauer P, Pupe S, Tort ABL, Kullander K, Leão RN. Ventral hippocampal OLM cells control type 2 theta oscillations and response to predator odor. Nature Communications. 2018 Sep; 9(1):3638. doi: 10.1038/s41467-018-05907-w.

Molleman A. Patch clamping: an introductory guide to patch clamp electrophysiology. New York: J. Wiley; 2003.

Myatt D, Hadlington T, Ascoli G, Nasuto S. Neuromantic – from Semi-Manual to Semi-Automatic Reconstruction of Neuron Morphology. Frontiers in Neuroinformatics. 2012; 6. doi: 10.3389/fninf.2012.00004.

Müller C, Remy S. Dendritic inhibition mediated by O-LM and bistratified interneurons in the hippocampus. Frontiers in Synaptic Neuroscience. 2014; 6. doi: 10.3389/fnsyn.2014.00023.

Narayanan R, Johnston D. Functional maps within a single neuron. Journal of neurophysiology. 2012 Nov; 108(9):2343–2351. doi: 10.1152/jn.00530.2012.

O’Leary T, Sutton AC, Marder E. Computational models in the age of large datasets. Current Opinion in Neurobiology. 2015 Jun; 32:87–94. http://www.sciencedirect.com/science/article/pii/S095943881500015X, doi: 10.1016/j.conb.2015.01.006.

Rall W, Burke RE, Holmes WR, Jack JJ, Redman SJ, Segev I. Matching dendritic neuron models to experimental data. Physiological Reviews. 1992 Oct; 72(suppl_4):S159–S186. doi: 10.1152/physrev.1992.72.suppl_4.S159.

Ransdell JL, Nair SS, Schulz DJ. Neurons within the Same Network Independently Achieve Conserved Output by Differentially Balancing Variable Conductance Magnitudes. Journal of Neuroscience. 2013 Jun; 33(24):9950–9956. doi: 10.1523/JNEUROSCI.1095-13.2013.

Roth A, Bahl A. Divide et impera: optimizing compartmental models of neurons step by step. The Journal of Physiology. 2009 Apr; 587(Pt 7):1369–1370. doi: 10.1113/jphysiol.2009.170944.

Rotstein HG, Pervouchine DD, Acker CD, Gillies MJ, White JA, Buhl EH, Whittington MA, Kopell N. Slow and Fast Inhibition and an H-Current Interact to Create a Theta Rhythm in a Model of CA1 Interneuron Network. Journal of Neurophysiology. 2005 Aug; 94(2):1509–1518. doi: 10.1152/jn.00957.2004.

Roux L, Buzsáki G. Tasks for inhibitory interneurons in intact brain circuits. Neuropharmacology. 2015 Jan; 88:10–23. doi: 10.1016/j.neuropharm.2014.09.011.

Santoro B, Chen S, Luthi A, Pavlidis P, Shumyatsky GP, Tibbs GR, Siegelbaum SA. Molecular and functional heterogeneity of hyperpolarization-activated pacemaker channels in the mouse CNS. J Neurosci. 2000; 20(14):5264–5275.

Santoro B, Baram TZ. The multiple personalities of h-channels. Trends in Neurosciences. 2003 Oct; 26(10):550–554. doi: 10.1016/j.tins.2003.08.003.

Saraga F, Wu CP, Zhang L, Skinner FK. Active dendrites and spike propagation in multi-compartment models of oriens-lacunosum/moleculare hippocampal interneurons. The Journal of Physiology. 2003 Nov; 552(Pt 3):673–689. http://www.ncbi.nlm.nih.gov/pubmed/12923216, doi: 10.1113/jphysiol.2003.046177.

Schneider CA, Rasband WS, Eliceiri KW. NIH Image to ImageJ: 25 years of image analysis. Nature Methods. 2012 Jul; 9(7):671–675. doi: 10.1038/nmeth.2089.

Schulz DJ, Goaillard JM, Marder E. Variable channel expression in identified single and electrically coupled neurons in different animals. Nature Neuroscience. 2006 Mar; 9(3):356–362. doi: 10.1038/nn1639.

Sekulić V, Chen TC, Lawrence JJ, Skinner FK. Dendritic distributions of Ih channels in experimentally-derived multi-compartment models of oriens-lacunosum/moleculare (O-LM) hippocampal interneurons. Frontiers in Synaptic Neuroscience. 2015; 7:2. http://journal.frontiersin.org.myaccess.library.utoronto.ca/article/10.3389/fnsyn.2015.00002/abstract, doi: 10.3389/fnsyn.2015.00002.

Sekulić V, Lawrence JJ, Skinner FK. Using multi-compartment ensemble modeling as an investigative tool of spatially distributed biophysical balances: application to hippocampal oriens-lacunosum/moleculare (O-LM) cells. PloS One. 2014; 9(10):e106567. doi: 10.1371/journal.pone.0106567.

Sekulić V, Skinner FK. Computational models of O-LM cells are recruited by low or high theta frequency inputs depending on h-channel distributions. eLife. 2017 Mar; 6:e22962. https://elifesciences.org/content/6/e22962v1, doi: 10.7554/eLife.22962.

Sekulić V, Skinner FK. Experiment-Modelling Cycling with Populations of Multi-Compartment Models: Application to Hippocampal Interneurons. In: Cutsuridis V, et al, editors. Hippocampal Microcircuits, Springer Series in Computational Neuroscience Cambridge, MA: Springer; 2018.

Sivagnanam S, Majumdar A, Yoshimoto K, Astakhov V, B A, Martone M, Carnevale NT. Introducing The Neuroscience Gateway, vol. 993 of CEUR Workshop Proceedings of CEUR Workshop Proceedings; 2013.

Siwani S, França ASC, Mikulovic S, Reis A, Hilscher MM, Edwards SJ, Leão RN, Tort ABL, Kullander K.OLM*α*2 Cells Bidirectionally Modulate Learning. Neuron. 2018 Jul; 99(2):404–412.e3. doi: 10.1016/j.neuron.2018.06.022.

Skinner FK, Ferguson KA. Hippocampus, Model Inhibitory Cells. In: Jaeger D, Jung R, editors. Encyclopedia of Computational Neuroscience New York, NY: Springer; 2018.

Soofi W, Archila S, Prinz A. Co-variation of ionic conductances supports phase maintenance in stomatogastric neurons. Journal of Computational Neuroscience. 2012; 33(1):77–95. http://www.springerlink.com/content/t57805006qu12385/abstract/, doi: 10.1007/s10827-011-0375-3.

Stuart GJ, Spruston N. Dendritic integration: 60 years of progress. Nature Neuroscience. 2015 Dec; 18(12):1713–1721. doi: 10.1038/nn.4157.

Swensen AMA, Bean BPB. Robustness of burst firing in dissociated purkinje neurons with acute or long-term reductions in sodium conductance. J Neurosci. 2005; 25(14):3509–3520.

Tang LS, Taylor AL, Rinberg A, Marder E. Robustness of a Rhythmic Circuit to Short” and Long-Term Temperature Changes. The Journal of Neuroscience. 2012 Jul; 32(29):10075–10085. doi: 10.1523/JNEUROSCI.1443-12.2012.

Tennøe S, Halnes G, Einevoll GT. Uncertainpy: A Python Toolbox for Uncertainty Quantification and Sensitivity Analysis in Computational Neuroscience. Frontiers in Neuroinformatics. 2018; 12. doi: 10.3389/fn-inf.2018.00049.

Ulens C, Tytgat J. Functional Heteromerization of HCN1 and HCN2 Pacemaker Channels. J Biol Chem. 2001; 276(9):6069–6072. doi: 10.1074/jbc.C000738200.

Urban-Ciecko J, Barth AL. Somatostatin-expressing neurons in cortical networks. Nature Reviews Neuroscience. 2016 Jul; 17(7):401–409. doi: 10.1038/nrn.2016.53.

Vaidya SP, Johnston D. Temporal synchrony and gamma-to-theta power conversion in the dendrites of CA1 pyramidal neurons. Nature Neuroscience. 2013 Dec; 16(12):1812–1820. doi: 10.1038/nn.3562.

Van Geit W, Gevaert M, Chindemi G, Rössert C, Courcol JD, Muller EB, Schürmann F, Segev I, Markram H. BluePyOpt: Leveraging Open Source Software and Cloud Infrastructure to Optimise Model Parameters in Neuroscience. Frontiers in Neuroinformatics. 2016; 10:17. doi: 10.3389/fninf.2016.00017.

Varga C, Golshani P, Soltesz I. Frequency-invariant temporal ordering of interneuronal discharges during hippocampal oscillations in awake mice. Proceedings of the National Academy of Sciences. 2012 Oct; 109(40):E2726–E2734. doi: 10.1073/pnas.1210929109.

Wilson RI. It takes all kinds to make a brain. Nature Neuroscience. 2010 Oct; 13(10):1158–1160. http://www.nature.com/articles/nn1010-1158, doi: 10.1038/nn1010-1158.

Yi F, Ball J, Stoll KE, Satpute VC, Mitchell SM, Pauli JL, Holloway BB, Johnston AD, Nathanson NM, Deisseroth K, Gerber DJ, Tonegawa S, Lawrence JJ. Direct excitation of parvalbumin positive interneurons by M1 muscarinic acetylcholine receptors: roles in cellular excitability, inhibitory transmission and cognition. The Journal of Physiology. 2014; 592(16):3463–3494. doi: 10.1113/jphysiol.2014.275453.

Zemankovics R, Káli S, Paulsen O, Freund TF, Hájos N. Differences in subthreshold resonance of hippocampal pyramidal cells and interneurons: the role of h-current and passive membrane characteristics: Impedance characteristics and h-current of hippocampal neurons. The Journal of Physiology. 2010 Jun; 588(12):2109–2132. doi: 10.1113/jphysiol.2009.185975.

